# Evolutionary and epidemiological considerations for anthelmintic treatments in ruminant livestock

**DOI:** 10.64898/2026.04.29.721584

**Authors:** Neil Philip Hobbs, John Graham-Brown, Eric R. Morgan, Hannah Rose Vineer

**Author notes:** Department of Vector Biology, Liverpool School of Tropical Medicine, Liverpool, L3 5QA, UK.

## Abstract

Anthelmintic drug resistance is a concern for the sustained control of gastrointestinal nematodes (GINs) in ruminant livestock globally. Evolutionary-epidemiological modelling, which considers both parasite dynamics and resistance dynamics in response to interventions, can be useful in determining which anthelmintic resistance management (ARM) strategies may be effective without compromising parasite control. We address two key questions in ARM. First, how to improve the measurement of AR in populations. Second, identifying effective ARM strategies to slow the spread of AR while maintaining effective parasite control. We developed a simulation framework which tracks the weather-dependent epidemiology of GINs and AR evolution, providing a highly flexible methodology to evaluate multiple ARM strategy options in a single modelling framework, allowing for novel insights due to direct comparisons between strategies. Simulations to refine our understanding of anthelmintic resistance management evaluated the impact of key areas of uncertainty, including transmission intensity, resistance intensity, resistance frequency, drug decay and linking faecal egg count reduction tests (FECRT) to resistance allele frequency. Large-scale simulations present a methodologically thorough evaluation of how treatment choices simultaneously impact epidemiological and evolutionary outcomes. Phenotypic classifications of resistance status using FECRT failed to capture fine scale changes in resistance allele frequency. The pharmacokinetics of drug decay strongly influenced ARM outcomes, and trade-offs between ARM and effective parasite control depends on genetic factors underpinning resistance. Combination therapies appear to be the most effective resistance management strategy evaluated. Our findings suggest practical implementations to manage anthelmintic resistance must simultaneously consider parasite transmission, pharmacology and parasite genetics to be robust and sustainable. We provide a rigorous simulation framework to enable such discussions allowing for a refinement into our understanding of parasite control in the presence of resistance evolution.

## 1. Introduction

Chemical control has been widely used to target pathogens, pests and vectors worldwide for decades, leading to the inevitable evolution of resistance [2, 3, 4]. Today, resistance to treatments are some of the most pervasive threats, undermining the sustainability of vector and parasite control in livestock management practices (Charlier et al., 2020; Githaka et al., 2022; Rose Vineer et al., 2020). Resistance management is set of practices (Roush, 1989) intended to delay, slow or mitigate the emergence and spread of resistance are widely recommended across disease systems (Stubbings et al., 2020; WHO, 2012); and their evaluation is rapidly becoming one of the most urgent challenges in livestock veterinary research.

Mathematical modelling has played a key guiding role in the design and evaluation of resistance management strategies (Rose Vineer, 2020; Tabashnik, 1990). While resistance management strategies used in different systems often share cross-cutting themes, the biological, legislative, operational and economic differences necessitate system-specific models. We focus on anthelmintic resistance (AR) and AR management (ARM) in gastrointestinal nematodes (GINs) of ruminant livestock. AR is estimated to cost the ruminant livestock industry €38 million annually in Europe (Charlier et al., 2020), a cost that is likely to increase as AR increases. Despite ARM modelling being conducted since the 1980s (Gettinby et al., 1989), there remains uncertainty around the conditions impacting ARM strategy success or failure. Many resistance management claims (REX Consortium, 2013) were neither investigated theoretically nor empirically. For ARM there are two areas requiring attention. First, how to improve the measurement of AR in populations, and how this is subsequently used to inform strategy choice. Second, identifying effective ARM strategies to slow the spread of AR while maintaining effective parasite control.

The faecal egg count reduction test (FECRT) is a practical, pragmatic measure used to measure anthelmintic treatment efficacy of GINs (Coles et al., 1992). FECRT results can be used to classify GIN populations as “resistant” or “susceptible” using heuristic thresholds from World Association for the Advancement of Veterinary Parasitology (WAAVP) guidelines (Kaplan, 2023). Recent discussions around FECRTs have focused on the statistical requirements for measurements to inform treatment practices (Denwood et al., 2023), but have not considered the genetics of resistance. Incorrect classification of resistance in populations may lead to incorrect ARM practices.

Transmission intensity, AR genetics and drug pharmacology may concurrently impact parasite control and AR dynamics (Coffeng et al., 2024; Masserey et al., 2022), and it is essential to understand these factors. Areas where GIN transmission is high (due to climate or previous farm management practices) likely have high contamination levels, and therefore a large free-living population in *refugia*, which may itself interact with ARM (van Wyk, 2001) and infection. Models commonly assume a dominance of 0.5 (e.g.,(Patel et al., 2024)), such that the RS phenotype (heterozygote) is intermediary to the SS (homozygous susceptible) and RR (homozygous resistant) phenotypes. Dominance has important interactions with the decay in treatment efficacy (Hastings, 2001), which may be overlooked as a result. The reduction in the efficacy due to decay has been shown to be an important consideration for both monogenic (South et al., 2020) and polygenic (Hobbs and Hastings, 2025a) resistance. Despite this, the decay in treatment efficacy over time is often a neglected parameter in ARM models. Considering drug decay is most important for treatments with long residual efficacies, such as “newer” anthelminthic formulations (Papadopoulos et al., 2009).

There is a lack of communication between resistance modelling disciplines (Hastings, 2001; Peck, 2001; REX Consortium, 2007) and inevitably, diverse terminologies are used, necessitating individual studies to clarify their terminologies. Unclear and inconsistent terminologies lead to misunderstanding and misinterpretation. To clarify, we consider ARM strategies as the overarching way in which anthelmintic drugs are deployed over space and time. Other aspects such as “*refugia*” or “rotational grazing” are supplementary considerations which may further influence ARM strategy performance. We will discuss the strategies which can generally be placed under the umbrella terms of pharmacological rotations, cohort mosaics and combinations, which we clarify below.

“Rotations” refer to the process of deliberate, pre-planned switches between active ingredients. Synonymously called cycling or alternation, which are also used *ad hoc* (REX Consortium, 2013). For farm livestock management, the term rotation can also be used to describe grazing practices, such as rotational grazing, which is itself a GIN control option (Stromberg and Averbeck, 1999). To distinguish, we use “pharmacological rotations” to describe rotating between different anthelmintic classes. Rotations are suggested as a strategy because resistance to one active can decline while rotated out (Madgwick and Kanitz, 2024).

“Combinations” classically refer to single formulations containing two different active ingredients, which both drugs are effective against the same target species (“narrow spectrum”). The nearest equivalent strategies would be pyramiding for transgenic crops and mixtures for fungicides, herbicides and insecticides. The resistance management rationale of combinations is redundant kill (Curtis, 1985), whereby the dual exposure to two different active ingredients means that even if an individual is resistant one, they are still likely to be killed by the other partner active ingredient (Madgwick and Kanitz, 2023). For anthelmintics combinations can also refer to “broad spectrum” products, where each active ingredient may be effective against different parasite species.

“Mosaics” refer to the deployment of different active ingredients over space. albeit the spatial scale is not usually further clarified, with spatial scale varying between biological/operational systems. For example, classically in agriculture, mosaics refer to different fields being treated with different pesticides. While for mosquito control, mosaic generally refers to different villages. The term “micro-mosaic” has been used to describe different households in the same village receiving different treatments (Hobbs and Hastings, 2025b). For drug treatments, this could also be at the farm/hospital scale, or at the individual patient level, as each patient acts as its own spatial unit. One option for livestock would be to treat different animals within the same herd with different active ingredients, which we term as a “cohort mosaic”.

“*Refugia*” is a common consideration in resistance modelling; including even the very earliest resistance models (Comins, 1977). To put simply, *refugia* is the subset of the population of the target organism which remains untreated. The use of *refugia* is a considered an anthelmintic resistance management (ARM) strategy (Hodgkinson et al., 2019), but is better considered as an additional component. The term *refugia* in the anthelmintic literature can conflate between “free-living larval *refugia*” and “untreated livestock *refugia*” (Hodgkinson et al., 2019), which both meet the technical definition of *refugia*. *Refugia* in the anthelmintic resistance literature tends to refer to the “free-living larval *refugia*” (van Wyk, 2001). These distinctions are important considering the ability to measure and quantify the size of each type of *refugia,* and how different types of *refugia* may impact ARM and disease control.

The aim of this manuscript is to provide improved clarity and conceptual guidance for the discussion of ARM by farmers, veterinarians, researchers and policy makers, by extension of the GLOWORM modelling framework (Ingle et al., 2025; Rose et al., 2015; Rose Vineer et al., 2020) to evaluate the implications of various ARM options.

## 2. Methods

### 2.1 Model Overview

We extend the GLOWORM simulation framework, which in brief, is a GIN transmission model which captures the lifecycle of GIN infecting ruminant livestock and encompasses the free-living (Rose et al., 2015) and parasitic (Rose Vineer et al., 2020) life stages of the GINs. The model has been further extended to simulate parasite metapopulations, comprising sub-populations on separate patches (fields within a farm) (Ingle et al., 2025), the META extension. We further extend this by including livestock metapopulations (COHORT extension) and resistance genetics, as either a single-gene (AR extension) or two-gene (AR2 extension). The technical detail is fully described in the supplementary file (Section S1), but given as a conceptual overview here. The conceptual modularisation and extension of the GLOWORM framework is presented in Figure 1. Full model code and simulation output are provided in an Open Science Framework repository (Hobbs and Rose Vineer, 2026).

**Figure 1.**
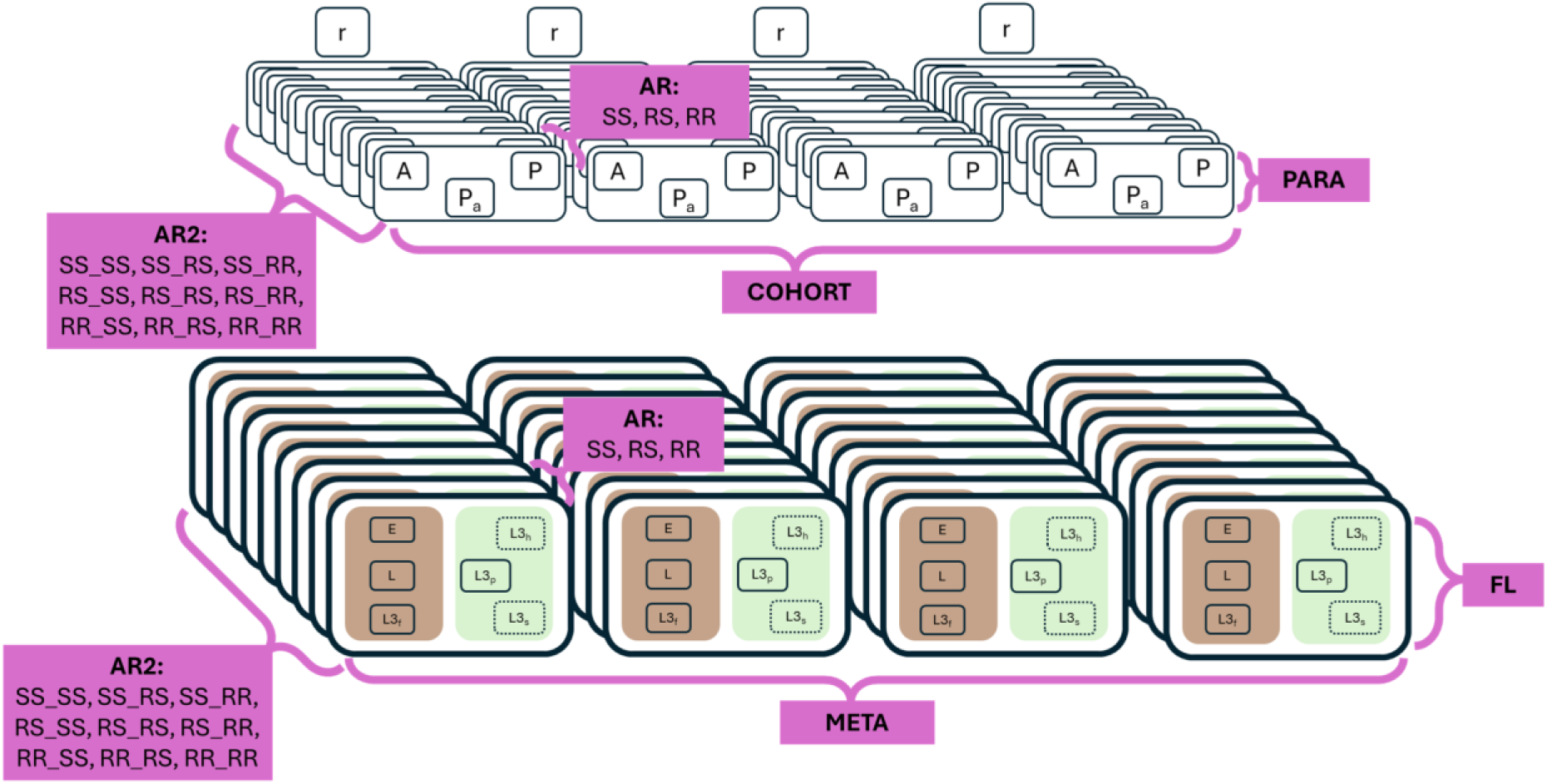
GLOWORM-META-COHORT-{AR, AR2} Modularisation. The GLOWORM model framework consists of a modular format, allowing for the different extensions to be used in unique permutations. The FL (lower half of diagram) (Rose et al., 2015) and PARA (upper half of diagram) (Rose Vineer et al., 2020) are the core parasite transmission components of the model, used across all other model extensions. The META (Ingle et al., 2025) component extends to allow for separate patches for free-living parasites (represented by replication from left to right of the lower half of the diagram). The new COHORT extension allows for separate groups of grazing livestock (represented by replication from left to right in the upper half of the diagram). The AR and AR2 extensions are when the number of different anthelmintic genes being tracked is specified (represented by replication from front to back of the diagram, on the z axis).

The COHORT extension duplicates the in-host equations, so that each cohort has its own set of equations, which links pasture subpopulations by ingestion of the L3 (infection) and excretion of eggs in faeces (contamination). Each cohort can be given unique properties such as age, patch movements or treatments. For the AR and AR2 extensions, each genotype permutation is tracked individually for each life stage of the GIN, in each patch and cohort. This allows for treatments to have unique impacts against each genotype. The assumption of monogenic resistance is widespread throughout the resistance modelling literature regardless of biological system (e.g., (Helps et al., 2020; Levick et al., 2017; Madgwick and Kanitz, 2022; Reichert et al., 2023; REX Consortium, 2013), and is consistent with single nucleotide polymorphisms conferring resistance to benzimidazoles and levamisole (Doyle et al., 2022). This approach can also represent more complex multiple co-selected and linked genomic regions conferring resistance, as may be the case for ivermectin resistance, for example (McIntyre et al., 2025).

The addition of the COHORT and AR/AR2 extensions to the GLOWORM framework, allows for an increase in the number, variety and complexity of ARM strategies that can be investigated, some demonstrations of the updated model capability are in the supplementary file (see Section S4). Model validation is conducted against previously published data (Sargison et al., 2012; Stear et al., 2006; Waller and Thomas, 1975), and is detailed in the Supplementary File (Section S5).

### 2.2 Simulation sets

We consider three simulation sets, which each have a different focus.

- Simulation Set 1: Resistance classification and linking FECRT to resistance frequency and treatment efficacy
- Simulation Set 2: Impact of genetic and drug efficacy properties on parasite control and resistance spread.
- Simulation Set 3: Evaluating strategies for parasite control and managing resistance.

The GLOWORM simulation framework has a large degree of capability and flexibility (Figure 1); running these sets separately allows for the specific aspects to be investigated in isolation, providing better mechanistic understanding than if all features were investigated simultaneously. For all our presented simulations we use the GLOWORM model with the *Haemonchus contortus* parameterisation (Rose et al., 2015), this is fully detailed in the model code in the “*gin*” function. As we are using the model and reporting the results in highly generalisable ways, the high-level results (e.g., directionality) would be broadly applicable to any GIN species in any livestock species. Given drug formulations vary in efficacy depending on the livestock breed (Gotardo et al., 2020; Hosking et al., 2010), parasite species, and exact formulation/method of application (Garg et al., 2007), we are also agnostic with regards to the specific anthelmintic deployed, but explore different characteristics which are detailed later. The generalisable performance of different ARM options to understand the evolutionary and epidemiological implications has the added benefit of results being broadly applicable to potential “new” anthelmintics for which pharmacokinetics, spectrum of activity and genetic mechanisms of resistance are unknown. In all simulations, resistance allele frequencies in the adult GIN population are initiated in Hardy-Weinberg equilibrium (*R^2^ + 2RS + S^2^ = 1*) for both within-host and free-living stages.

We consider livestock (sheep) reared at a stocking rate of ten per hectare (equivalent to 1 livestock unit (Eurostat, n.d.)). The livestock remain at a fixed weight of 65kg. We consider only single-patch simulations (not using the META extension), where sheep remain on the same pasture throughout the simulations (there is no rotational grazing), because this ensures the impact of the cohort *refugia* and larval *refugia* can be better isolated and investigated explicitly. Location-specific precipitation and temperature were extracted with the *openmeteo* R package (Pisel, 2023).

#### 2.2.1 Simulation Set 1 Design: Resistance classification and linking FECRT to resistance frequency and treatment efficacy

Simulation Set 1 considers how observed drug efficacy, from FECRTs, translates to the GIN resistance allele frequency. FECRT is calculated given the input resistance allele frequency, the genotype, drug parasite kill rate and dominance. Simulations only need to be of a short duration (10-days) to cover the treatment period, and the impact of climate can be ignored. Simulations are run for a single anthelmintic, to align with current WAAVP guidelines (Kaplan, 2023) which do not consider combination therapies.

We vary the following parameters:

- *Drug efficacy*: This is the parasite kill rate against the SS genotype. We consider the drug efficacy values of 1, 0.999, 0.99, 0.98, 0.95, 0.9. Where a value of 1 indicates all SS genotypes are killed.
- *Resistance intensity*: This is the parasite kill rate against the RR genotype. We consider resistance intensity values of 0, 0.001, 0.01, 0.02, 0.05, 0.1. Where a value of 0 indicates no RR genotypes are killed.
- *Dominance*: We consider the dominance of resistance, which describes the RS phenotype in relation to the SS and RR phenotypes. Where a dominance of 0 indicates the RS phenotype is the same as the SS phenotype, and a dominance of 1 indicates the RS phenotype is the same as the RR phenotype. We explore all values between 0 and 1 at 0.05 intervals.
- *Resistance frequency*: This is the frequency of the R allele in the population. We explore all values between 0 and 1 at 0.01 intervals.

To clarify, resistance intensity describes the level of resistance the RR genotype provides, while resistance frequency is the occurrence of the resistance allele in the population. GIN populations could have frequent but low intensity resistance or, conversely, infrequent but intense resistance.

The parameter space described above gives a total of 76,356 unique parameter permutations. For simplicity, we assume the drug has an instantaneous effect on reducing the adult worm burden by the specified level given the genotype, and that there is no drug decay. The outcome extracted from each simulation permutation is the FEC on day 1 (pre-treatment) and day 10 (post-treatment), which is used to calculate the FECRT. The exact FECRT is reported for each simulation permutation. We further classify simulated populations as “resistant” or “susceptible”, considering various thresholds which are or have been used or proposed: 95%, 90% and 80% (Coles et al., 1992; Kaplan, 2023), to demonstrate how changing threshold impacts the classification of population.

#### 2.2.2 Methods common to simulation sets 2 and 3

Simulation sets 2 and 3 concern the deployment of anthelmintic drugs over multiple years. Each simulation is three years in duration, starting 01/01/2018 and ending 31/12/2020 from 100 randomly sampled locations in the UK to account for weather variability. Simulations start with a 1-year “burn-in” to establish the desired conditions for the subsequent years (i.e., suitable pasture contamination levels and immunity). No anthelmintic treatments are given during the burn-in period, and the burn-in is not included in the analysis. The burn-in generates two generalised transmission intensity settings of “lower” and “higher” for each location. Here, transmission intensity corresponds to the level of pasture contamination that would on average (across all locations) give a peak FEC of 1250 to 1500 (“higher” transmission) and 750 to 1000 (“lower” transmission) in the second year in the absence of drug treatments. Transmission intensity calibration details are presented in the supplementary file (Section S2, Figure S2.1).

**Figure 2.**
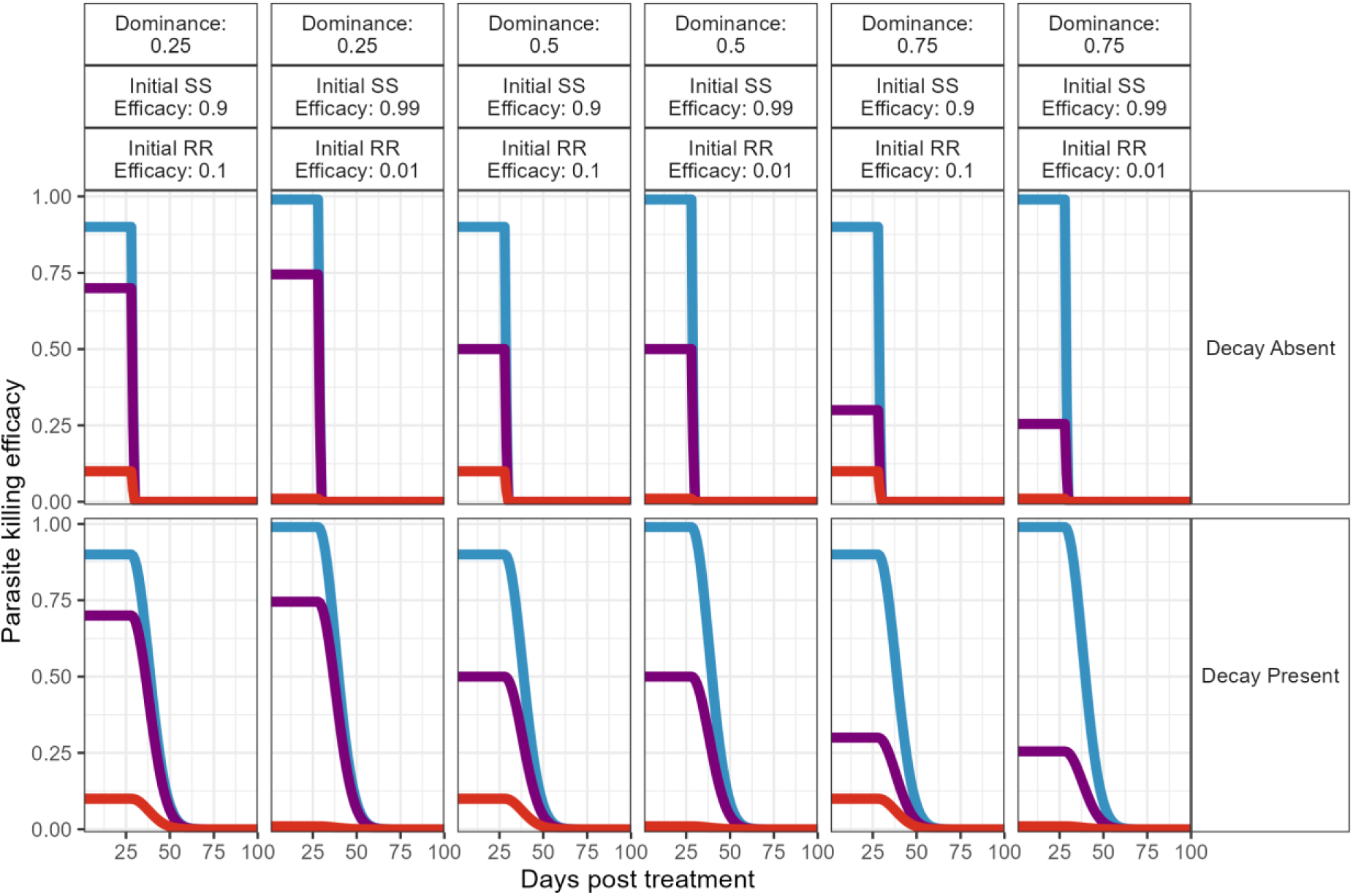
Anthelminthic drug decay profiles used in the simulations. Plot shows the drug efficacy (parasite killing) for various drug decay profiles and resistance genetic situations. In each plot, the line colours are the parasite killing efficacy over time for the SS (blue), RS (purple) and RR (red) genotypes. The panel descriptions to the right indicate how drug decay is implemented. The top panels describe the parasite genetics and resistance status, such that the top row is dominance, middle is the initial (at treatment) efficacy to the SS genotype, and the bottom row descriptor is the initial (at treatment) efficacy to the RR genotype. The y axis describes the time since treatment, that is the number of days since the drug treatment was given, where day 1 is the day treatment is given.

From the simulations we extract the following outcomes:

a. the FEC in each cohort and the whole population as a measure of parasitism,
b. L3 (infectious larval stage) pasture count as a measure of transmission intensity,
c. resistance allele frequency in the L3 on pasture, as a measure of ARM.

#### 2.2.3 Simulation Set 2 Design: Impact of genetic and drug efficacy properties on parasite control and resistance spread

Simulation Set 2 explores the implication notable parameters on the spread of drug resistance and their interaction with parasite control. These parameters are frequently highlighted as important considerations (REX Consortium, 2013), albeit under-investigated in the veterinary GIN context. We consider treatments with a single active ingredient to focus on the mechanistic understanding in relation to these parameters. Anthelmintic treatments are given on day 100 of years 2 and 3 of the simulation to the livestock in Cohort 1 (the treatment cohort) irrespective of the worm burden. Cohort 2 (the *refugia* cohort) never receives any drug treatment. The anthelmintic drug targets only the adult life stage of the GIN. All possible permutations were used (288 permutations) and replicated across the 100 locations (28,800 simulations). The parameters explored were:

- *Refugia size*: “Untreated livestock *refugia*”, such as achieved through targeted selective treatment (TST) (Gaba et al., 2010); is implemented by dividing the sheep into two cohorts which co-graze the same pasture and differ only in the anthelmintic treatments given. The stocking rate of cohort 1 (“treatment cohort”) and cohort 2 (“*refugia* cohort”) therefore sums to ten. We investigate TST by withholding the drug treatments from 0, 10 or 20% of the livestock population, which describes the relative sizes of the treatment and *refugia* (untreated) cohorts. Only Cohort 1 (“treatment cohort”) receives anthelmintic treatments.
- *Transmission intensity:* higher and lower as previously detailed.
- *Drug decay*: We consider an anthelmintic treatment of a single active ingredient with the general properties of having an effective duration of four weeks (28 days) after which drug decay either is present or absent. Details of the inclusion are in section S2 of the Supplementary file. A visual representation is in Figure 2.
- *Resistance frequency (allele frequency):* The starting resistance allele frequency is 0.01 (1%) or 0.1 (10%) in the simulations.
- *Drug Efficacy*: parasite killing efficacy against the SS genotype of 99 or 90%
- *Resistance Intensity*: The initial parasite killing efficacy against the RR genotype of 1 or 10%.
- *Dominance*: The initial parasite killing efficacy against the RS genotype depends on the RR and SS phenotypes and dominance, which was investigated considering dominance to be 0.25, 0.5 or 0.75.

#### 2.2.4 Simulation Set 3 Design: Evaluating strategies for parasite control and managing resistance

The purpose of simulation set 3 is to consider the deployment of different active ingredients (e.g., drug classes) as part of rotational, mosaic or combination ARM strategies over space and time, and evaluate their comparative performances/effectiveness (evolutionary and epidemiological), to identify broad recommendations of how to better deploy anthelmintics. We consider the deployment of two anthelmintic drugs (A and B), where resistance to each drug is encoded by a single gene. The anthelmintics differ in their mode of action, with no cross resistance or cross selection. Treatments are given monthly from February 1^st^ to July 1^st^ (six treatments) in years two and three. Each drug treatment has a residual duration of one week with no efficacy decay. Each round of treatment covers 80% of the livestock population. We evaluate a total of nine different ARM strategies. The parasite kill rates against each genotype to each drug were: 0.95 (RR), 0.5 (RS) and 0.05 (SS), dominance is 0.5. We consider only the “higher” transmission parameterisation previously detailed.

Further to the broad definitions given in the introduction, rotations, mosaics and combinations are interpreted for ARM simulations as follows:

*Pharmacological rotations:* The frequent rotation of active ingredients has often been recommended as a generalisable resistance management strategy (WHO, 2012) to prevent reliance on a single active ingredient. Although ARM guidance is shifting away from this generalisable recommendation to more farm-specific recommendations (Stubbings et al., 2020), in the absence of farm-level data, rotations may be defaulted to for ARM.

*Mosaics:* Mosaic strategies involve deploying different interventions in different locations. The mosaic term is agnostic to spatial scale, which may have different impacts. For pesticide/herbicide applications mosaics could be at the farm, field or row scale. For anthelmintics in livestock we can consider the “landscape” to be the field the livestock animals are grazing on, or when considering landscape from the GIN perspective each individual livestock animal is itself a component of the mosaic. We therefore distinguish these as “paddock mosaics” and “cohort mosaics”. We do not investigate “paddock mosaics” further. For “cohort mosaics” we can further consider how treatments are given to the cohorts over time. In all scenarios, both cohorts co-graze the same patch (pasture). We acknowledge cohort mosaics are likely challenging to implement logistically.

*Combinations:* both drugs (A and B) are administered simultaneously as a single drug formulation, where each drug is given at their recommended therapeutic dose. We assume (as is a common assumption [e.g., 33–36]) that survival is multiplicative with drug combinations. For example, the RR_RR survival probability would be 0.95 (drug A) * 0.95 (drug B), which is 0.9025. And for the SS_SS genotype would be 0.05 (drug A) * 0.05 (drug B), which is 0.0025 (see Figure S3.1 in Section 3 of the Supplementary File). The parasite kill rate is therefore one minus the survival probability. To account for the expected higher financial costs of combinations (as each treatment contains two active ingredients), we also consider two potential compromises (assuming drug dose is not compromised) regarding treatment coverage and frequency.

**Figure 3:**
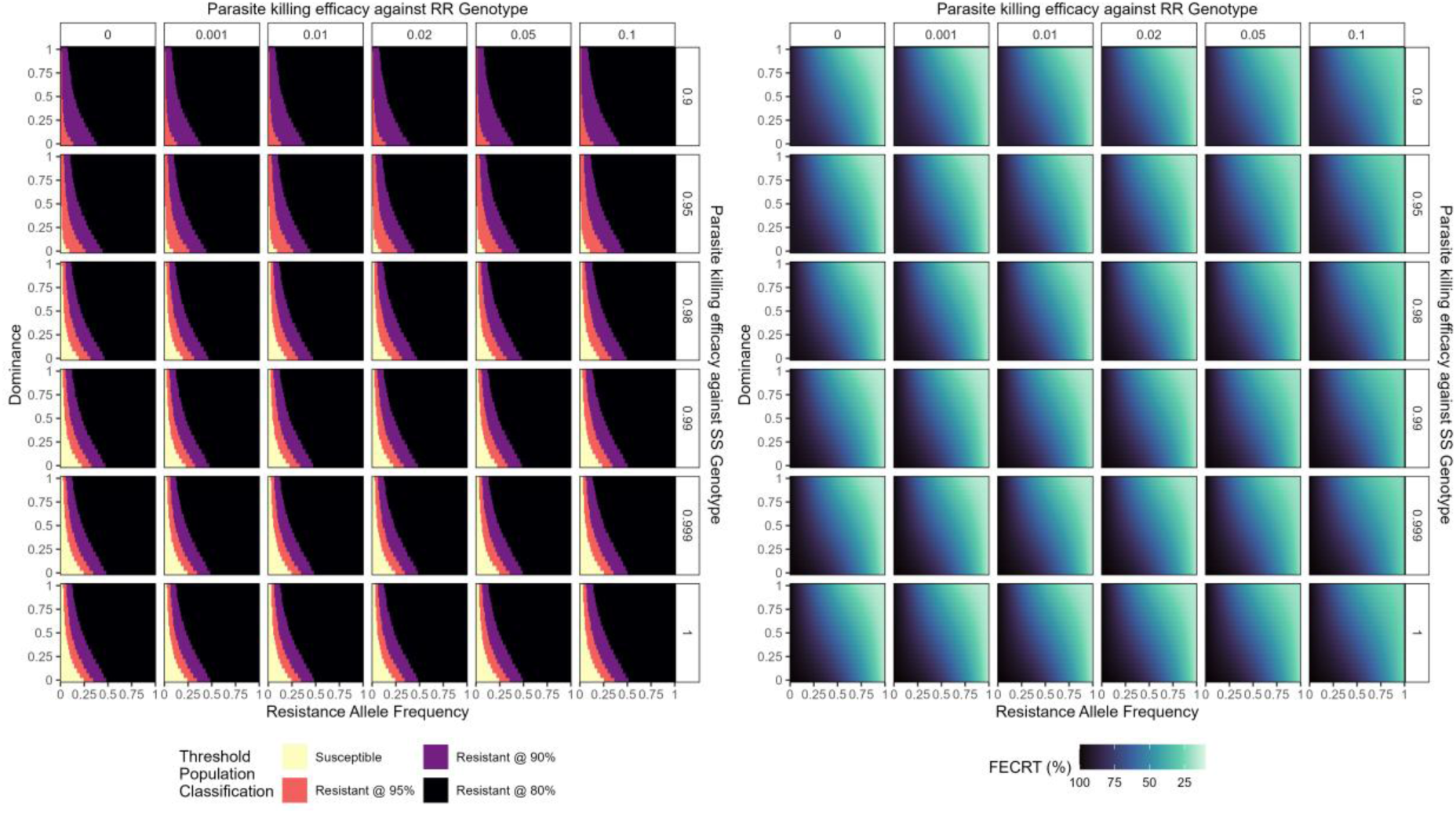
Faecal Egg Count Reduction Test given Drug Efficacy and Resistance. Both panels share the same overarching format: the resistance allele frequency is the x-axis, the dominance of resistance is the y-axis, the parasite killing efficacy against the RR genotype is represented by panels from left-right; and the parasite killing efficacy against the SS genotype is represented by panels from top-bottom. The left-hand plot shows the FECRT resistance categorisation assuming various threshold faecal egg count reduction percentages (Coles et al., 1992; Kaplan, 2023).

The parameters explored were:

- *Resistance frequency:* Each drug (A and B) can have a starting resistance allele frequency: 10^-6^, 0.01, 0.1 or 0.25; and all permutations are explored. We explored a wider range of resistance frequencies to allow for a better understanding of how using two different drugs with different initial frequencies influences ARM performance.

The strategies evaluated were:

- Monotherapy: a single anthelmintic (drug A) is given during each treatment event. This acts as the baseline non-strategy comparator.
- Monthly pharmacological rotation: the drugs (A and B) are alternated at each treatment event.
- Yearly pharmacological rotation: the drugs (A and B) are alternated yearly. Such that all treatments in year 2 are with drug A, and all treatments in year 3 are with drug B.
- Fixed cohort mosaic: Livestock receiving treatment are assigned to either to cohort 1 or cohort 2. Cohort 1 receives drug A for each treatment. Cohort 2 receives drug B for each treatment.
- Monthly rotation cohort mosaic: Livestock receiving treatment are assigned to either to cohort 1 or cohort 2. Cohort 1 receives drug A for the first treatment, then rotating between drugs A and B at each treatment. Whereas Cohort 2 receives drug B for the first treatment, then rotating between drugs A and B at each treatment. Cohort 1 and 2 never receive the same drugs at the same treatment.
- Yearly rotation cohort mosaic: the yearly rotation equivalent to the monthly rotation cohort mosaic described above.
- Combinations: where both drug A and drug B are combined in a single formulation and livestock therefore receive both drugs simultaneously.
- Combinations with reduced coverage: Each round of treatment covers 70% of the livestock population.
- Combinations with reduced frequency: Only 5 treatments are given each year, the July 1^st^ treatment (the last treatment of the year) is not given.

This generates a total of 144 resistance-strategy permutations which are run over 100 geographic locations to account for the impact of climate on GIN population dynamics (total n = 14,400 simulations).

We report the percentage differences in the outcome measures when compared against the baseline monotherapy deployment for each timestep (*t*):

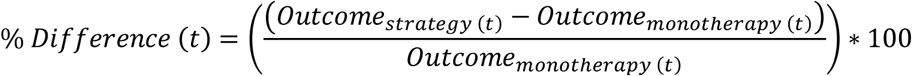

Therefore, when the % *Difference*(*t*) is positive this indicates the monotherapy performed better, and when the % *Difference*(*t*) is negative the comparator strategy performed better when evaluating for the specified outcome measure. The overall performance of any strategy is the composite of these differences across all outcomes, however where the challenge comes is in weighting the outcomes against each other, and their importance depends on the time horizon and priority of the strategy in question.

## 3. Results

### 3.1 Simulation Set 1 Results: Resistance classification and linking FECRT to resistance frequency and treatment efficacy

Figure 3 (left-hand plot) shows the classification of GIN populations based on FECRT, when considering a range of resistance frequencies, resistance intensities and dominance alongside some pre-defined thresholds which constitute the target GIN population is resistant or susceptible. The right-hand plot of Figure 3 conversely reports the corresponding exact FECRT results, providing a more nuanced picture on how resistant the target GIN populations are.

Phenotypic classification varied with dominance; increasing the dominance of resistance alleles in heterozygotes resulted in phenotypic population classifications of resistance at a lower resistance allele frequency (Figure 3, left panels), thus, knowledge of the dominance of resistance alleles is required to infer resistance phenotype from molecular investigations. Likewise, where resistance phenotype and allele frequency are known, inference can be made regarding possible dominance.

Where dominance is low and drug efficacy against homozygote susceptible GINs is high, phenotypic measures of resistance (FECRT) did not meet the threshold for classification of resistance until resistance allele frequencies were as high as 25%, even under the most conservative threshold of 95% FECRT (Figure 3, bottom left panels).

### 3.2 Simulation Set 2 Results: Impact of genetic and drug efficacy properties on parasite control and resistance spread

Increasing the proportion of the in-host parasitic stages *in refugia* by increasing the proportion of livestock left untreated (the different coloured lines in Figure 4) increases the overall FEC compared with the baseline of whole-group treatment (0% left untreated). In subsequent years, this results in a similarly elevated larval pasture count compared with baseline, and therefore subsequent transmission intensities are higher. The benefit of maximising *refugia,* though, is demonstrated in the simulations as slowing the increase in resistance intensity (allele frequency), and this effect can be considerable when also considering drug decay.

**Figure 4.**
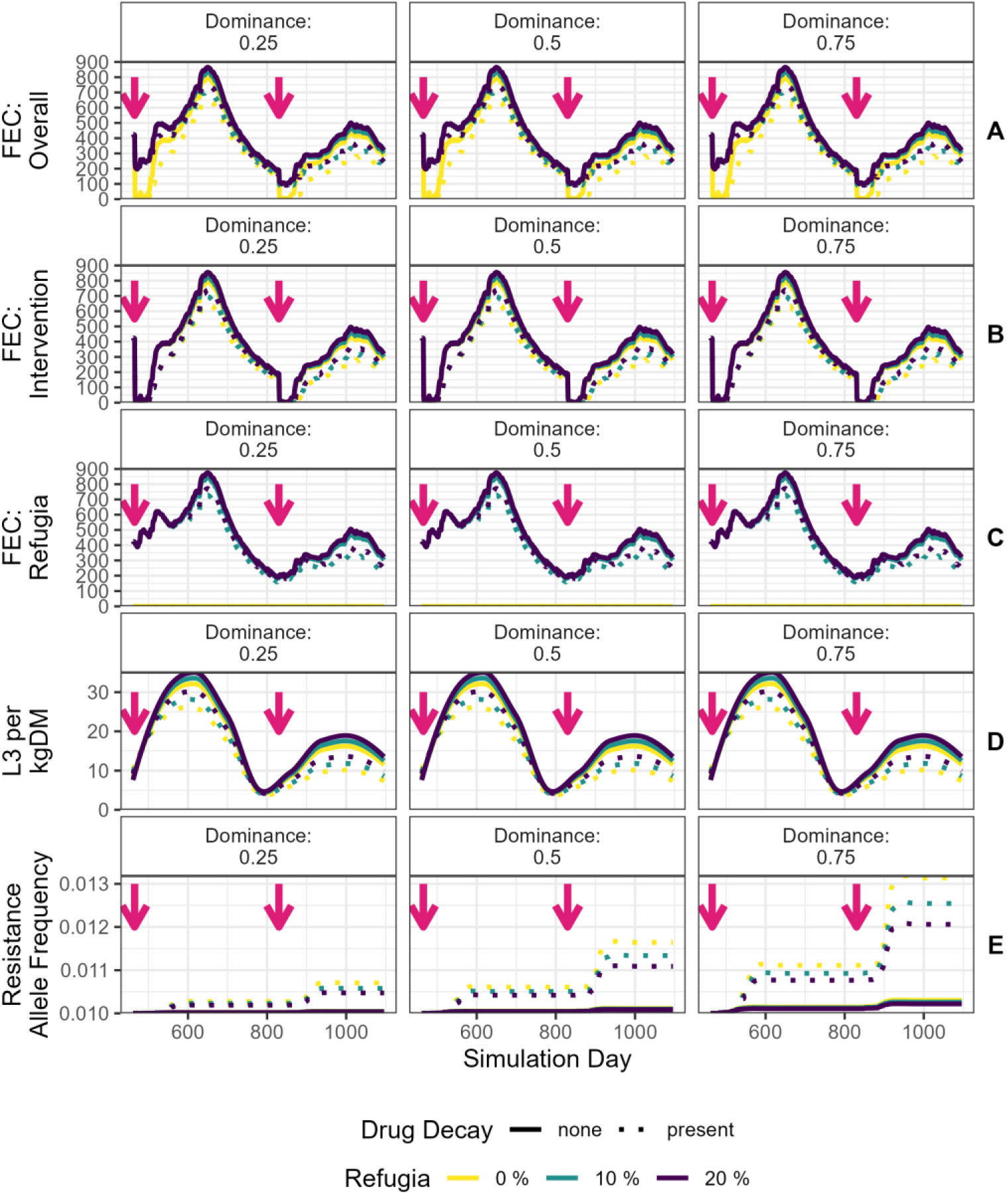
Results for Simulation Set 2: Influence of parameters on transmission and resistance. In all panels the colour of the lines indicates the proportion of the total livestock population that is untreated (*refugia*), from 0% (yellow) to 10% (green) and 20% (purple). The line type indicates whether drug decay was not (solid line) or was (dashed line) included in the simulations. The top row (A) shows the mean faecal egg count for the whole livestock population (treated and untreated), and is the mean value from the 100 geographic locations. The second row (B) shows just the mean value for the treated cohort, and the third (middle) row of plots (C) shows the mean FEC for untreated cohort (*refugia* population). The fourth row (D) shows the smooth mean value (using loess smoothing) for the L3 larval densities on pasture. The bottom row (E) shows the change in the resistance allele frequency over time. Plots are restricted to only showing years 2 and 3 of the simulation as year 1 is used as burn-in to seed the simulation. The pink arrows indicate the days the drug treatments are administered. The plots are stratified left-right by dominance. Only initial genotype frequencies of SS = 0.9 and RR = 0.1, and the higher transmission settings are shown in this plot, the full results are displayed in the Supplementary File (Section S3), but importantly do not alter the qualitative conclusions which can be drawn.

Including drug decay (the dashed lines in Figure 4) has the short-term epidemiological benefit of treatment being improved (as the killing time is extended, even if diminished, as shown by the dashed lines in the FEC plots in Figure 4 being lower than the comparator solid lines where drug decay was not included). Second, but far more importantly, is that this benefit is offset by the drug decay tails increasing selection for resistance, see the bottom panels of Figure 4 where the dashed lines (decay present) are notably higher than when drug decay was not included). The impact of drug decay on the development of resistance, as demonstrated by changes in resistance alleles over time, was also positively influenced by the predicted dominance of resistance alleles in heterozygotes, with higher dominance levels associated with substantial differences between simulations with and without decay.

Exploring the impact of transmission intensity and initial frequency of resistance (Figures S2.3 to S2.7) show that there remains a similar pattern. However, we note that in higher transmission settings the pre-specified treatment plan may not be sufficient to adequately control parasites and so treatment number would perhaps need increasing to compensate for this.

### 3.3 Simulation Set 3 Results: Evaluating strategies for parasite control and managing resistance

Figures 5 and 6 show the results for the simulations directly evaluating ARM strategies. To aid reading between all the plots, the colour scheme and plot layout is consistent between all plots and differ only in the outcome being evaluated (Figure 5 is FEC, and Figure 6 is resistance allele frequency).

**Figure 5:**
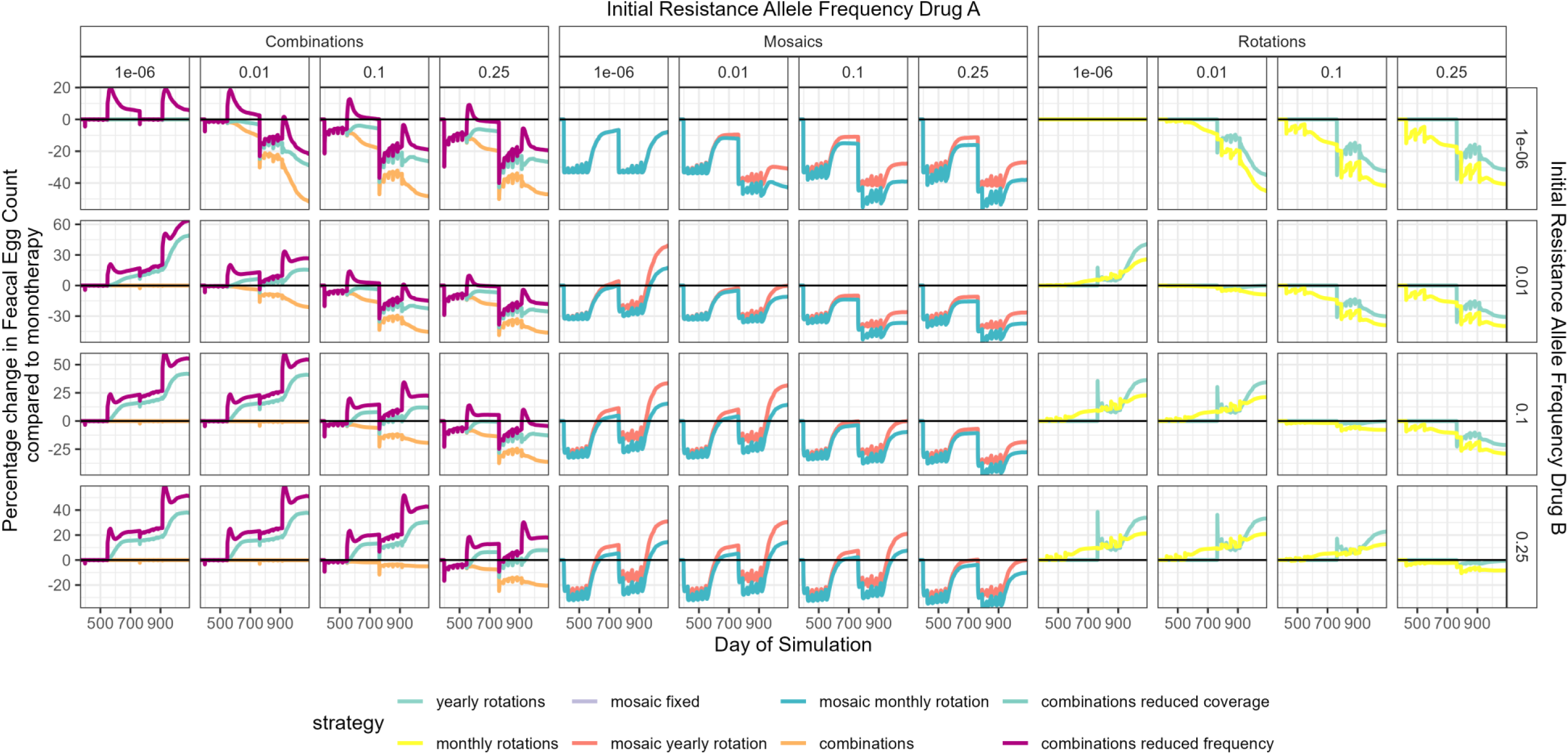
Simulation Set 3 – Percentage Difference in the Faecal Egg Count Compared to Monotherapy. The y axis is the percentage difference, as calculated from Equation 1. The horizontal black line (at zero) represents no difference between strategies. Values below the black line indicate the proposed strategy performed better, and values above performed worse than the Monotherapy comparator. The colour of the lines indicates the specific strategy. The x-axis is the simulation day and is restricted to not include the 1-year burn-in, which is the same across all strategies. Each line is the mean of 100 simulations, covering a range of geographic locations.

**Figure 6:**
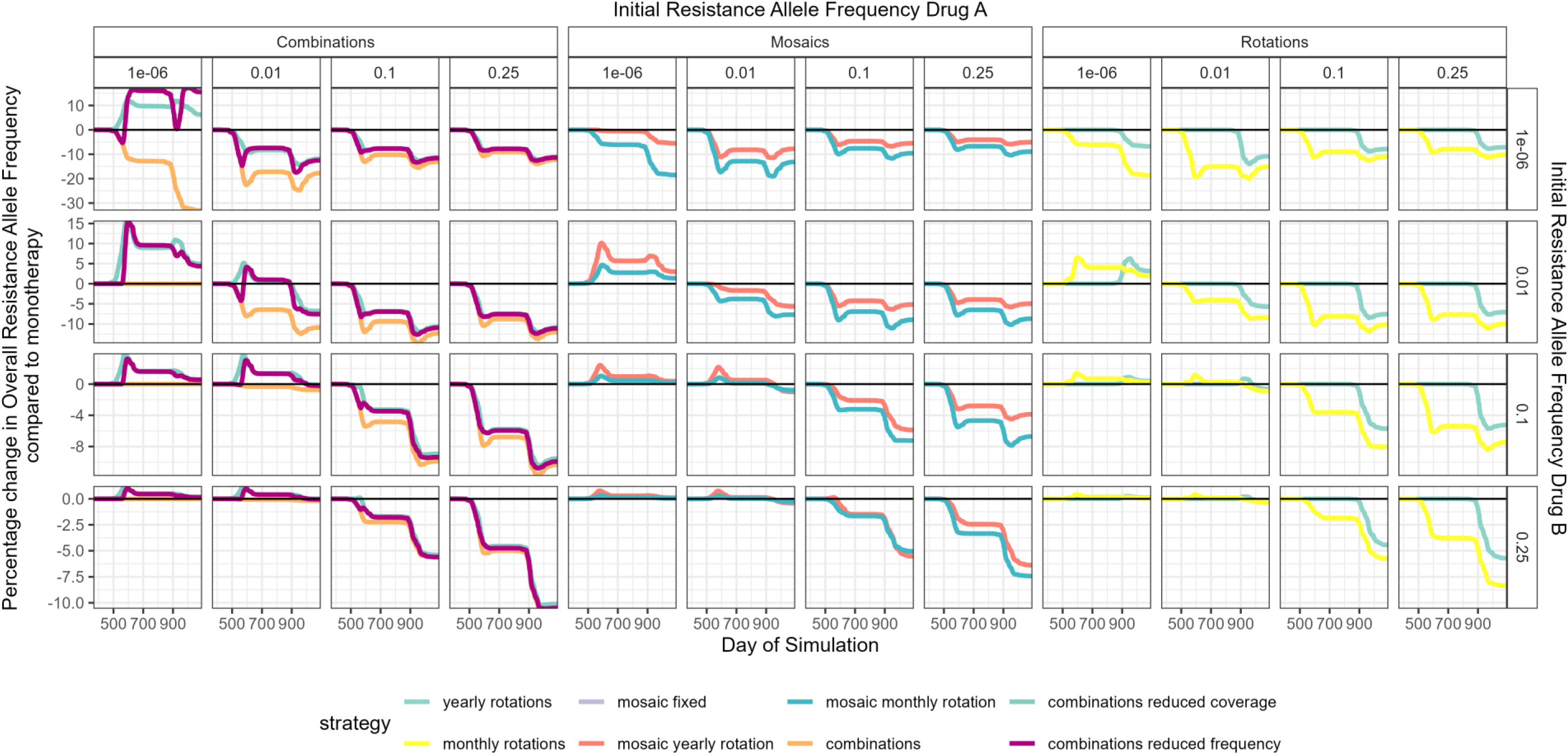
Simulation Set 3 – Percentage Difference in the Resistance Allele Frequency Compared to Monotherapy. The y axis is the percentage difference, as calculated from Equation 1. Note, Drug 1 is the drug used in the monotherapy simulations, and Drug 2 is not deployed in the monotherapy simulations. The horizontal black line (at zero) represents the no difference between strategies. Values below the black line indicates the proposed strategy performed better, and values above performed worse than the Monotherapy comparator. The x-axis is the simulation day, and is restricted to not include the 1-year burn in which is the same across all strategies. Each line is the mean of 100 simulations, covering a range of geographic locations.

The effectiveness of simulated ARMs in reducing FEC, was highly dependent on the initial resistance allele frequency for both drugs. When resistance to the monotherapy drug (drug A) was lower than drug B the ability for alternative strategies to suppress FECs was often impaired (Figure 5), highlighting that using the most effective anthelmintic treatment for parasite control is recommended. When drug B had lower initial resistance than drug A (e.g. a scenario where a “new” anthelmintic is used in rotation with an “older” anthelmintic) using strategies that had both drugs in use was generally beneficial in reducing FEC (Figure 5). When initial resistance allele frequency to each drug was equal, then strategies that used both drugs were beneficial, except the combinations that had either reduced coverage or reduced frequency. Although it should be noted, the absolute differences in FECs were not that different (Figure S3.2 in the Supplementary File).

When considering the impact on resistance, all strategies were found to be better than monotherapies, except for reduced coverage and reduced frequency combinations when resistance was very low (Figure 6); this is because reduced coverage and reduced frequency kills fewer worms, reducing the dilution benefit from L3 *in refugia* on pasture. Combination therapy outperformed all other strategies from a resistance management perspective, especially when no constraints are applied. Cohort mosaics performed surprisingly well from a parasite control perspective (Figure 5), consistently reducing FECs compared with baseline monotherapy. While the combination therapies did not necessarily outperform other strategies epidemiologically (when looking at FEC, Figure 5), the performance of combinations when considering resistance management was generally better than other strategies (Figure 6). This is an issue which would unlikely be considered in clinical trials or in the development of clinical guidelines focussing on measures of intensity of infection (FECs) as their outcome of interest.

## 4. Discussion

We presented results from an extended GLOWORM model framework, which included AR genetics and host cohorts, providing a novel evolutionary-epidemiological modelling framework for GIN control in ruminant livestock. These results provide avenues for a refined, high-level discussion regarding ARM and interpretation of molecular studies of AR. When considering evolutionary-epidemiological dynamics with respect to ARM, there are two outcomes of interest which are often in conflict: parasite control and resistance management. ARM guidelines need to provide practical and actionable recommendations without compromising parasite control, livestock welfare and productivity, and input costs (Learmount et al., 2015).

While short-term epidemiological dynamics are readily observable in trials (e.g., Little et al., 2010), the longer-term evolutionary dynamics of AR are often less tractable. This is in part because trials are focused primarily on whether an intervention is effective in the moment, because resistance management strategies cannot be considered if they fail at the primary task of reducing the immediate parasite/transmission burden, and due to the lack of genetic markers for resistance for some anthelmintic classes (McIntyre et al., 2025). Evolutionary-epidemiological models can be used to make qualitative statements regarding the comparative performance of strategy options, and to generate broad guiding principles behind resistance management.

Our simulations suggest: that resistance phenotype is an imperfect measure of resistance allele frequency; that pharmacokinetics (specifically residual activity) strongly influences the outcome of ARM; and that the balance between long-term ARM and effective parasite control in the short-term depends on genetic factors underpinning resistance within these parasite populations. We made two notable assumptions in our modelling approach:

First, our simulations gave treatments at set times regardless of the level of parasitism experience by the livestock. Strategies which perform worse at clearing parasitic infections may require repeated treatments selecting further for resistance; and similarly increased transmission would lead to re-infection and therefore the need to re-treat; which explains our findings that transmission intensity had minimal impact of resistance propagation.

Second, we assumed resistance to be monogenic, a common assumption across a range of models and diseases (Barbosa et al., 2018; Helps et al., 2020; Madgwick and Kanitz, 2022; Patel et al., 2024; Reichert et al., 2023; REX Consortium, 2013). Where models assume polygenic resistance (Coffeng et al., 2024; Hobbs et al., 2023; Hobbs and Hastings, 2025b) they are giving qualitatively similar results to their monogenic equivalents (Levick et al., 2017; South and Hastings, 2018) regarding strategy evaluation, which gives confidence that assumptions regarding the genetics of resistance do not drastically alter the qualitative performance of resistance management strategies, although the magnitudes of benefit may differ.

We demonstrated how minor changes in resistance intensity (the level of resistance from the RR genotype), dominance and resistance frequency can change whether a population would be categorised as resistant based on established thresholds (Figure 2, left panel), masking far less drastic changes in anthelmintic efficacy (Figure 2, right panel). This challenges the tendency for practitioners, policy makers and scientists to apply thresholds to a quantitative continuum and classify whole populations as resistant or susceptible. Although this is useful for prompting early farmer behaviour change in favour of ARM strategies, the variability in potential resistance allele frequency at these resistance thresholds, as shown in our simulations, may undermine ARM strategies reliant on a “low” initial resistance allele frequency (Bartram et al., 2012). With an increase in modelling resistance as a quantitative trait [15, 28], the communication of resistance as a continuum of phenotypes is starting to make headway. Without these more refined considerations of what the “resistance” problem is, assessing it becomes challenging.

Considering the importance of drug decay in impacting resistance dynamics in other systems (Stepniewska and White, 2008), failures to include drug decay may mean we are not fully understanding ARM strategy performance. Of note is the importance of dominance and drug decay. High levels of dominance combined with drug decay hugely increases the change in the resistance allele frequency; these are two parameters not widely investigated and the effect of them appears to be as influential as the inclusion of *refugia* in influencing resistance dynamics. Thus, inclusion of drug decay in cost-benefit analyses is essential.

Of note is that, when dominance is high and drug decay is included, this leads to a situation allowing for rapid spread of resistance. This is an especially important consideration, as many new anthelmintic drug formulations are intended to have long efficacy durations, meaning there is ample time for the drug to decline in concentration. We also assume there are no mistakes made in dose of the anthelmintics given, which would be most likely be due to underdosing in a practical setting. Under-dosing has two notable potential implications. The first is the impact on the evolution of resistance. The second is having an incorrect estimate of the resistance phenotype of the target population, as the reduced observed efficacy was due to under-dosing and not resistance. Whilst our simulations are agnostic to product, there is a need to further investigate the use of long-lasting formulations when used for targeting non-GIN parasites, such as moxidectin and sheep scab. It is critical that researchers and policy-makers utilising the same interventions communicate and coordinate their resistance management strategies, for example, SCOPS have consolidated quarantine recommendations for nematode and psoroptic mange control (Stubbings et al., 2020). The two-gene module (AR2) developed here allows for future exploration of the combined impacts of anthelmintic treatments applied for both GIN and ectoparasite control on the development of resistance.

One often highlighted component of ARM strategies is to maintain a “*refugia*” (Hodgkinson et al., 2019), where untreated livestock or pre-contaminated patches can act as a source of susceptible alleles. A key aspect is considering what level of pasture contamination is needed to provide an effective *refugia*, and what level of pasture contamination is possible without allowing for rapid reinfection and the need to re-treat livestock. One of the current core tenets of resistance management is to reduce the usage of, and reliance on, anthelmintic treatments (Stubbings et al., 2020). Although a degree of reduced anthelmintic use and implementation of *refugia-*based strategies without production impacts is possible (Learmount et al., 2015), it is clearly important to note that this may come at the cost of increased transmission as the target GIN parasites may no longer be as effectively controlled, as was evident in our simulations considering reduced coverage and frequency of anthelmintic treatments (Figure 5). This is consistent with industry messaging around using as little anthelmintic as possible to reduce AR selection but at the same time as much as necessary to control disease (Loeb, 2019). By moving livestock to contaminated patches to allow for reinfection, this increases the need for future treatments, which themselves provide further selection for resistance.

The challenge is to balance epidemiological goals, which are often short-term, with resistance management goals, which consider far longer time horizons, with no guarantee of pay-off, due to the inherent stochasticity of evolutionary processes. Previous modelling studies have identified potential tipping points whereby the benefit of reduced rate of development of resistance is offset by increased FEC (Berk et al., 2016) or increased economic costs (Pech et al., 2009) when considering reduced coverage and frequency of treatment. This is consistent with our simulations, which further suggest that dominance appears to be a far more important consideration than *refugia* in the absence of drug decay (Figure 4). Interestingly, our simulations show that *refugia* is more important to consider when there are long tails of drug decay (Figure 4); therefore guidelines for treatment may need to consider whether short course treatments are given, whether the drug is rapidly metabolised (and so the decay time is short), and whether in these circumstances universally treating all livestock may be preferable due to no appreciable resistance management benefit to offset the reduced clinical benefit.

When faced with the likely financial constraints of ARM strategies, e.g., combinations (due to their higher costs), the simulations presented here offer several options for further exploration. For example, from both a resistance management and parasite control (FEC reduction) aspect, it appears that reducing the coverage of anthelmintic treatments is preferable to reducing the frequency of treatments. Although we note we did not fully balance these two for equal financial cost. Furthermore, some options such as cohort mosaics are likely to be challenging to implement, and did not appear to offer a resistance management benefit to offset this challenge (Figure 5); especially when compared to combinations.

Mathematical models may well be able to identify the optimal pasture contamination, but a key question becomes, how robust does the *refugia* strategy become outside any optima, and more importantly, can optimally identified conditions be practically implemented by farmers? The design and implementation of resistance management strategies need to be robust even when implemented incorrectly, sub-optimally, or in the face of incomplete knowledge e.g., regarding the genetic basis of resistance (McIntyre et al., 2025). Ensuring strategies are implementable and robust regardless of variable conditions relating to the climatic/biological/operational nature of the system is a core consideration e.g. evaluation of SCOPS guidelines on focal farms (Learmount et al., 2016).

We would note two factors which we did not investigate but are widely considered important, being fitness costs and cross resistance. For example, our presented simulations did not include fitness costs, despite “*A priori*, one expects such fitness cost” (Hodgkinson et al., 2019). Fitness costs are needed to maximise any benefit of *refugia.* Fitness costs are essentially always beneficial for any resistant management strategies, but strategies should not rely on them for effectiveness. Cross resistance is rarely included in models (REX Consortium, 2010). However in recent modelling of insecticide resistance which included cross resistance, this did not alter the choice between insecticide mixtures (equivalent to drug combinations) and rotations; remaining in favour of mixtures (Hobbs et al., 2023; Hobbs and Hastings, 2025b).

Further to the above ARM robustness considerations, although broad economic evaluations of anthelmintic usage and AR have been made (Charlier et al., 2020), economic evaluations of ARM have been limited to a subset of treatment scenarios (e.g., drenching frequency and *refugia* size (Pech et al., 2009) and allied strategies such as grazing management (van der Voort et al., 2017)). The former identifies potential tipping points whereby the benefit of reduced rate of development of resistance is offset by the economic costs. This observation relates to the fact that scenarios that result in higher transmission (and the higher resulting FECs) may lead to production impacts and more treatments being required, and this would further select for resistance. This was not explored in the present study, although further model extensions to represent recommended FEC-based treatment decisions (Stubbings et al., 2020) would allow further quantification of the treatment costs of different ARM strategies against the rate of development of resistance.

In conclusion, the interventions we use to control and treat parasitic diseases are also those supplying the selection pressure for resistance and there is a need to balance treatment frequency with selection frequency. However, the need to treat is also dependent on the level of transmission, which itself can be minimised by effective treatment. Resistance management is inherently interwoven into parasite control, albeit often as an afterthought. Discussions of anthelmintic resistance management must simultaneously consider parasite transmission, pharmacology and genetics. We provide a rigorous simulation framework to enable such discussions allowing for a refinement into our understanding of parasite control in the presence of resistance evolution. We also provide quantitative output to infer dominance from coupled molecular studies of resistance frequency and FECRTs, to further aid ARM strategy applications.

## Acknowledgements

This work was funded by the Biotechnology and Biological Sciences Research Council, project reference number: BB/X000923/1

## Supplementary file

### S1 Model Equations

We further provide the technical extensions of the GLOWORM (Rose et al., 2015; Rose Vineer et al., 2020) framework to include both resistance genetics and multiple livestock cohorts. Previously the GLOWORM framework was methodologically extended to include different patches (META) (Ingle et al., 2025), which allows for the movement of livestock populations between different patches. We now further add the COHORT and AR/AR2 extensions. Note, that META and COHORT can be combined with just the single gene AR extension. Practically, each extension can be used in isolation or together such that the current full GLOWORM framework (Figure S1) is: GLOWORM-META-COHORT-{AR/ AR2}. This flexibility allows for a range of pharmacological interventions to be investigated in an evolutionary epidemiological context (see Section S5 for examples).

**Figure S1.**
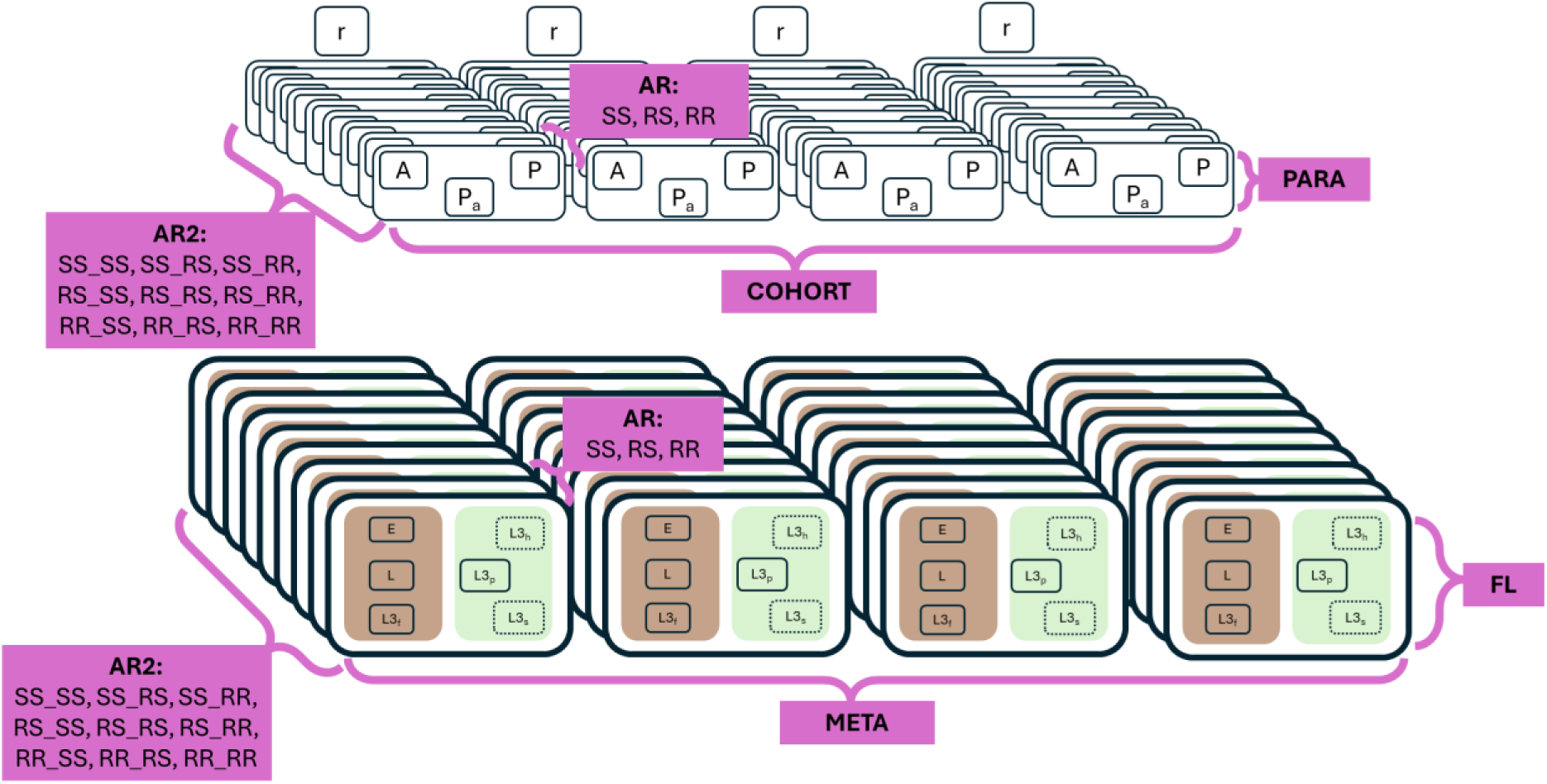
GLOWORM-META-COHORT-{AR, AR2} Modularisation. This is duplication of Figure 1 in main manuscript. The GLOWORM model framework consists of a modular format, allowing for the different extensions to be used in unique permutations. The FL (lower half of diagram) (Rose et al., 2015) and PARA (upper half of diagram) (Rose Vineer et al., 2020) are the core parasite transmission components of the model, used across all other model extensions. The META (Ingle et al., 2025) component extends to allow for separate patches for free-living parasites (represented by replication from left to right of the lower half of the diagram). The new COHORT extension allows for separate groups of grazing livestock (represented by replication from left to right in the upper half of the diagram). The AR and AR2 extensions are when the number of different anthelmintic genes being tracked is specified (represented by replication from front to back of the diagram, on the z axis).

#### S1.1 COHORT extension overview

The extension to multiple livestock cohorts (COHORT) allows for treatment regimens (and the dynamics this produces) to be specified different between parents and offspring livestock. Each livestock cohort may have unique properties (age, treatments, movements), and this allows for two key areas to be investigated: explicit treatment *refugia* and parent-offspring treatment dynamics. The livestock cohorts are denoted by the subscript *Cohort*. Each livestock cohort has its own within host dynamics, but different livestock cohorts can exist on the same patch at the same time, be given different treatments at different times, and importantly, the livestock cohorts can be different weights to allow the simultaneous modelling of parents and offspring. Livestock are assigned to one cohort, which they remain in throughout a simulation.

We can extend the model to include separate “cohorts” of livestock. For example, adult livestock and juvenile livestock, who have large discrepancies in weight, acquired immunity, and dry matter intake. We define a “cohort” as a group of livestock which remain constant in their composition throughout any simulation who have the same biological parameters (e.g., weight and dry matter intake), and also receive the same treatments at the same time. Within each simulation, it is possible to give the cohorts different biological properties, which is especially important when considering the transmission dynamics occurring between parental and offspring livestock. Following from (Ingle et al., 2025), we denote the patches i, and specify the patch any particular cohort is on as j.

#### S1.2 AR and AR2 extension overview

The capability to simultaneously and separately model resistance to two different active ingredients is required to evaluate the evolutionary and epidemiological implications of ARM strategies including drug rotations and drug combinations where different active ingredients target the same parasite species. We consider a single gene extension (AR) and a two gene extension (AR2). The genotypes of the parasitic worms are denoted by the superscript *genotype*. In AR *genotype* ∈ {*SS*, *RS*, *RR*}, such that there are 3 permutations, the homozygous susceptible (SS), heterozygote (RS) and homozygous resistant (RR). For the AR2 extension, where there are two genes, *genotype* ∈ {*SS*_*SS*, *SS*_*RS*, *SS*_*RR*, *RS*_*SS*, *RS*_*RS*, *RS*_*RR*, *RR*_*SS*, *RR*_*RS*, *RR*_*RR*}, such that there are 9 possible genotype permutations when considering a two-gene system. The first two letters (before the underscore) are for the first gene (which would be resistance to drug A), and the two letters (after the underscore) correspond to the second gene (which would be resistance to drug B). As is common in the resistance modelling literature we use *S* for susceptible, and *R* for resistant. The assumption of monogenic resistance is widespread throughout the resistance modelling literature regardless of biological system (e.g., (Helps et al., 2020; Levick et al., 2017; Madgwick and Kanitz, 2022; Reichert et al., 2023; REX Consortium, 2013)).

#### S1.3 Model Equations

##### S1.3.1 Model Equations – Resistance Allele Frequency

The distribution of the resistance genotypes of the eggs laid will depend on the resistance allele frequency of the adult worm population, which resides in the various livestock cohorts. Because each cohort can have unique patch movements and drug treatments the frequency the resistance allele frequency may differ between cohorts, and therefore must be calculated separately for each cohort separately. We can then calculate the resistance allele frequency (*q*) for each gene in the adult worm population, and this is conducted separately for each livestock cohort. Such that 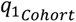 refers to the resistance allele frequency of gene 1 in the adult worm population in the livestock cohort *Cohort*, and 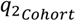 refers to the corresponding resistance allele frequency of gene 2. We must track the resistance allele frequencies in the adult worm population in the cohorts separately because different resistance allele frequencies may arise given different management practices between the livestock cohorts. The resistance allele frequencies in the livestock cohorts are subsequently used to calculate the distribution of the genotypes of the eggs laid by the adult worm population.

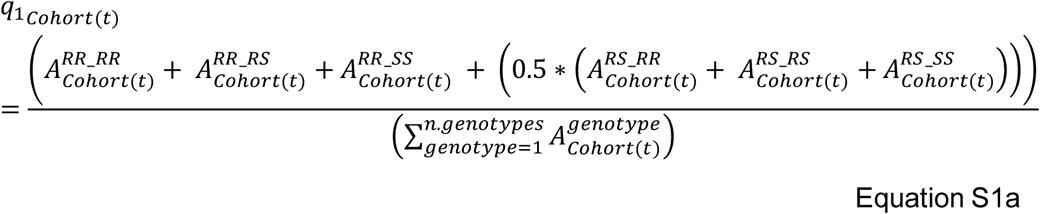

This is similarly calculated for the second gene:

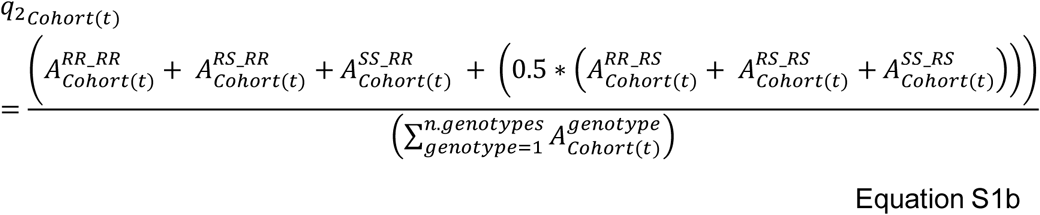

Note, the model can readily be adapted to track only a single gene:

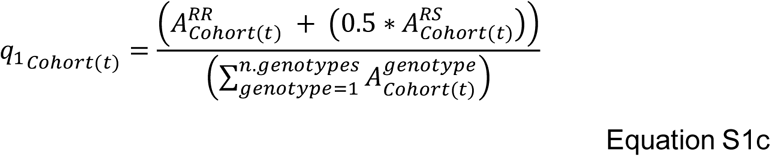

##### S1.3.2 Model Equations - Eggs

We next update the free-living worm equations for the eggs (Equations 2a to 2i) and free-living larval stages (Equation 3 to Equation 5). These equations can be adapted to a single gene format by removing all the *q*_2_ related terms.

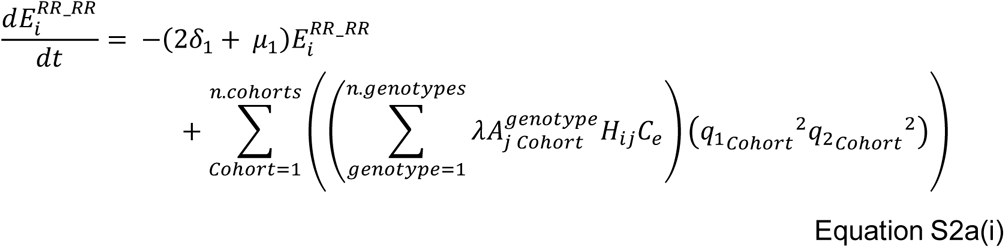

This equation tracks the number of eggs on patch *i*, which have the genotype RR_RR. 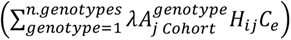 calculates the total number of eggs laid by a cohort on a patch, which is then converted to the number of the tracked genotype (in this case for the RR_RR genotype using 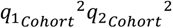. This is then wrapped in the 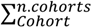 such that the total number of new eggs are counted across all cohorts.

These equations can be adapted to a single gene format by removing all the *q*_2_ related terms, for example:

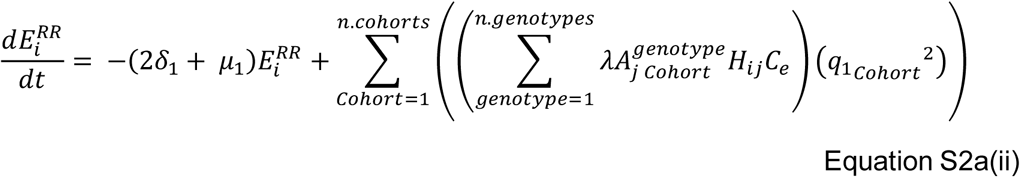

The following egg equations are similarly as above, differing in which genotype being tracked.

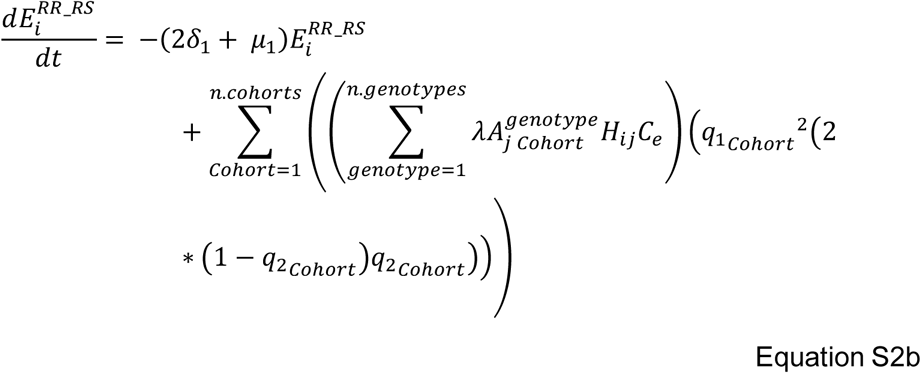

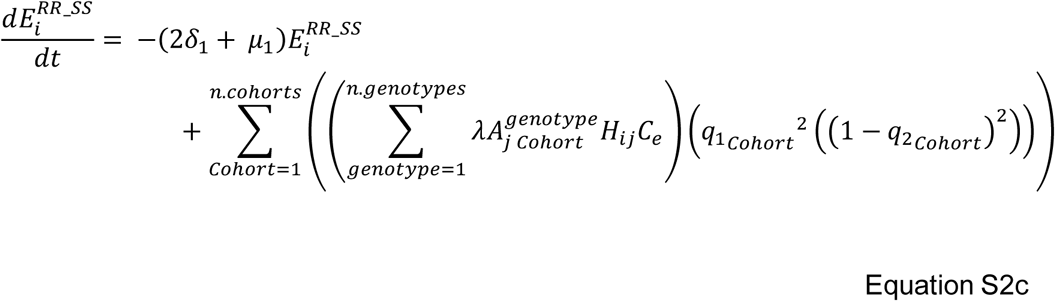

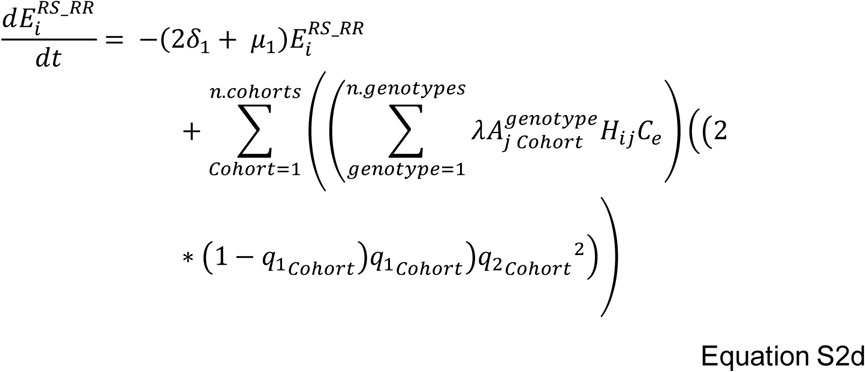

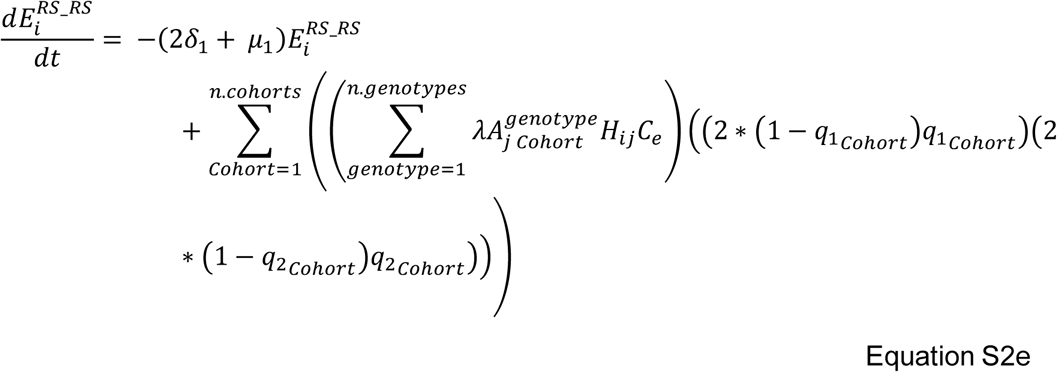

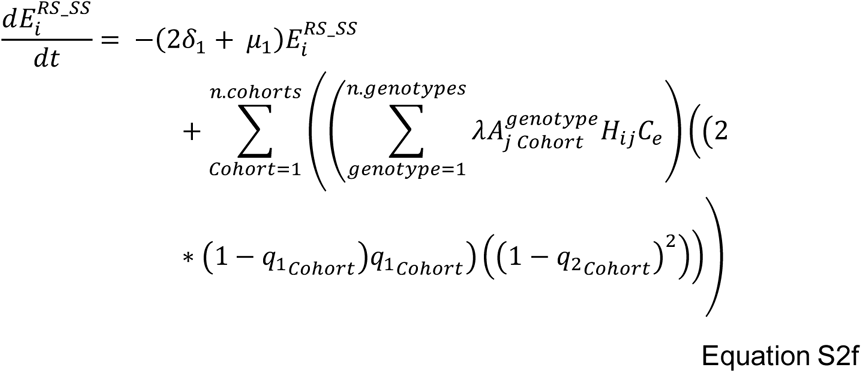

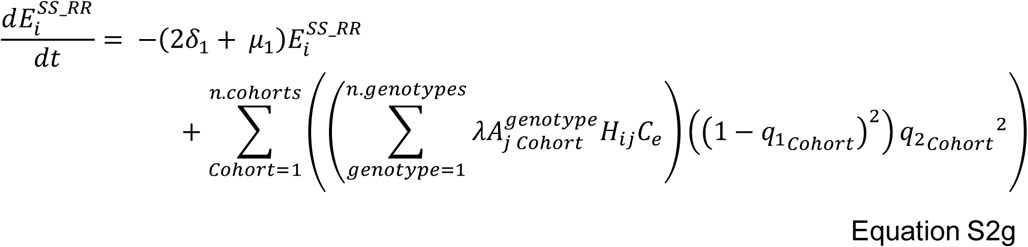

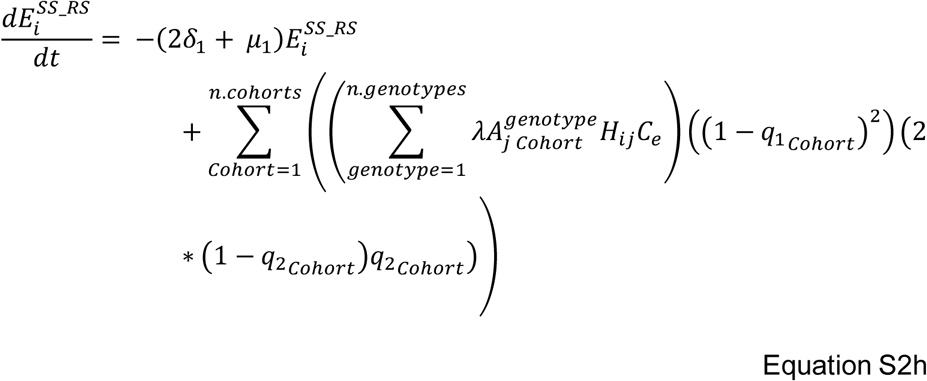

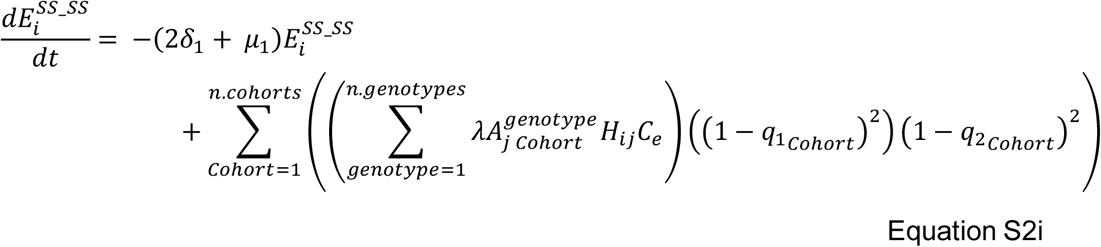

##### S1.3.3 Model Equations – Free-living larvae

The larval (L1-L2) equation is also updated to account for all genotypes, but can be written in a generalisable format, and is therefore the same equation as Equation 2 in (Rose et al., 2015) except for the extension to include the resistance genotype as a superscript, and the patch location denoted in the subscript.

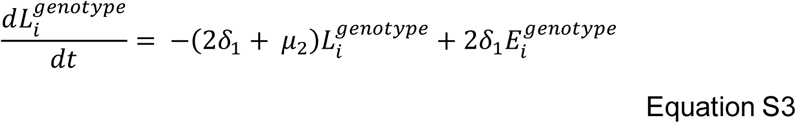

For the L3 larvae in faeces this can also be written in a generalisable format, and is therefore the same equation as Equation 3 in (Rose et al., 2015) except for the extension to include the resistance genotype as denoted in the superscript, and the patch location denoted in the subscript.

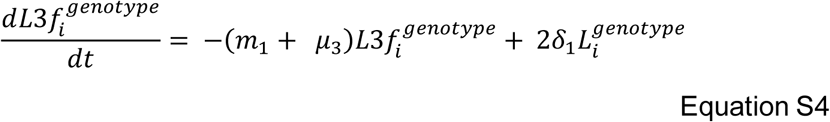

The number of larvae on pasture equation is therefore updated to account for the ability for different cohorts to feed on the pasture. It is key to note that we allow the dry matter intake to be different for each cohort (as specified by *β_Cohort_*). Here, as the equations for each genotype would be the same we write this as a generalisable form:

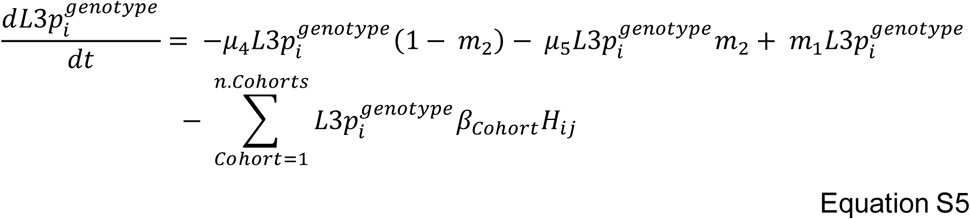

##### S1.3.4 Model Equations – Within-host

We next update the parasitic life-stages of the equations. For the parasitic life-stages (P, Pa, A), we must track the genotype of the worm (superscript Genotype), the cohort the worm resides in (subscript Cohort) and the patch the Cohort in on (subscript j).

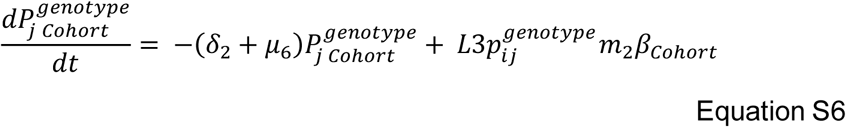

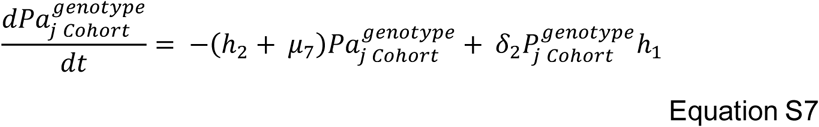

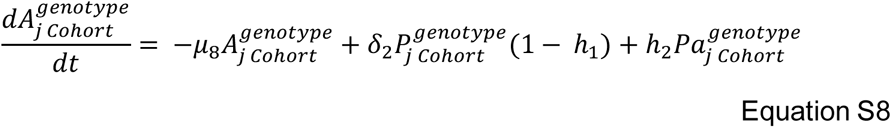

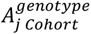 is therefore the number of adult worms of genotype *genotype* infecting the livestock cohort *cohort* who reside on patch *j*. 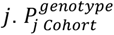 and 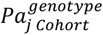 describe the same except for the Parasitic and Arrested life stages as previously detailed (Rose Vineer et al., 2020). It must be emphasised that the *genotype* superscript refer to the genotype of the worms infecting the cohort and not the genotype of the livestock cohort itself.

As each cohort can be given unique patch movements, treatment regimens or differences in dry matter intake (due to weight differences), immunity is calculated separately for each cohort.

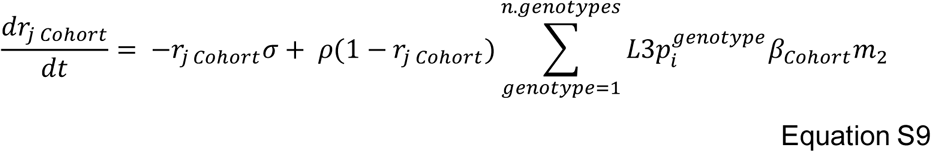

Other models have called this livestock immunity to parasites as “resistance”; but for clarity we use the term resistance only when describe a genetic process.

##### S1.3.5 Model Equations – Symbols and Descriptions

**Table S1.1.**
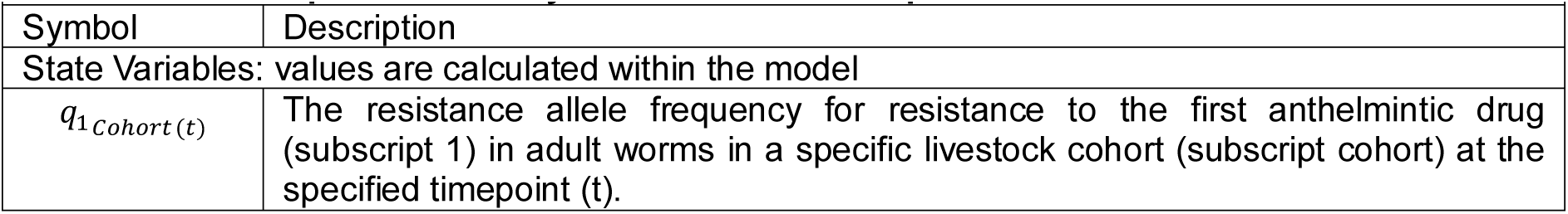
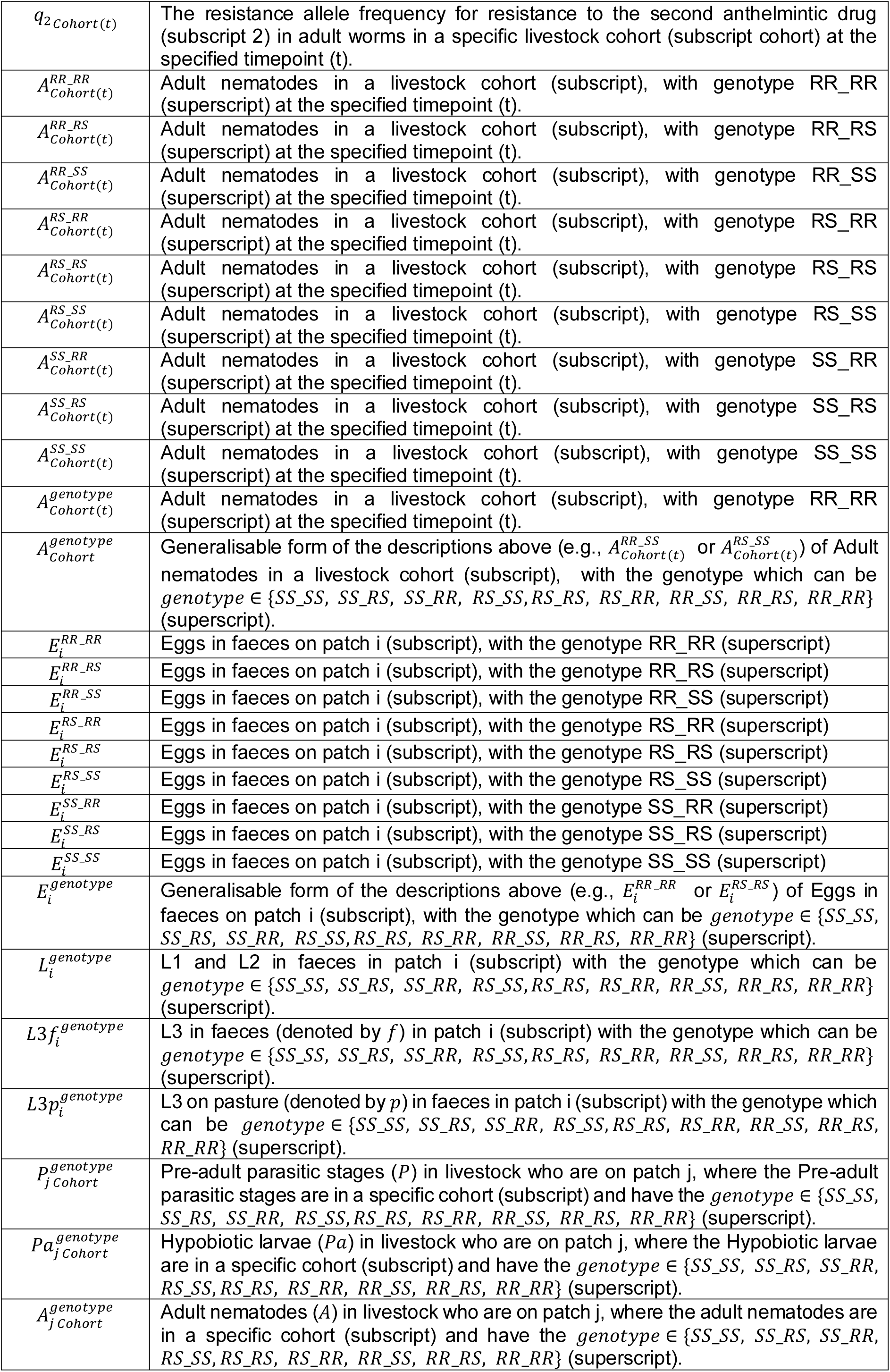
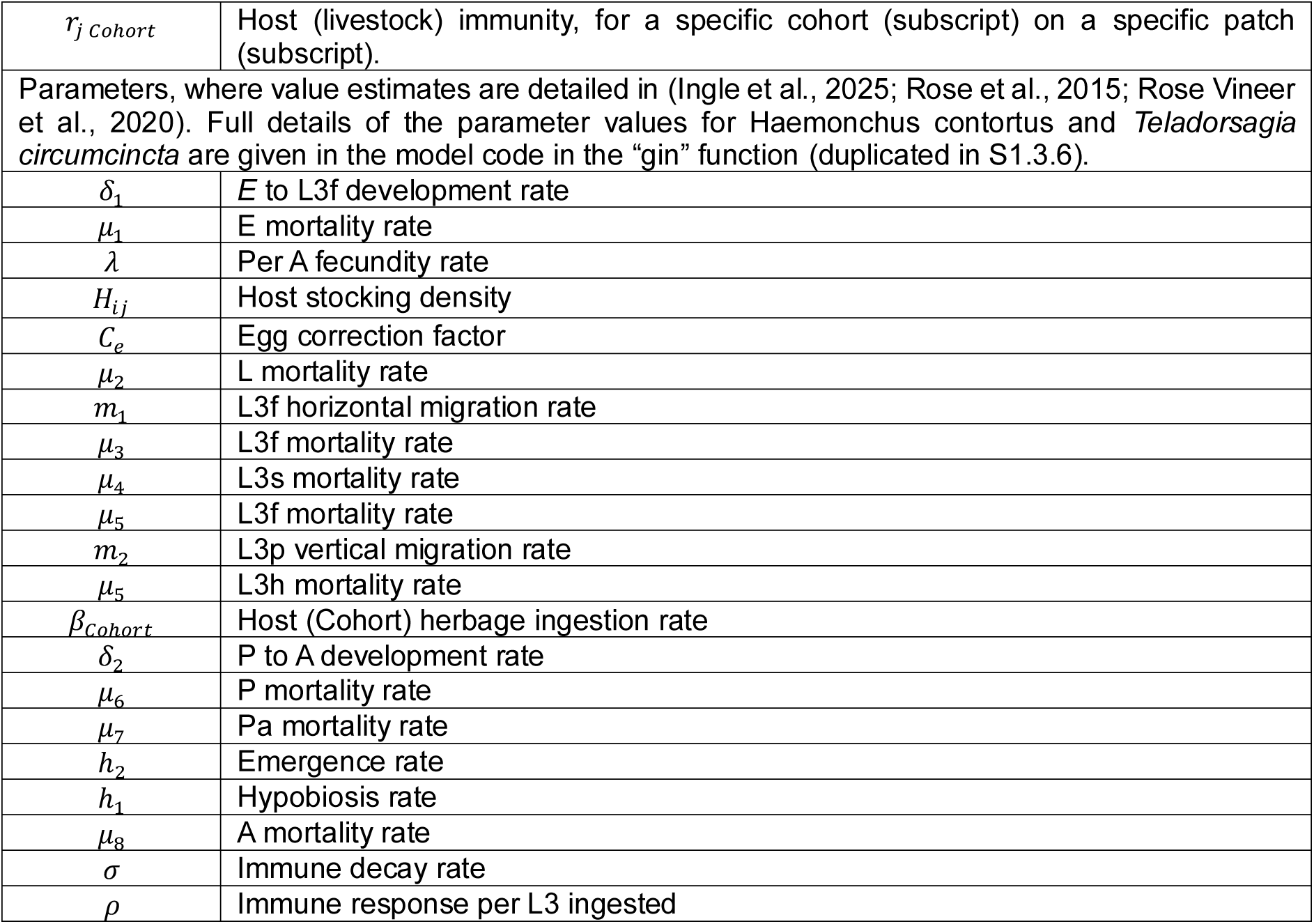
Descriptions of the Symbols used in the Equations.

##### S1.3.6 gin function

Printed out below is the gin function (written in R) which details the parameter values used for the respective parasite species, and therefore shows exactly how those values are calculated.

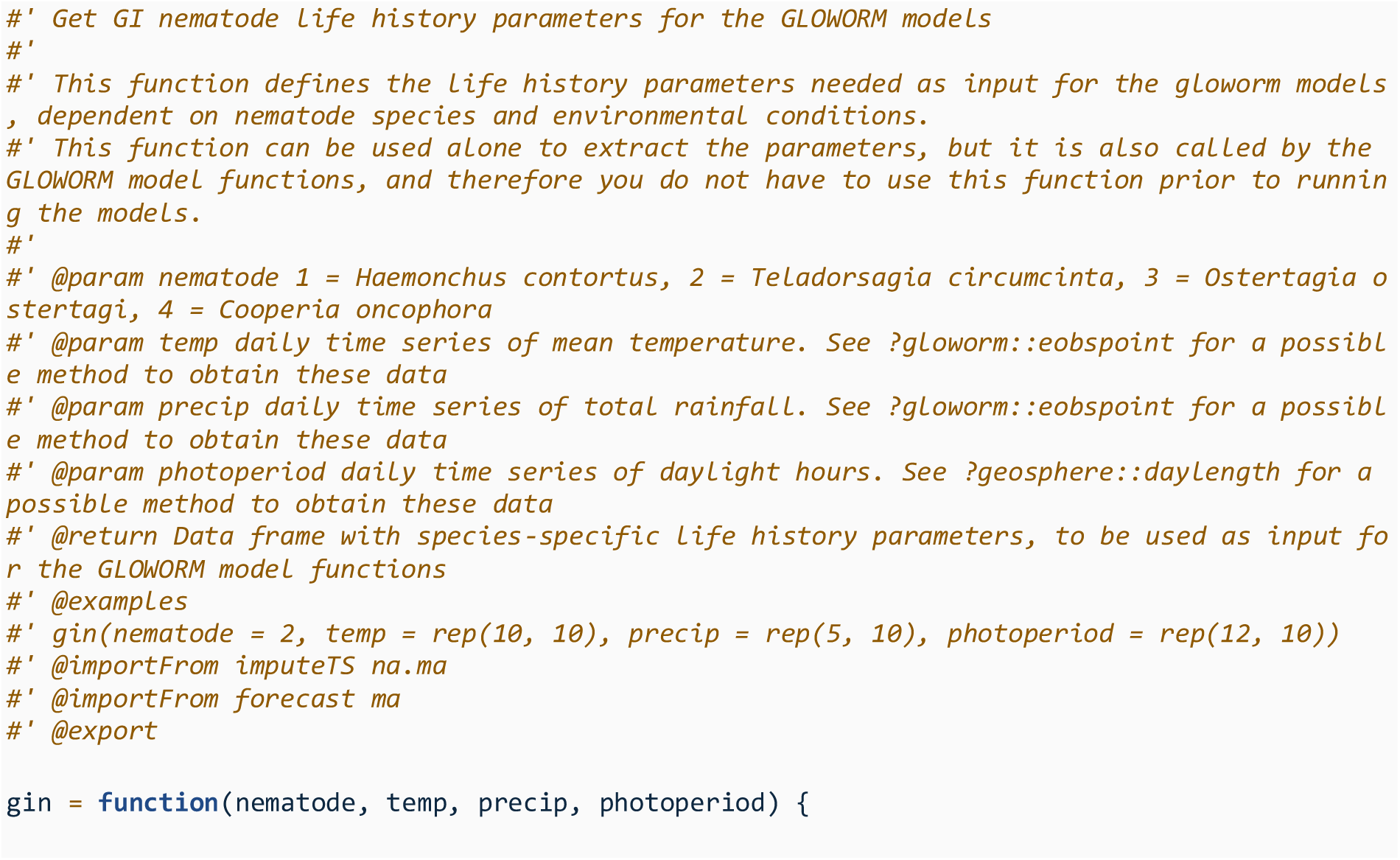
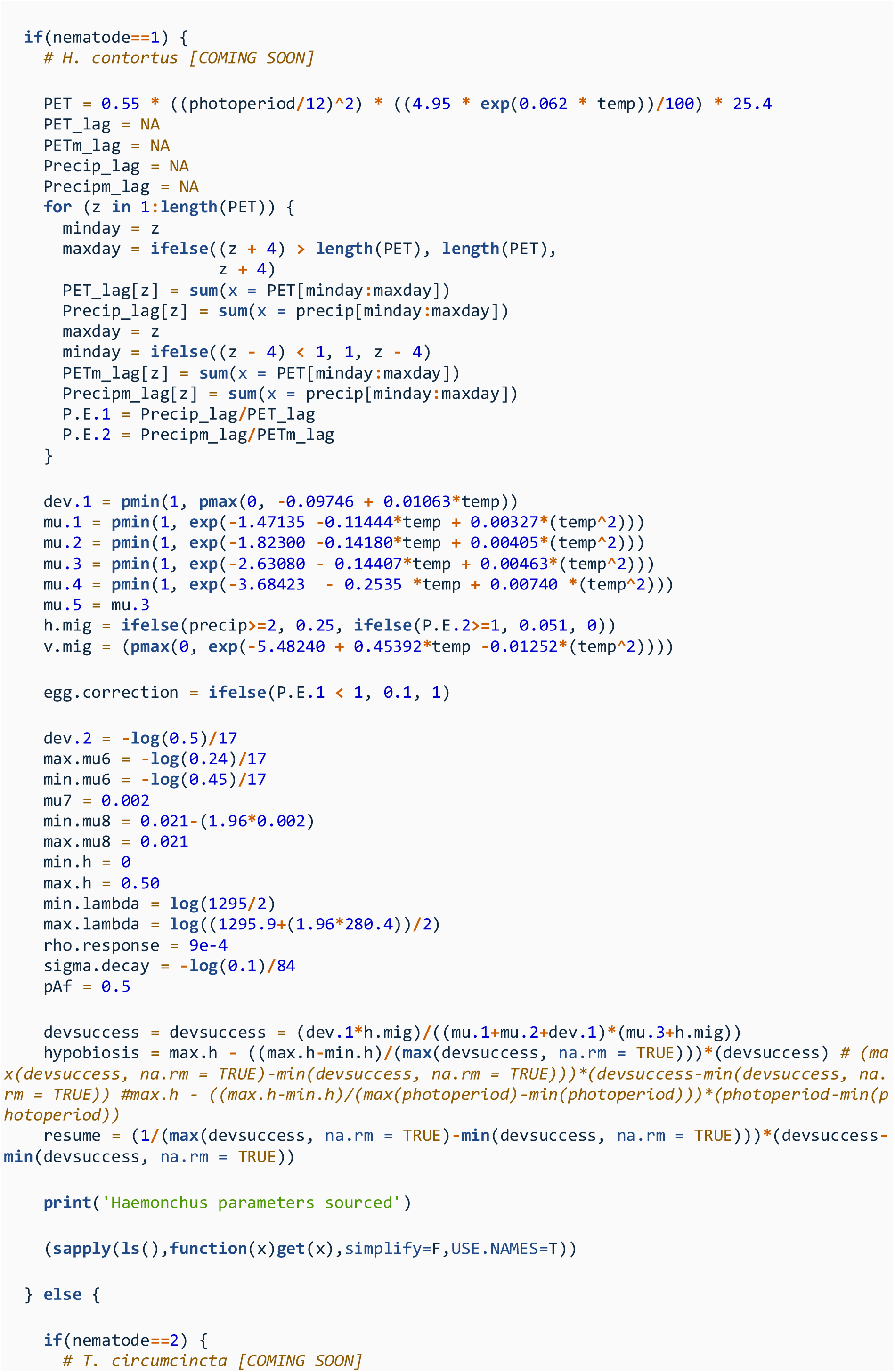
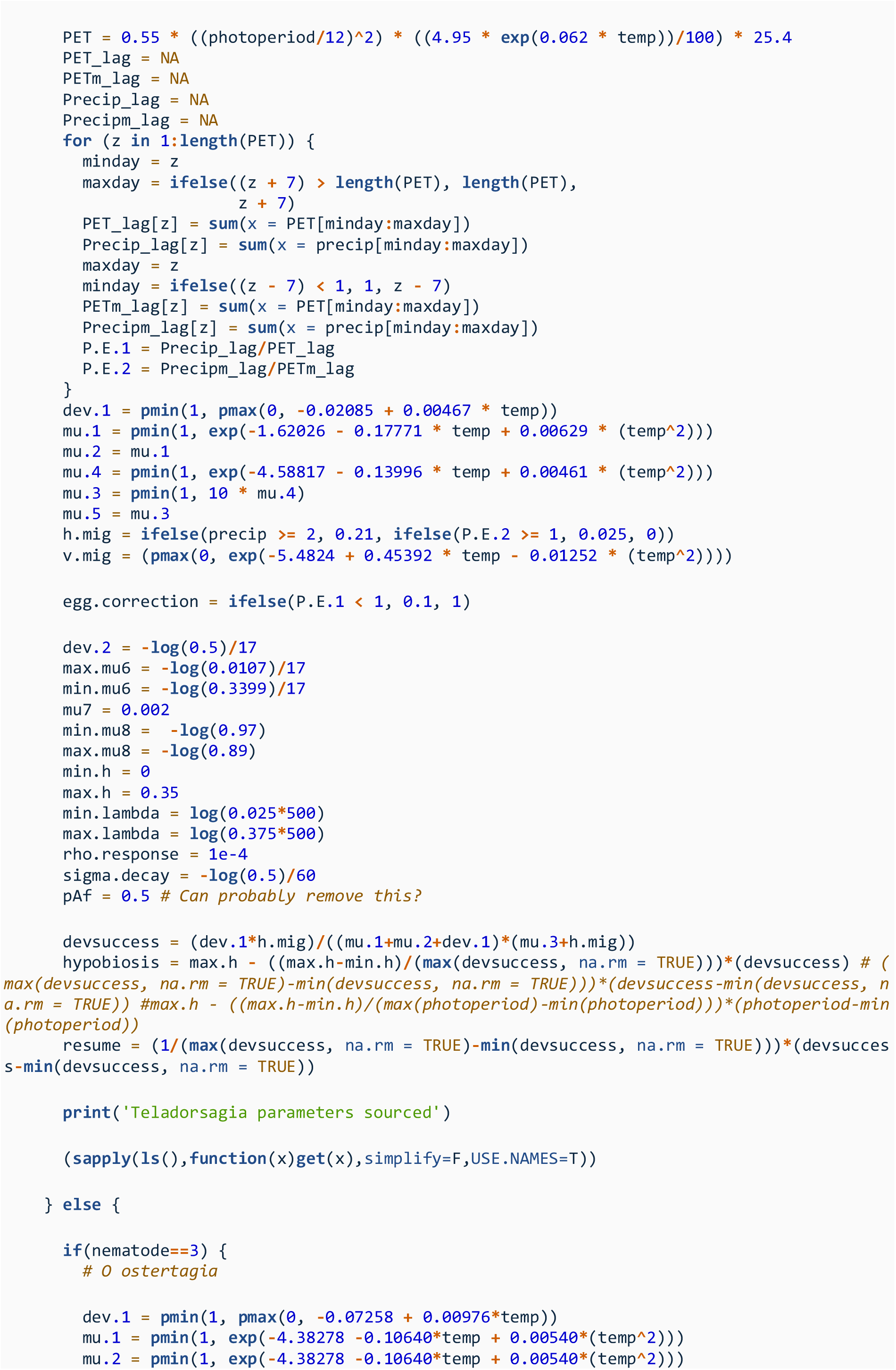
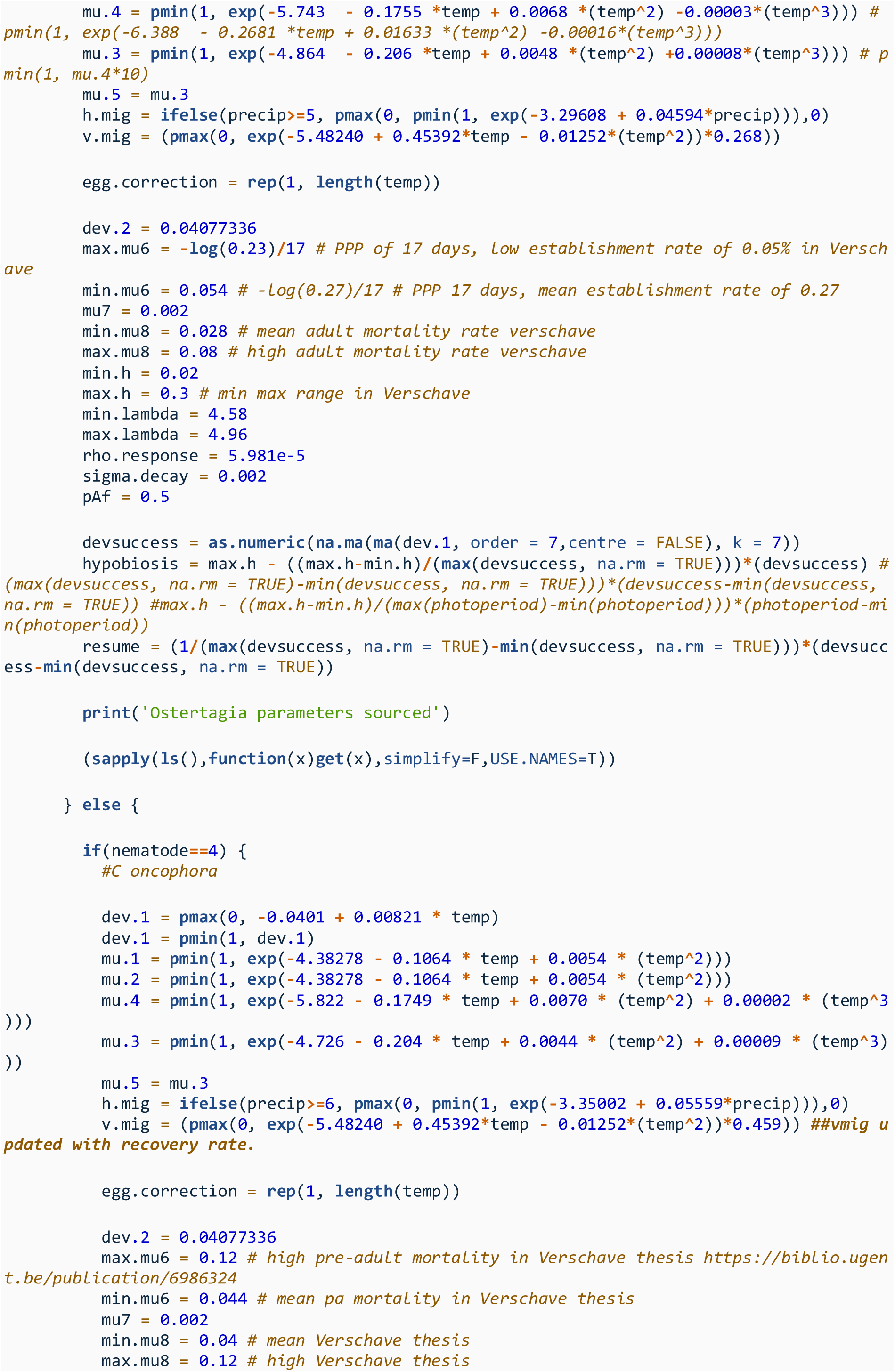
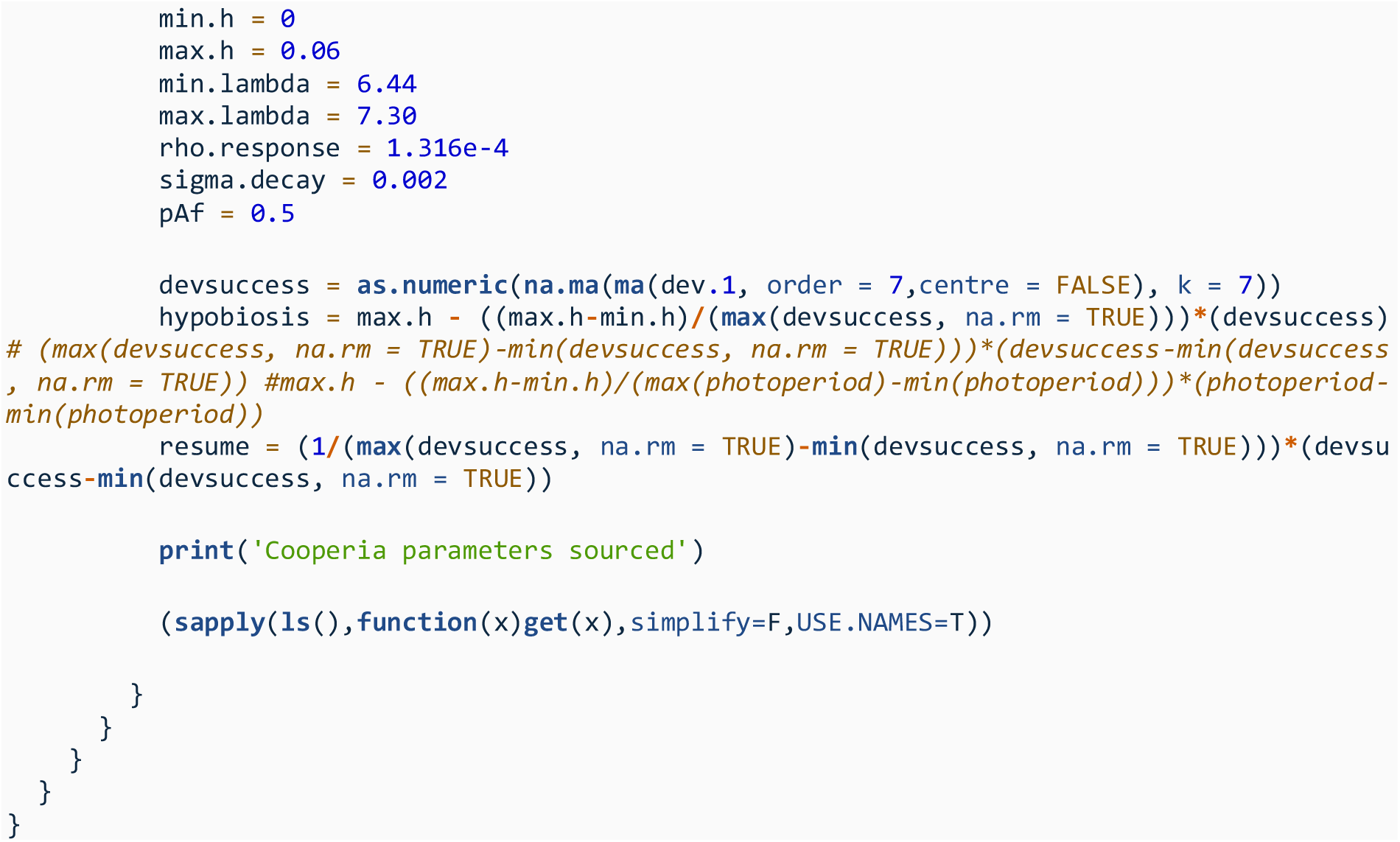

### S2 Simulation Set 1

#### S2.1 Overview

Simulation Set 1 concerned the investigation of parameters which may impact the evolutionary-epidemiological of GIN infections over time. The focus of this simulation set was to understand the factors which drive transmission and resistance.

#### S2.2 Calibration of Transmission Intensity

For simulation sets 1 (“Evolutionary epidemiological considerations of anthelmintic treatments”) and 2 (“Evaluating Anthelmintic Resistance Management Strategies”) in the main manuscript, we needed to generate simulations which covered a range of plausible transmission settings and transmission intensities. This is because there could be some prior expectation that transmission intensity may interact with both treatment efficacy (e.g., reinfection) and resistance management (e.g., the comparative size of refugia populations). It was therefore needed to explore the implication of different transmission levels.

We aimed to have two broad transmission settings, termed a “moderate” and “high” transmission setting. The aim was for the “moderate” transmission intensity simulations to have a mean (across the sampled locations) peak FEC in the second year in the absence of treatment or other interventions of ~700epg and for the “high” setting this was ~1250epg. It is important to note that because different climate settings (locations) also impact transmission through the population dynamics of the GIN populations, the actual values achieved in any an individual simulation are likely different. However, they can still be used to describe qualitatively different levels of transmission, such that for each climate setting there is a “moderate” and “high” transmission scenario. It is just the actual levels achieved may be either above or below the desired values.

In the simulation code this was set by defining the starting number of L3 free-living larvae on pasture, and was the varied until the mean peak FEC (with 100 geographic locations and climatic conditions used) in the second year hit the desired level when using different numbers of input L3. Figure S2.1 shows the result from this calibration, and we therefore used 65,000 L3p as the input for “moderate” transmission and 160,000 L3p as the input value for the “high” transmission setting.

**Figure S2.1.**
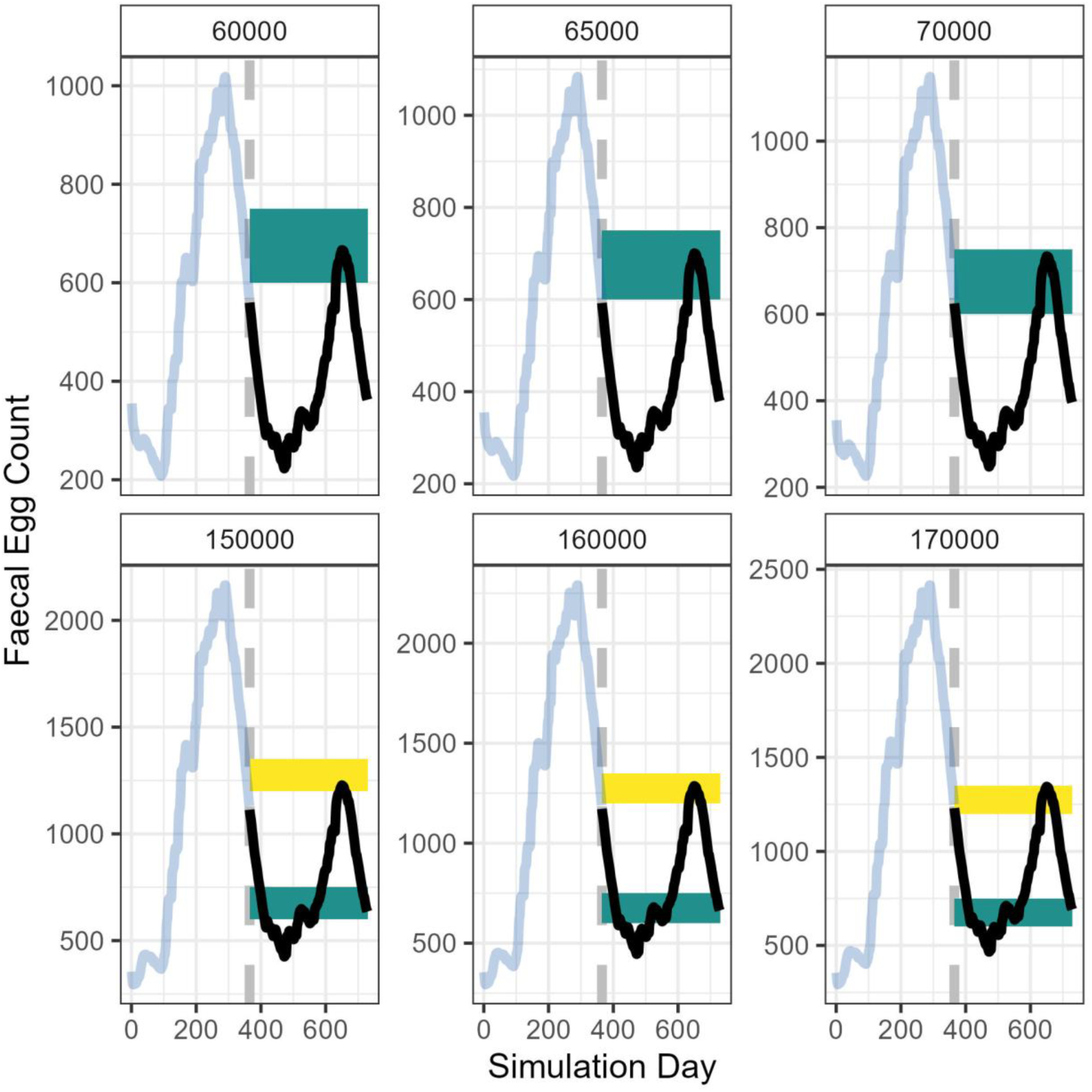
Calibration of the initial larval pasture seeding for the simulations to give required target faecal egg count for varying transmission intensity. Each panel represents the different input number of L3 pasture input into the model. The light blue line is the first year of the simulation and the black line is the second year of the simulation, where the aim is to obtain the target faecal egg count for the simulations. This plotted simulation line is the mean value of 100 geographic locations to represent a variety of climatic conditions which further impacts the dynamics of the nematode populations. The yellow rectangle indicates the range for which a “high” transmission setting would be defined, and the green rectangle for a “moderate” transmission setting.

#### S2.3 Calculating anthelmintic drug parasite killing efficacy given drug decay

In the simulations where drug decay was included there was a need to include a drug efficacy decay curve. Given that different drugs are given at different concentrations, and the drug concentration to parasite killing efficacy will vary between drug formulation, rather than tracking drug decay as the amount of the physical drug itself, the efficacy of the drug itself was used. This allows for a far more generalisable system, and further allows for an agnostic approach to precise drug formulations. While there are many potential ways to include drug decay, we consider a simplistic method of exponential decay:

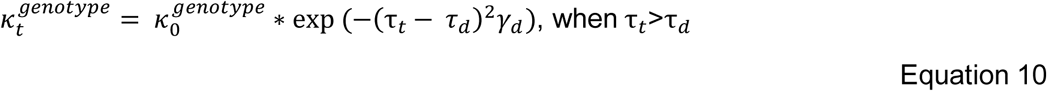

Where 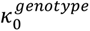 is the killing efficacy of the drug of the specified genotype when at the target drug concentration in the livestock, that is the registered therapeutic dose. *τ_t_* is the number of days since the anthelmintic drug was administered. *τ_d_* is the number of days after administration the drug begins to decay. The value *γ_d_* is the decay rate, which is 0.005 for simulations with decay, and 1 for simulations without decay, which can be used to give the drug efficacy and decay profiles given in Figure S2.2, and are those used in simulation set 1.

#### S2.4 Conditions varied for sensitivity analysis

**Table S2.1.**
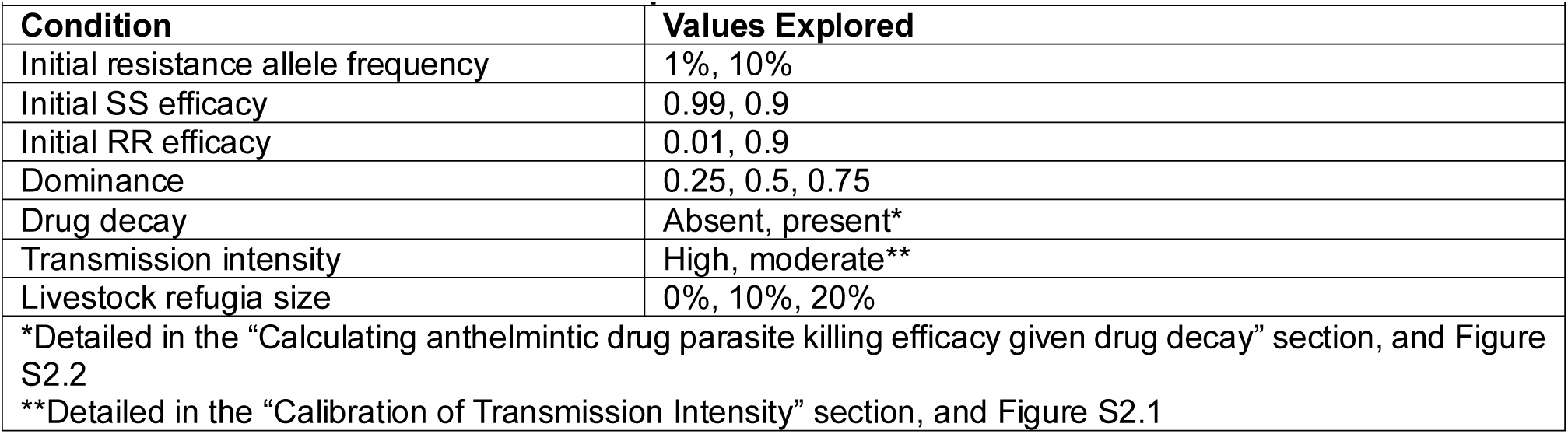
Parameters and conditions explored in Simulation Set 1.

| Table S2.1 Parameters and conditions explored in Simulation Set 1. |  |
| --- | --- |
| Condition | Values Explored |
| Initial resistance allele frequency | 1%, 10% |
| Initial SS efficacy | 0.99, 0.9 |
| Initial RR efficacy | 0.01, 0.9 |
| Dominance | 0.25, 0.5, 0.75 |
| Drug decay | Absent, present* |
| Transmission intensity | High, moderate** |
| Livestock refugia size | 0%, 10%, 20% |
| *Detailed in the “Calculating anthelmintic drug parasite killing efficacy given drug decay” section, and Figure S2.2 |  |
| **Detailed in the “Calibration of Transmission Intensity” section, and Figure S2.1 |  |

#### S2.5 Supplementary Analysis

Given the wide range of parameter values explored (288 total permutations, Table S2.1), we only presented a subset of this data in the main manuscript. In this supplementary analysis, we further consider all the permutations of parameters investigated. But note the main qualitative conclusions identified in the main manuscript are unchanged in this further exploration. Nevertheless, given the current uncertainty with how different parameter values may interact, it was still prudent to explore these additional permutations, and therefore provides further robustness of the presented conclusions.

**Figure S2.3.**
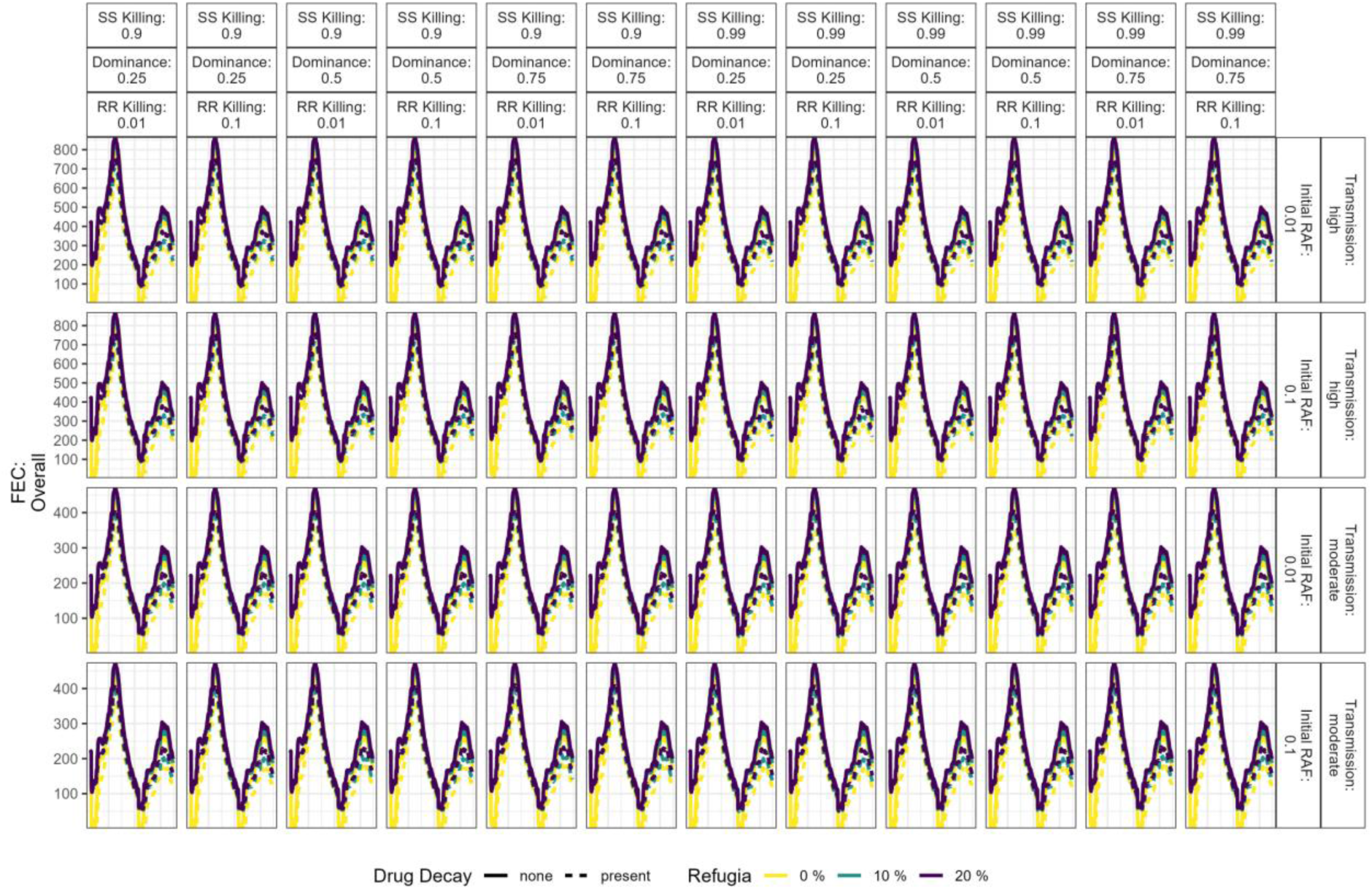
Full results for Simulation Set 1 Overall Faecal Egg Count, including all input parameter permutations. The plot is stratified left-right by the resistance phenotypes of the SS, RS and RR genotypes; where the RS phenotype is dependent on the dominance of resistance. The plot is stratified top-bottom by the overall level of transmission and the initial resistance allele frequency (RAF) at the start of the simulation. The outcome (y axis) is the overall faecal egg count for all livestock (cohort 1 and cohort 2).

**Figure S2.4.**
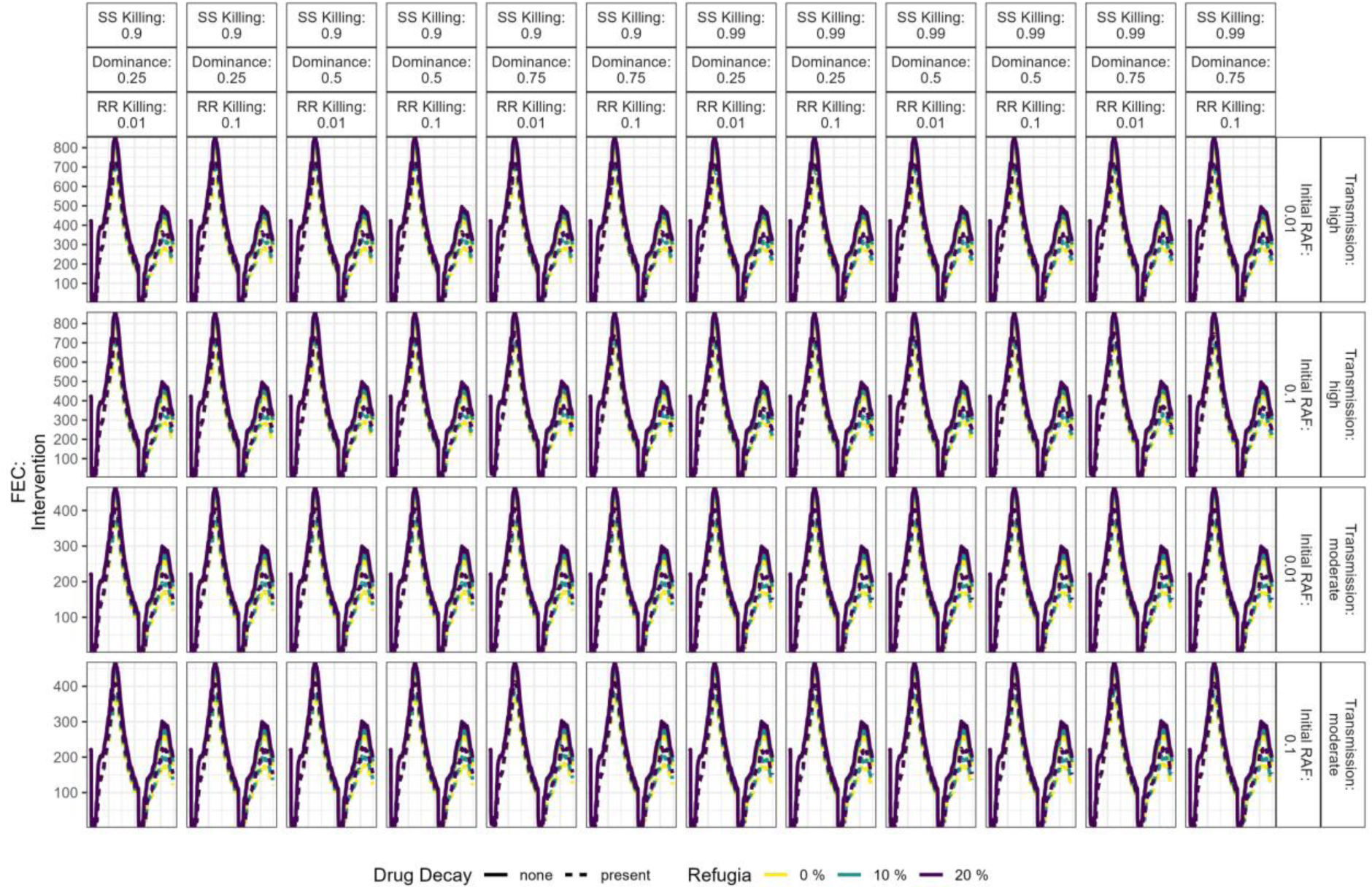
Full results for Simulation Set 1 Faecal Egg Count for Intervention Livestock, including all input parameter permutations. The plot is stratified left-right by the resistance phenotypes of the SS, RS and RR genotypes; where the RS phenotype is dependent on the dominance of resistance. The plot is stratified top-bottom by the overall level of transmission and the initial resistance allele frequency (RAF) at the start of the simulation. The outcome (y axis) is the overall faecal egg count for only the treated “intervention” livestock (cohort 1).

**Figure S2.5.**
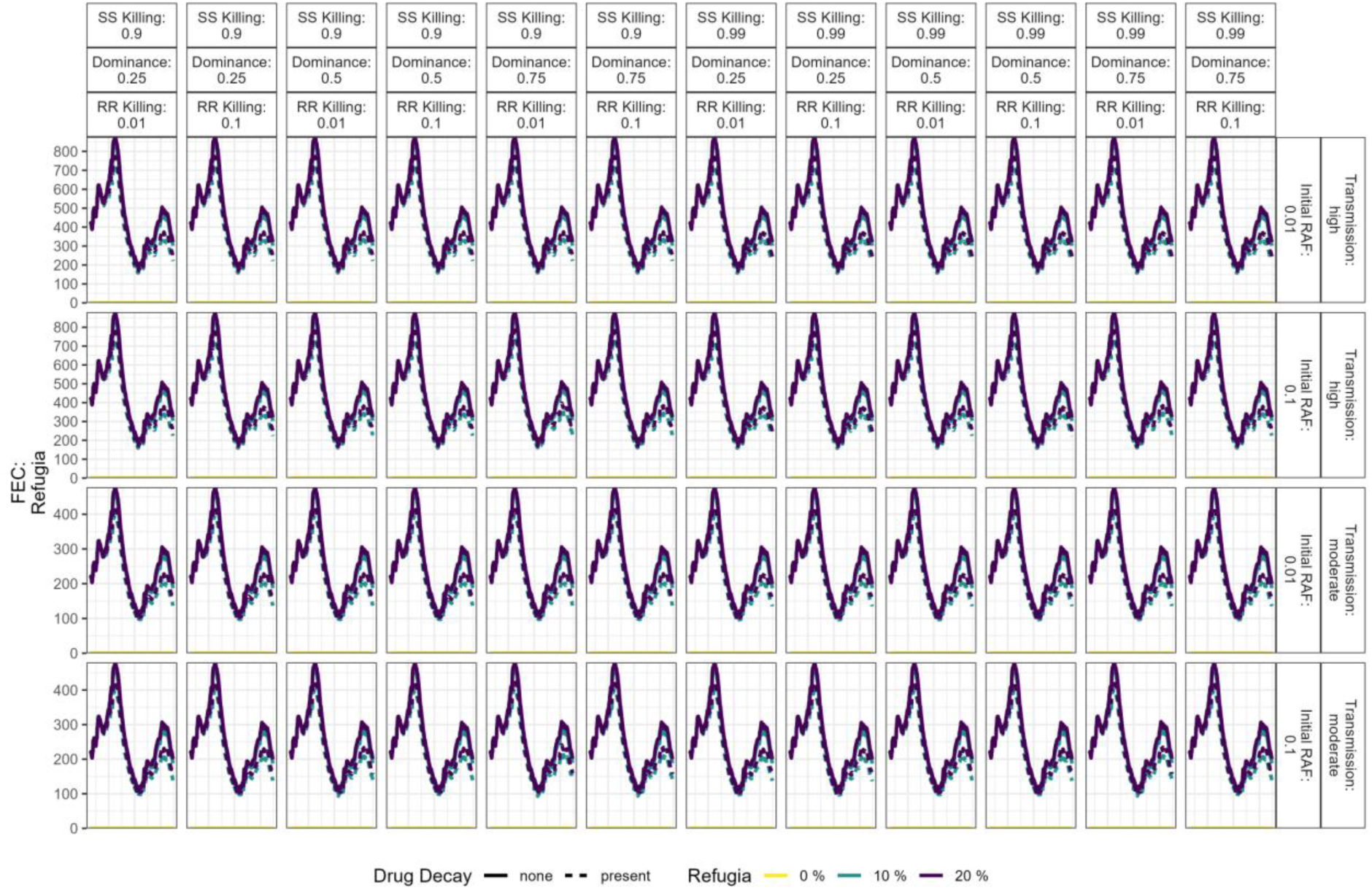
Full results for Simulation Set 1 Faecal Egg Count for Refugia Livestock, including all input parameter permutations. The plot is stratified left-right by the resistance phenotypes of the SS, RS and RR genotypes; where the RS phenotype is dependent on the dominance of resistance. The plot is stratified top-bottom by the overall level of transmission and the initial resistance allele frequency (RAF) at the start of the simulation. The outcome (y axis) is the overall faecal egg count for only the untreated “refugia” livestock (cohort 2).

**Figure S2.6.**
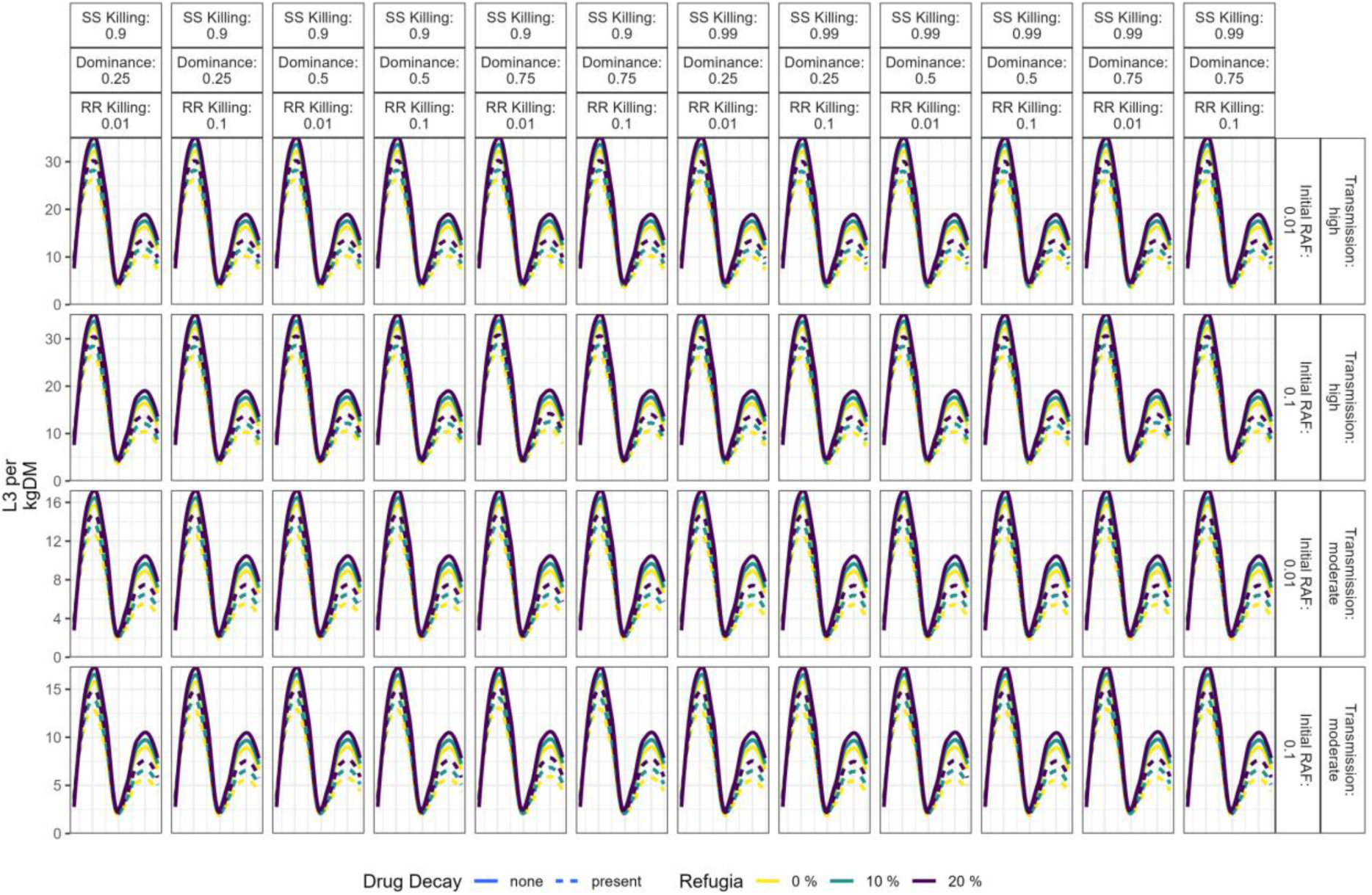
Full results for Simulation Set 1 pasture larval counts, including all input parameter permutations. The plot is stratified left-right by the resistance phenotypes of the SS, RS and RR genotypes; where the RS phenotype is dependent on the dominance of resistance. The plot is stratified top-bottom by the overall level of transmission and the initial resistance allele frequency (RAF) at the start of the simulation. The outcome (y axis) is the L3 per kg of dry matter, which can be considered a measure of transmission intensity.

**Figure S2.7.**
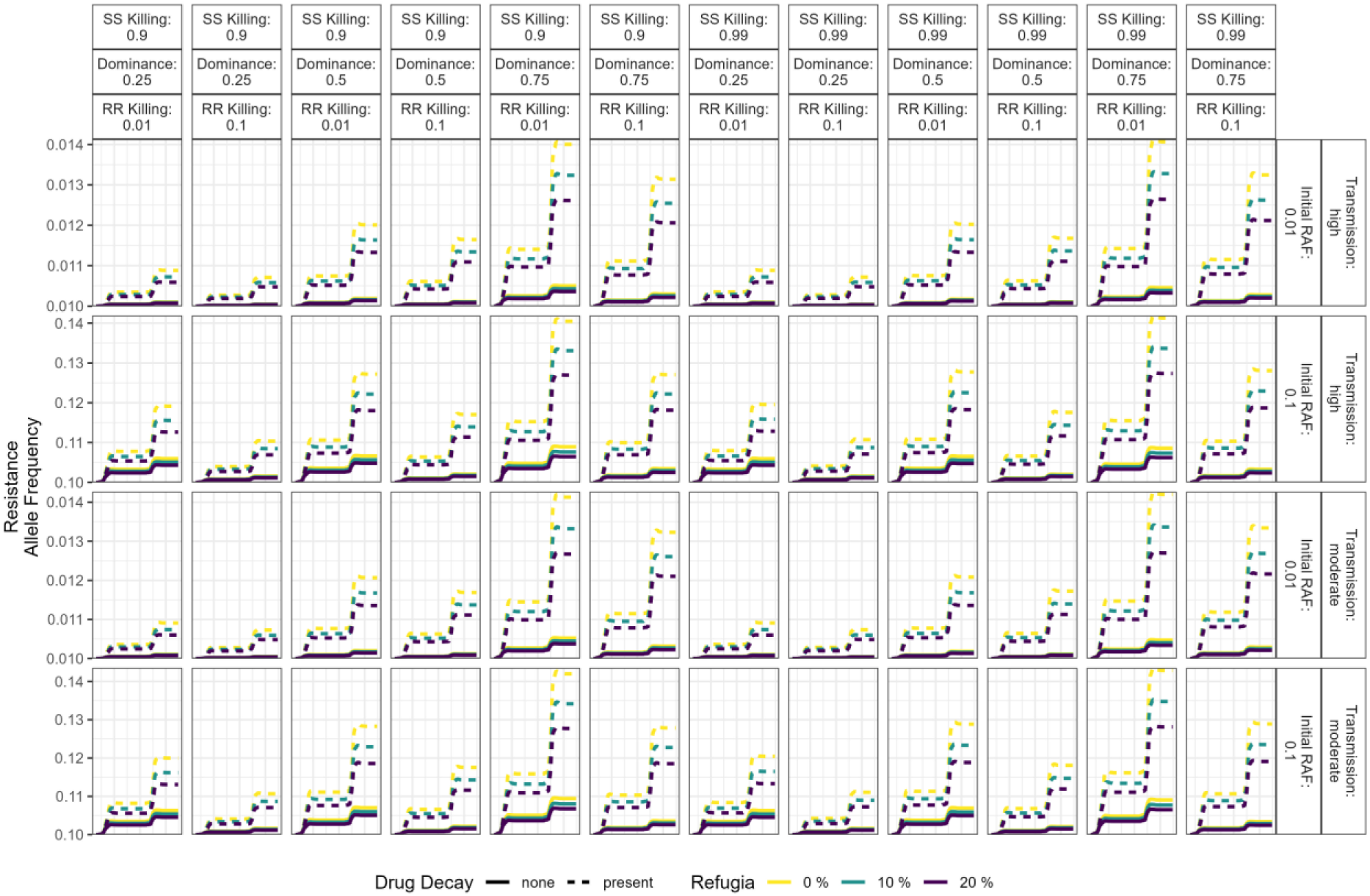
Full results for Simulation Set 1 Resistance Allele Frequency, including all input parameter permutations. The plot is stratified left-right by the resistance phenotypes of the SS, RS and RR genotypes; where the RS phenotype is dependent on the dominance of resistance. The plot is stratified top-bottom by the overall level of transmission and the initial resistance allele frequency (RAF) at the start of the simulation. The outcome (y axis) is the resistance allele frequency, which can be considered a measure of resistance management.

**Figure S2.8.**
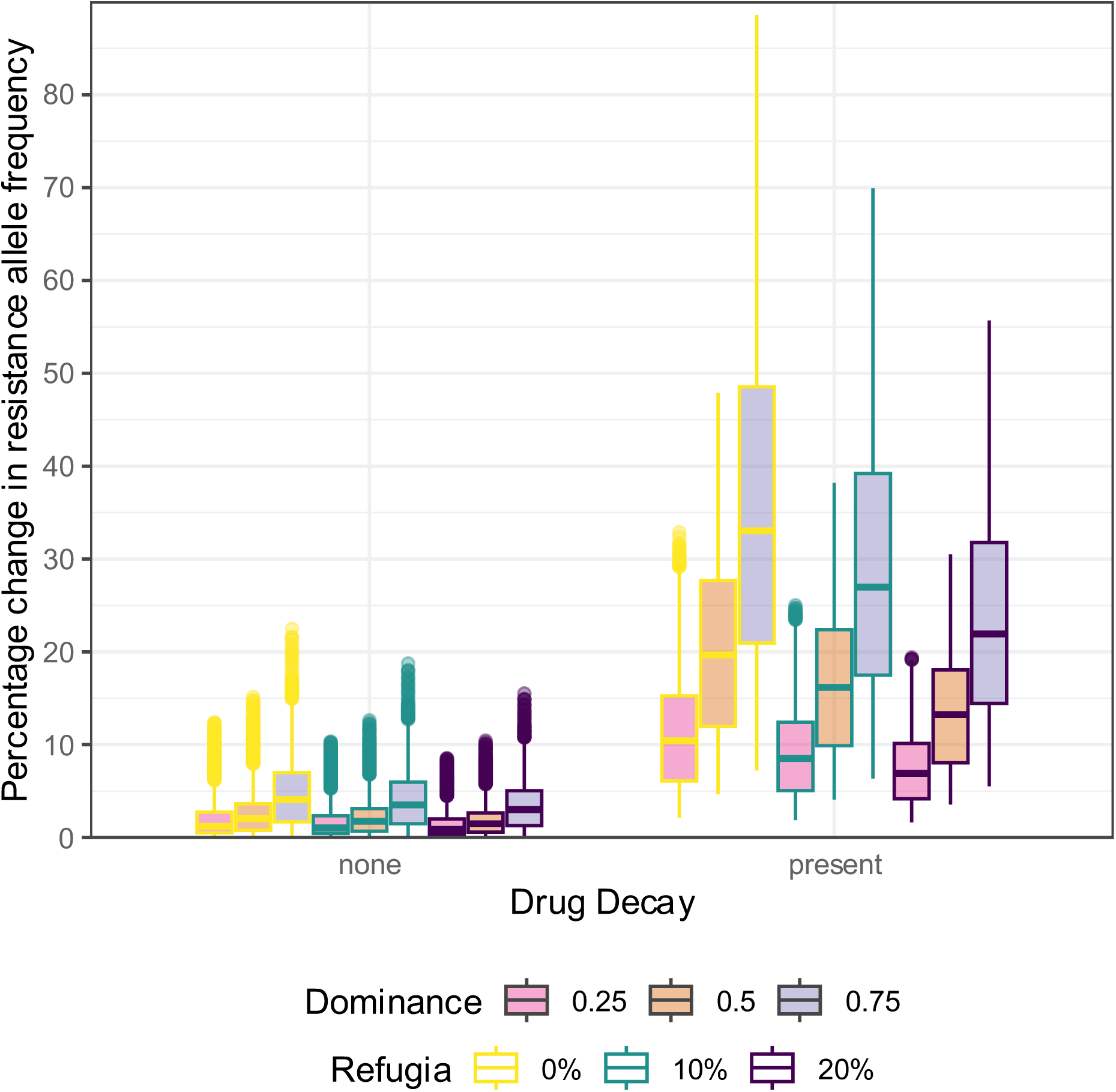
Percentage change in the resistance allele frequency at the end of the simulation (day = 1086).

### S3 Simulation Set 2

#### S3.1 Overview

Simulation set 2 (“Evaluating Anthelmintic Resistance Management Strategies”) involved to direct comparison of different ARM strategies, with the aim to identify the broad level performance of ARM strategies. It should be noted that there is much uncertainty with modelling resistance management (Hastings, 2001) and so only qualitative inferences should be made. Modelling can be used to explore otherwise complicated strategies (e.g., cohort mosaics), where a theoretical advantage would be needed to justify further (expensive and time consuming) empirical investigation.

#### S3.2 Detailed Parameter Descriptions

**Fig S3.1:**
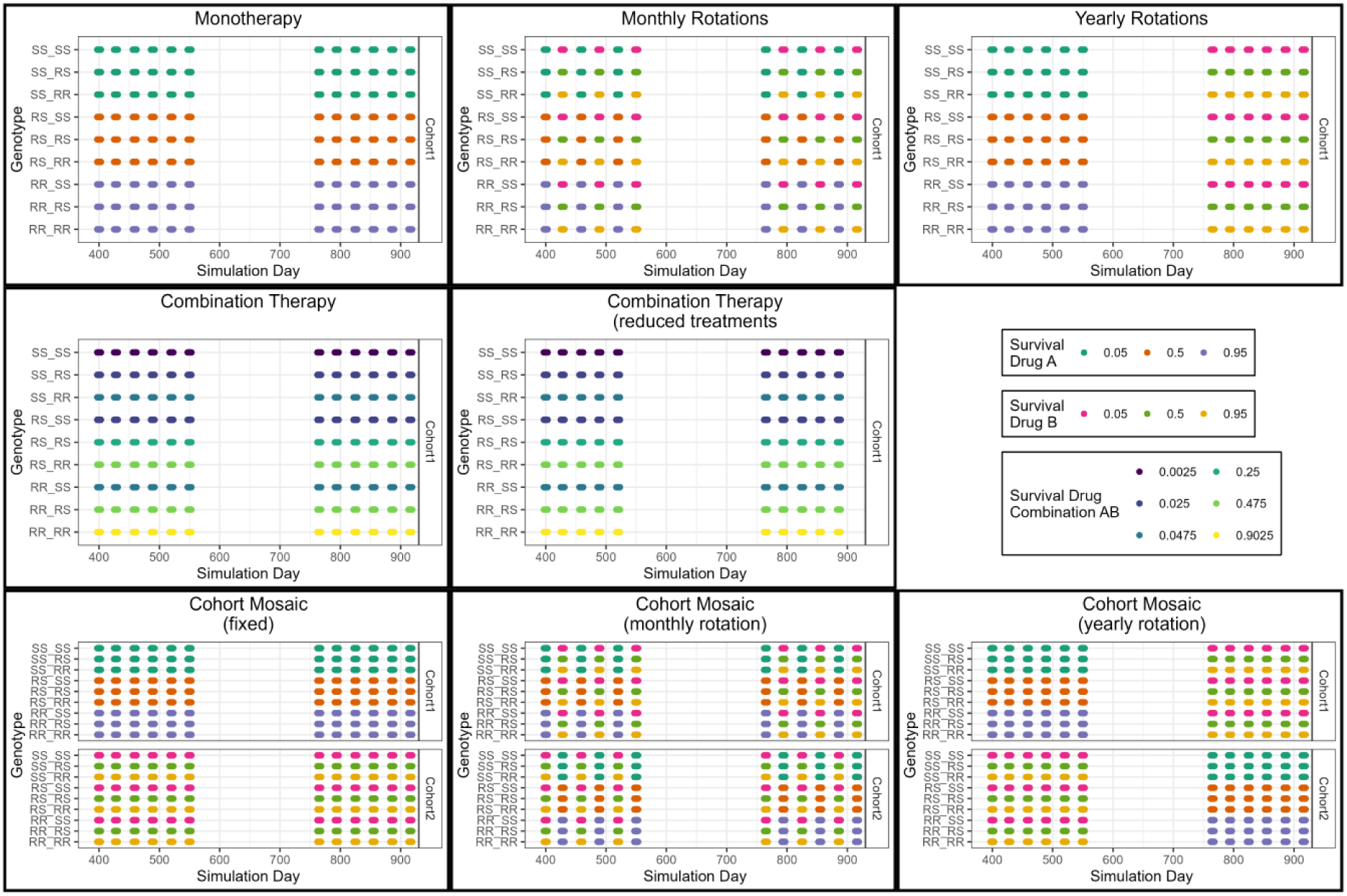
Conceptualisation of the ARM strategies with survival probabilities. Each panel shows the survival for the genotypes at the time points when given the designated drug treatments. The colours indicate the survival probability for the genotype at the specified time to the anthelmintic drug treatment which will be given (as detailed in the figure legend panel).

#### S3.3 Conditions varied for sensitivity analysis

**Table S2.1.** Parameters and conditions explored in Simulation Set 2.

| Condition | Values Explored |
| --- | --- |
| Initial resistance allele frequency: Drug 1 | 1%, 10% |
| Initial resistance allele frequency: Drug 2 | 0.99, 0.9 |
| Strategy | Monotherapy, Monthly pharmacological rotation, Yearly pharmacological rotation, Fixed cohort mosaic, Monthly cohort mosaic, Yearly cohort mosaic, Combinations, Combinations with reduced coverage, Combinations with reduced frequency |

**Table S2.1.** Parameters fixed in Simulation Set 2.

| Table S2.1 Parameters fixed in Simulation Set 2. |  |
| --- | --- |
| Condition | Values Explored |
| Parasite killing efficacy of each drug against homozygous susceptible | 95% |
| Parasite killing efficacy of each drug against homozygous resistance | 5% |
| Dominance of Resistance | 0.5 |
| Transmission intensity |  |
| Refugia Size | 20% (30% in Combinations with reduced coverage) |
| Number of treatments per year | 6 (5 in Combinations with reduced frequency) |

#### S3.4 Supplementary Analysis

Figure S3.2 shows the absolute faecal egg counts for the livestock populations. A clear indication here is that, regardless of the level of resistance or ARM strategy, the overall epidemiological difference is small. However these differences do become larger when there are higher levels of resistance.

**Figure S3.2.**
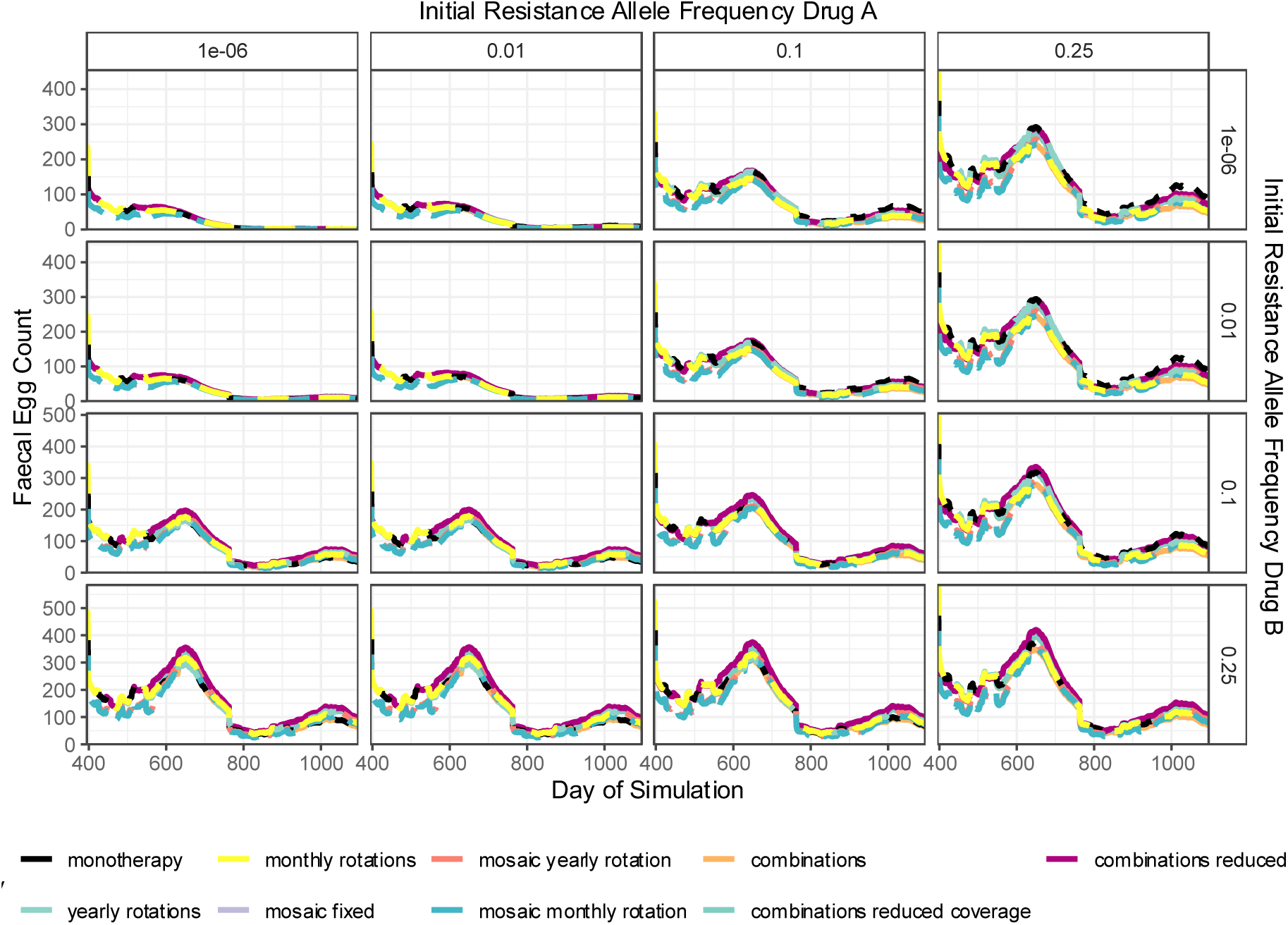
Simulation Set 2 – Faecal Egg Count in Response to Anthelmintic Drug Treatments. The y axis is the faecal egg count. The long-dash lines indicate a rotation strategy, short-dash lines a cohort mosaic strategy, and solid lines indicate a combination therapy strategy. The colour of the lines indicates the specific strategy. The x-axis is the simulation day and is restricted to not include the 1-year burn-in which is the same across all strategies. Each line is the mean of 100 simulations, covering a range of geographic locations.

**Figure S3.3.**
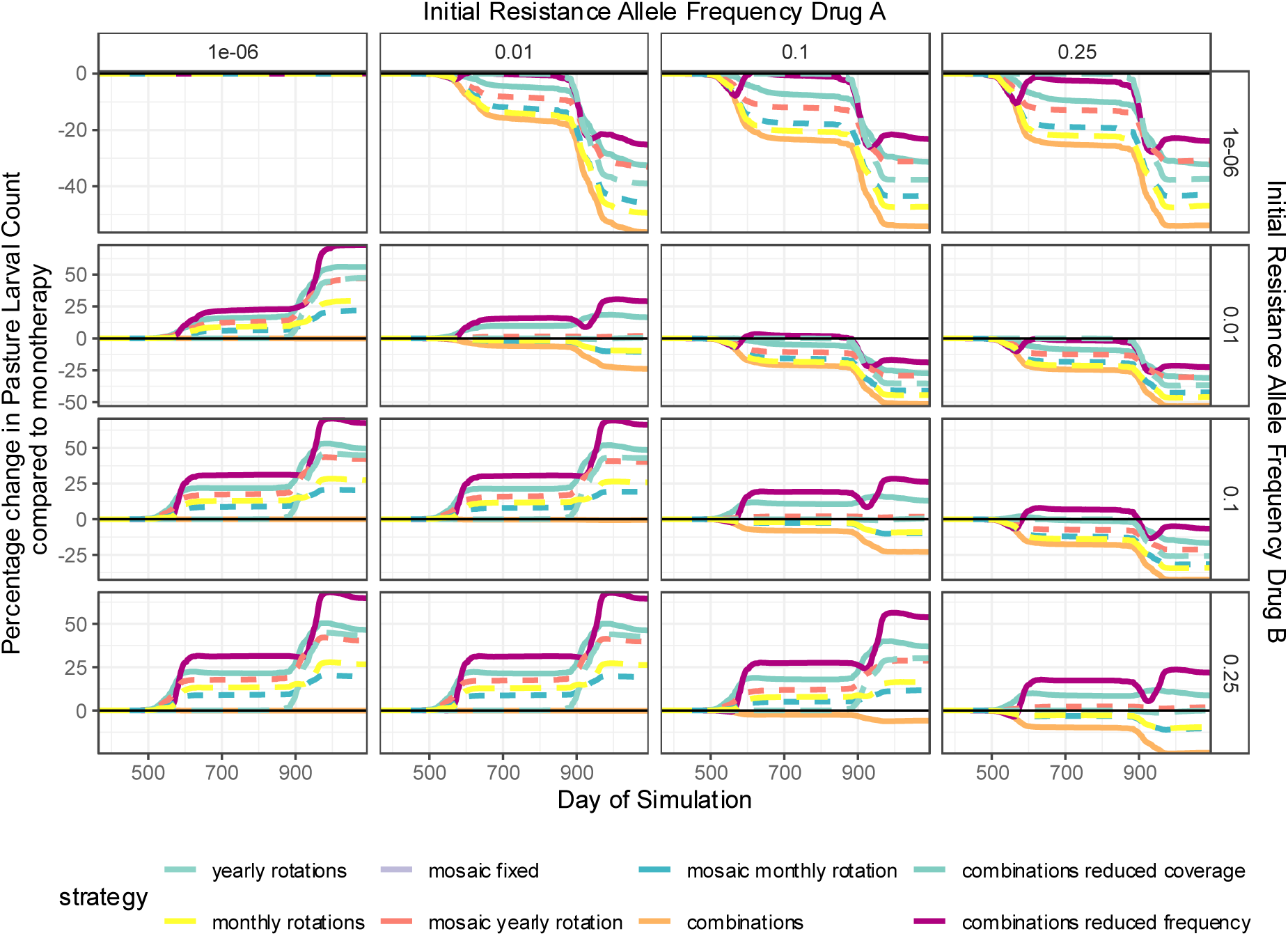
Simulation Set 2: Percentage change in Larval Pasture Counts. The y axis is the percentage change in the larval pasture count, as calculated using Equation 1 in the main manuscript. The horizontal black line (at zero) represents the no difference between strategies. Values below the black line indicates the proposed strategy performed better, and values above performed worse than the Monotherapy comparator. The long-dash lines indicate a rotation strategy, short-dash lines a cohort mosaic strategy, and solid lines indicate a combination therapy strategy. The colour of the lines indicates the specific strategy. The x-axis is the simulation day and is restricted to not include the 1-year burn-in which is the same across all strategies. Each line is the mean of 100 simulations, covering a range of geographic locations.

**Figure S3.4.**
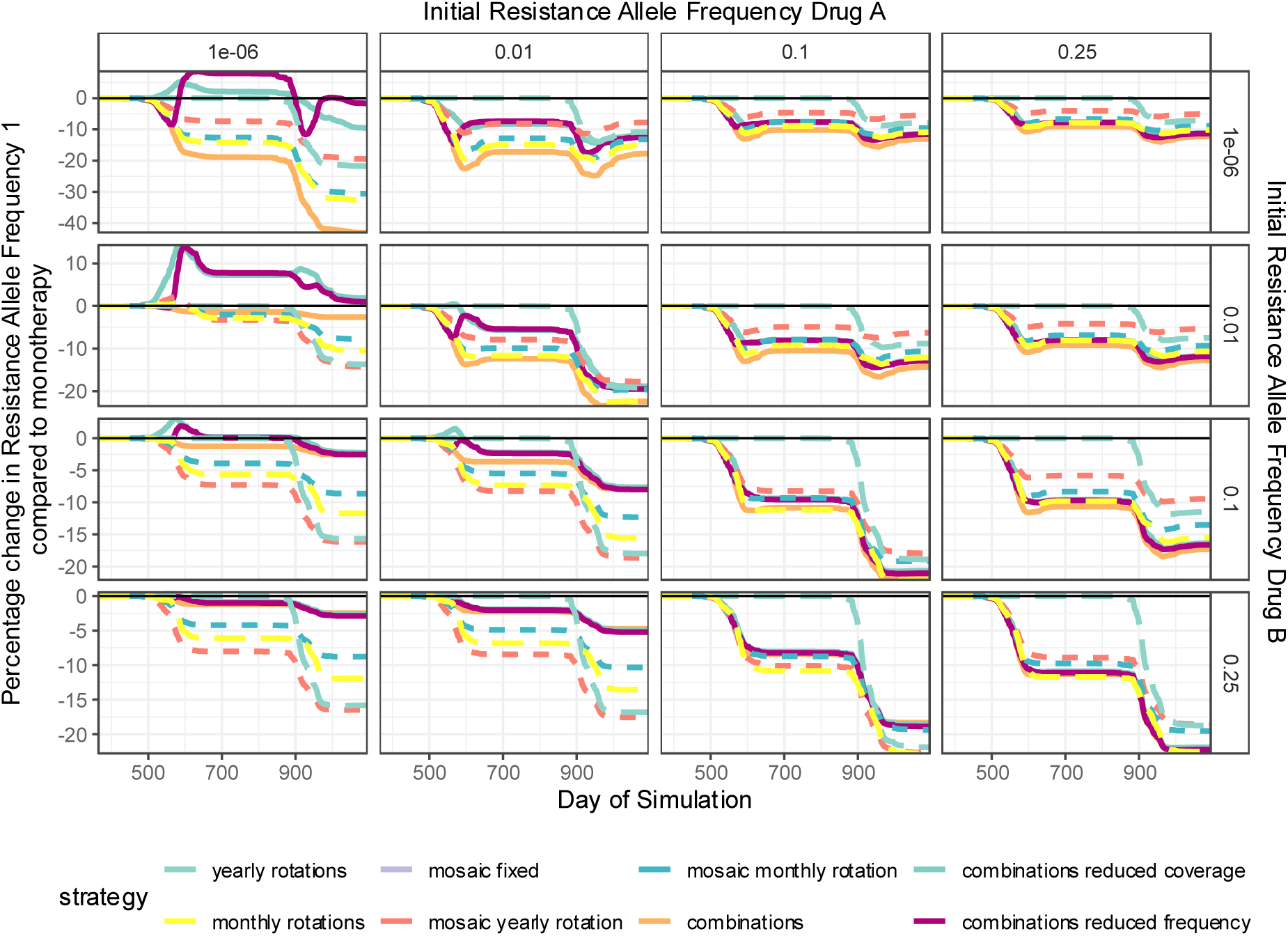
Percentage change in the resistance allele frequency to Drug 1. The y axis is the percentage change in the resistance allele frequency for Drug 1, as calculated using Equation 1 in the main manuscript. Note, Drug 1 is the drug used in the monotherapy simulations and Drug 2 is not deployed in the monotherapy simulations. The horizontal black line (at zero) represents the no difference between strategies. Values below the black line indicates the proposed strategy performed better, and values above performed worse than the Monotherapy comparator. The long-dash lines indicate a rotation strategy, short-dash lines a cohort mosaic strategy, and solid lines indicate a combination therapy strategy. The colour of the lines indicates the specific strategy. The x-axis is the simulation day and is restricted to not include the 1-year burn-in which is the same across all strategies. Each line is the mean of 100 simulations, covering a range of geographic locations.

**Figure S3.5.**
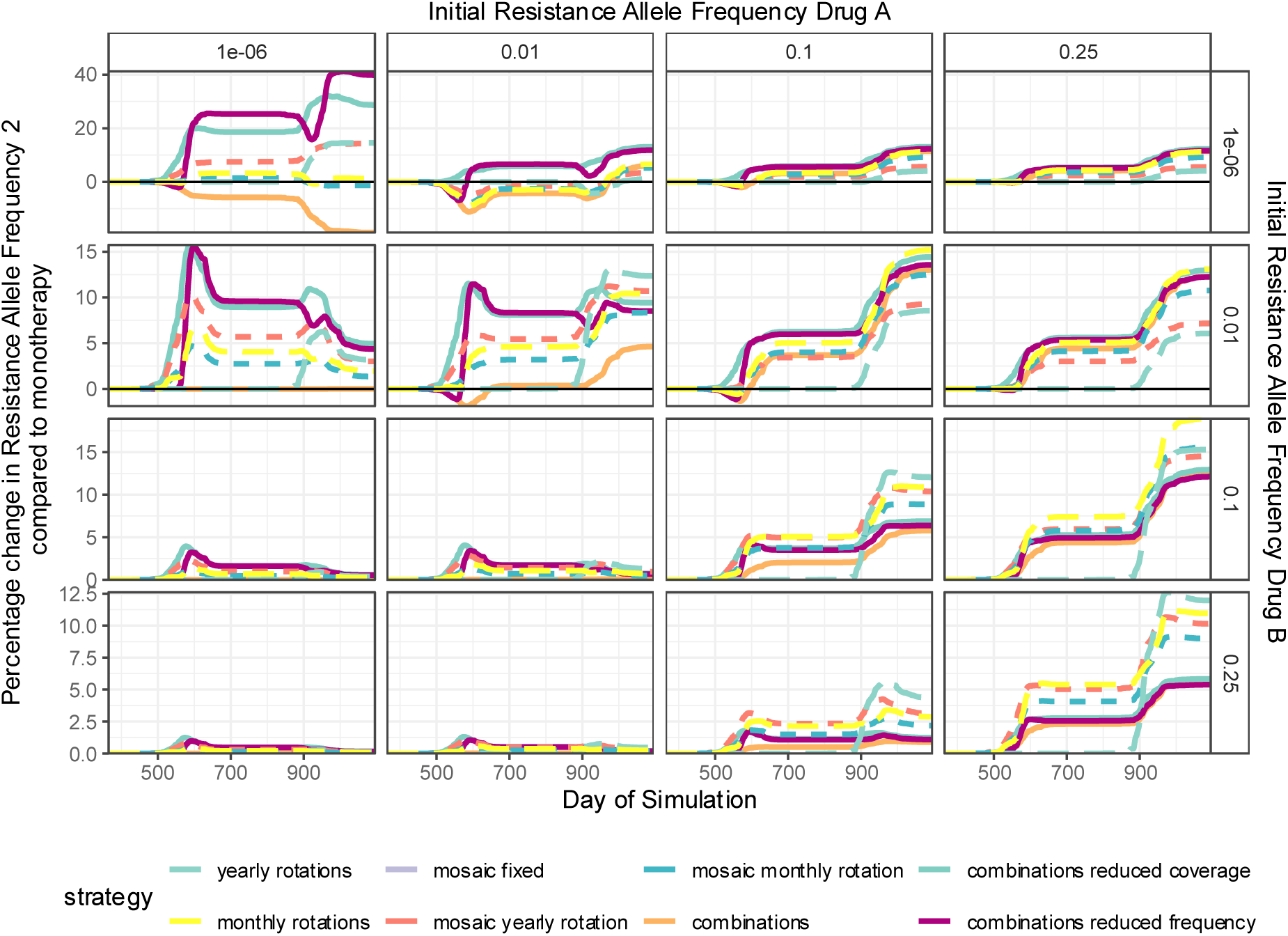
Percentage change in the resistance allele frequency to Drug 2. The y axis is the percentage change in the resistance allele frequency for Drug 2, as calculated using Equation 1 in the main manuscript. Note, Drug 1 is the drug used in the monotherapy simulations, and Drug 2 is not deployed in the monotherapy simulations. The horizontal black line (at zero) represents the no difference between strategies. Values below the black line indicates the proposed strategy performed better, and values above performed worse than the Monotherapy comparator. The long-dash lines indicate a rotation strategy, short-dash lines a cohort mosaic strategy, and solid lines indicate a combination therapy strategy. The colour of the lines indicates the specific strategy. The x-axis is the simulation day and is restricted to not include the 1-year burn-in which is the same across all strategies. Each line is the mean of 100 simulations, covering a range of geographic locations.

### S4 GLOWORM Simulation Framework Showcase

#### S4.1 Overview

Given the increase in the complexity of the GLOWORM simulation framework, it becomes beneficial to display the technical ability of the current simulation; especially as a proof of principal of what the simulation framework can be used to for. These simulations are to present only the technical simulation capability, and ***<u>conclusions regarding anthelmintic</u> <u>deployment practices must not be drawn from these presented examples</u>***. The simulations in these examples deliberately use arbitrary values, which we intentionally do not report, to showcase the model features. Each example simulation is presented to showcase the model capability, and should not be used to interpret the effectiveness of any worm control practices.

#### S4.2 Example Simulations

**Figure S4.1.**
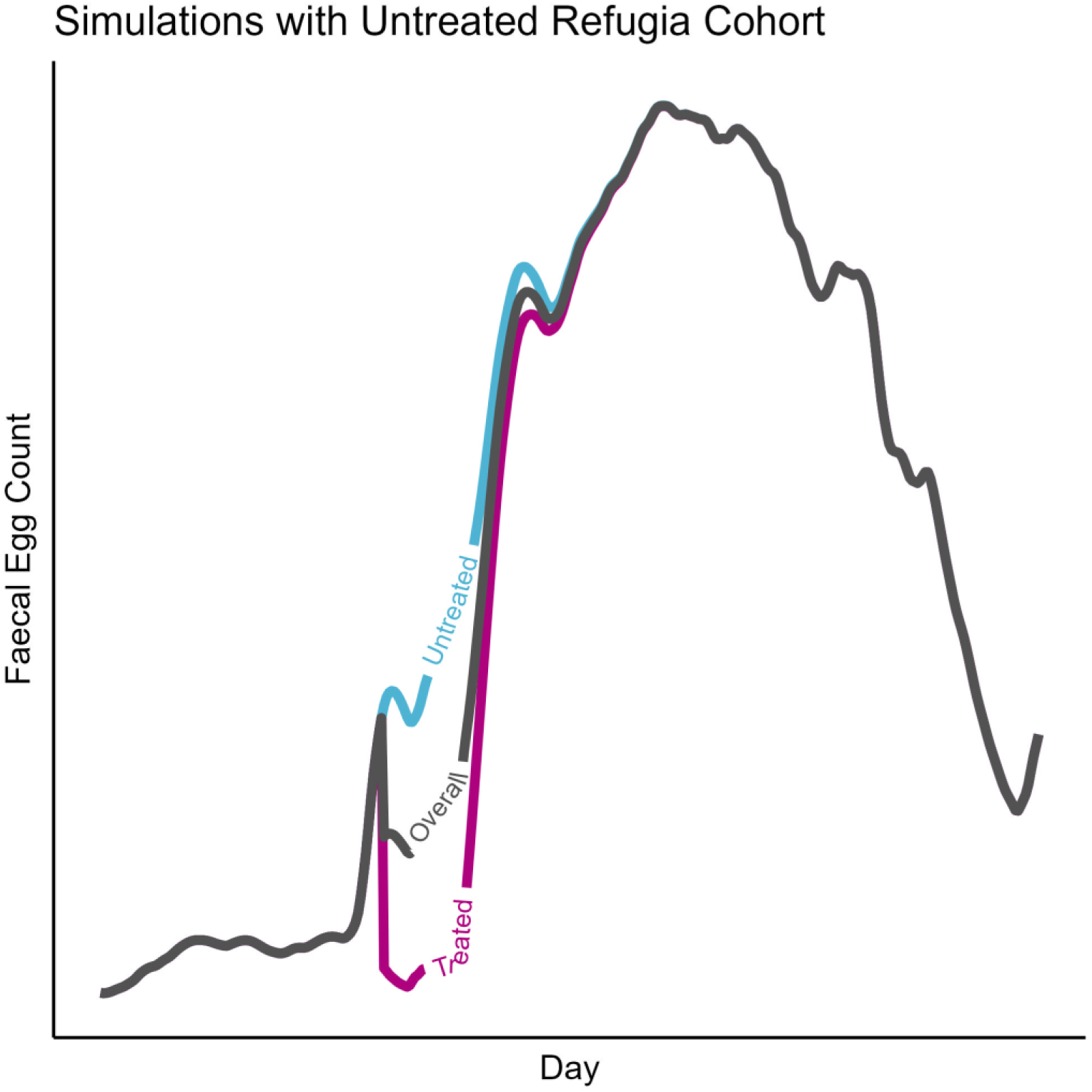
Untreated Livestock Refugia. Inclusion of explicit refugia populations. As each cohort in the model simulations can be given unique treatments, the inclusion of an explicitly modelled untreated livestock refugia can be included. The use case for this will be for evaluating how the relative size of refugia populations influences resistance frequency, but also understanding the disease control implications of leaving portions of the livestock population untreated.

**Figure S4.2.**
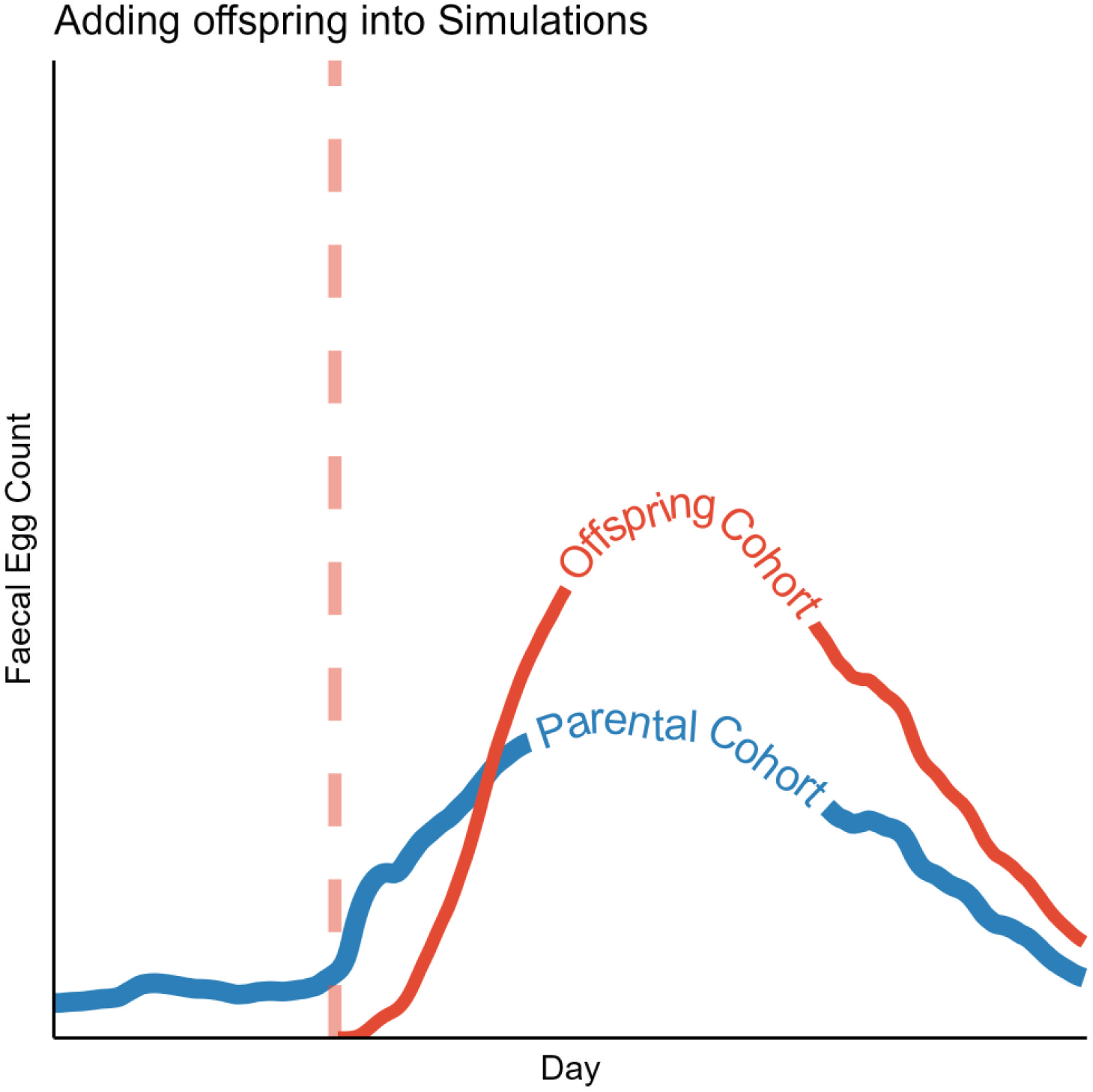
Adding in Parent-Offspring Livestock into the Simulations. The simulations can used to add in a new “cohort” which is the offspring of another “parental” cohort of in the simulations. The applicability of this can therefore be to look at the implication of the periparturient rise in faecal egg count which is observed to happen to parental ewes at lambing time. The offspring cohort can then have a different weight, which then relates to dry matter intake and faecal production.

**Fig S4.3.**
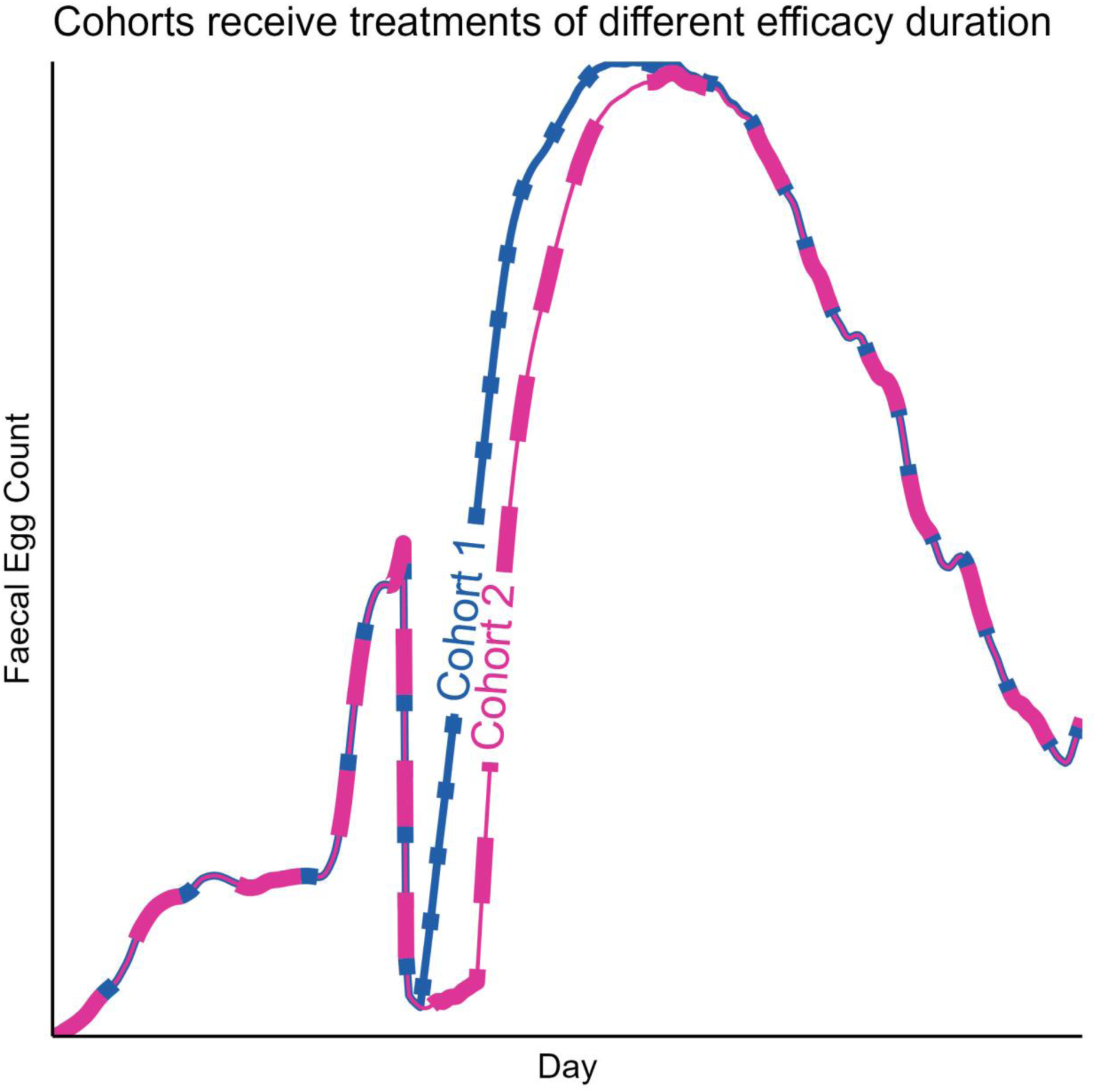
Livestock cohorts receive the same treatment, but has different efficacy durations. Each cohort can be given the same drug formulation, however to account for the heterogeneity in drug efficacies between livestock the drugs can be given different efficacy profiles for each cohort. The use case here is therefore to explore how treatment heterogeneities impact the spread of disease and resistance.

**Figure S4.4.**
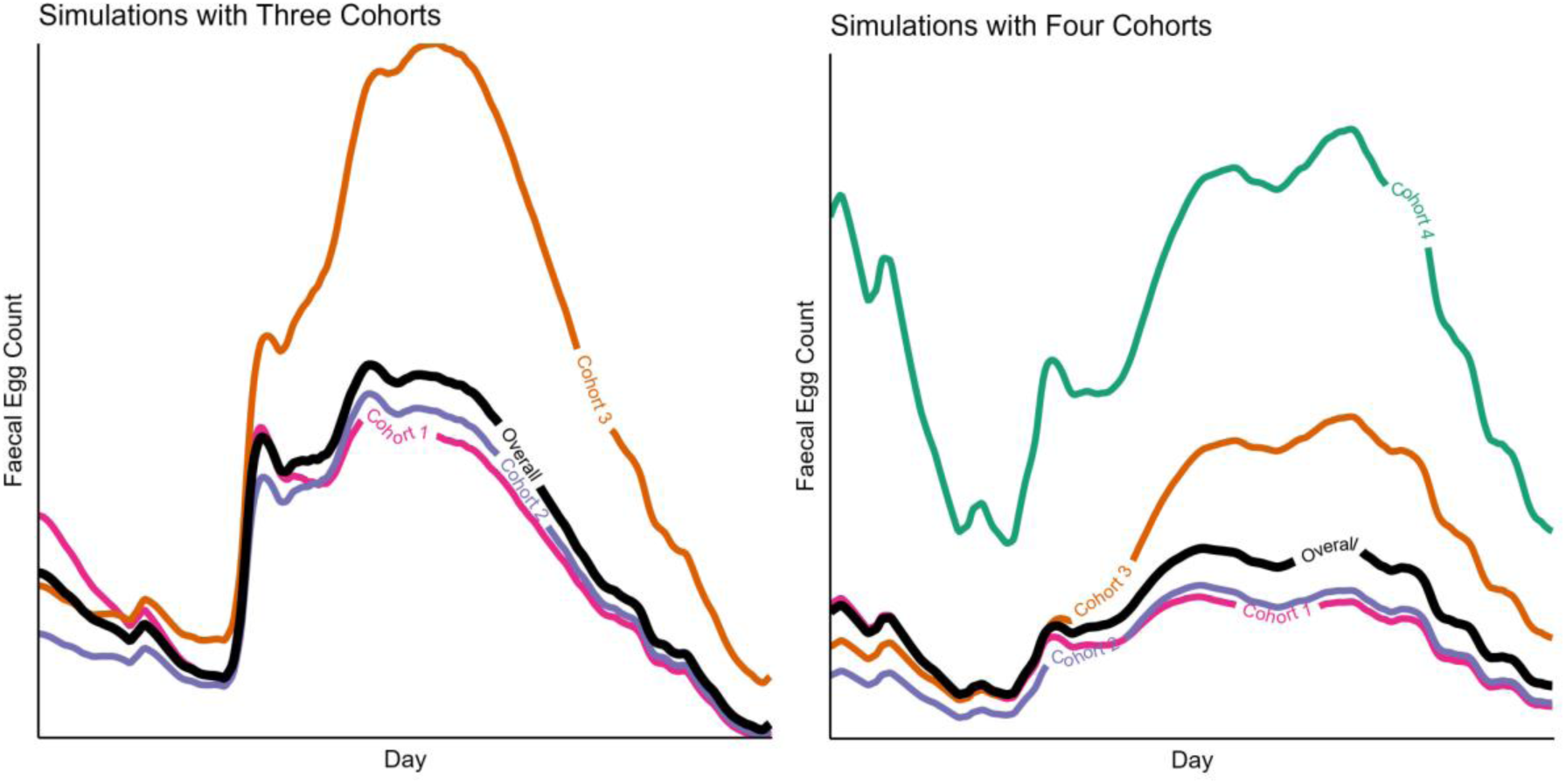
Simulations with multiple livestock cohorts. Simulations can be conducted with up to four separate livestock cohorts. Who can each be given unique ages (bodyweights) and dry matter intakes. The panel on the left shows a simulation with three cohorts, and the panel on the right shows the simulation with four cohorts.

**Figure S4.5.**
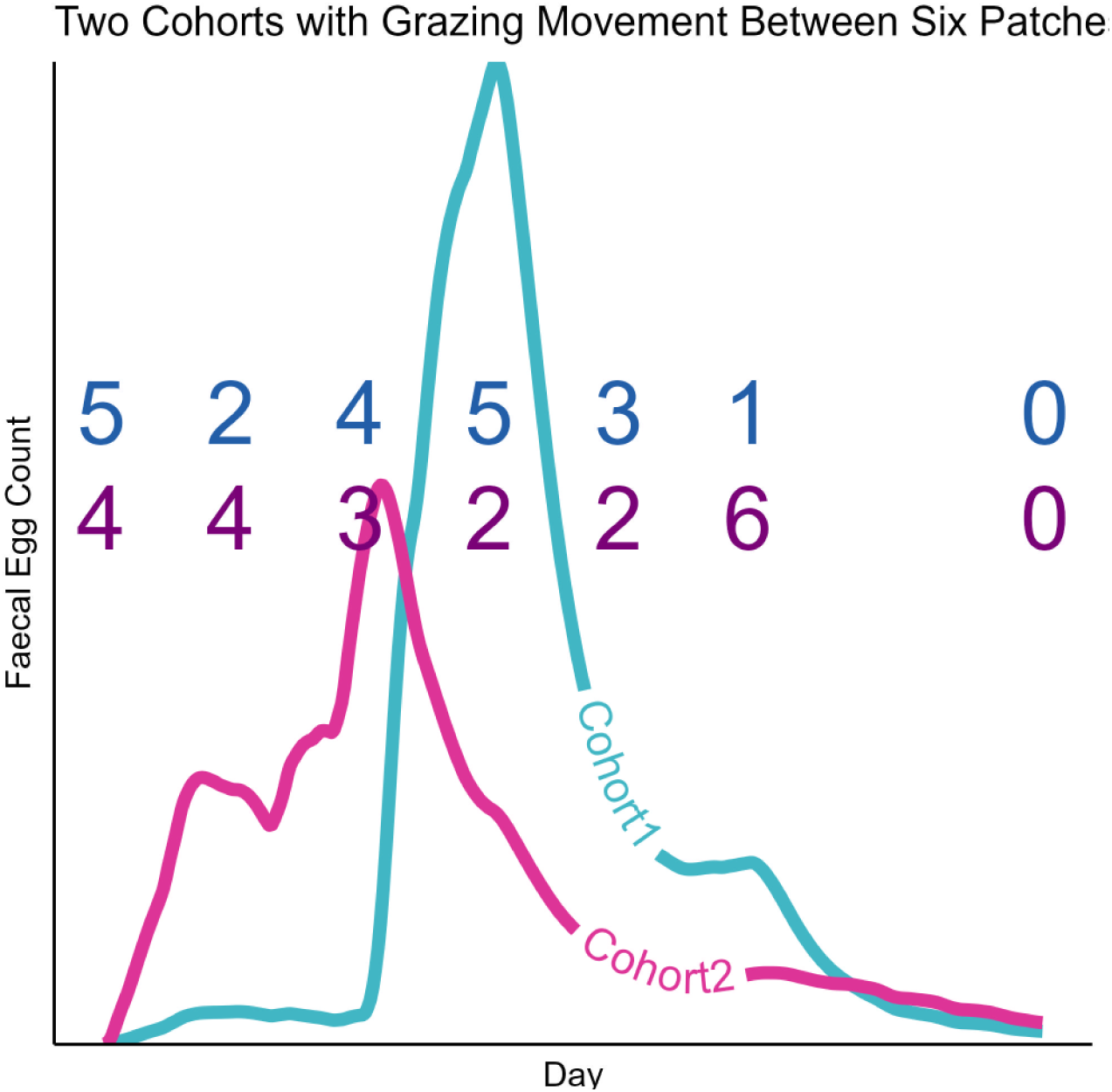
Two Livestock Cohorts with Different Rotational Grazing Movements. Each cohort can also be assigned a unique pattern of movements between patches. Here the location of the cohort is distinguished by the number, which shows when they moved the patch with the number identification number.

We also believe it is beneficial to show why not all features have been fully investigated in this manuscript, and this is to do with the computational speed of the simulations. Figure S4.6 shows how the computation time is altered, in some cases dramatically, by the increase in the number of cohorts, patches or genes. In addition to increased individual simulation run time, there is also an increase in the number of potential parameters with the more complex simulations, which would necessitate more simulations to be conducted to account for this increased parameter space. Together this compounds the total computational time.

**Figure S4.6.**
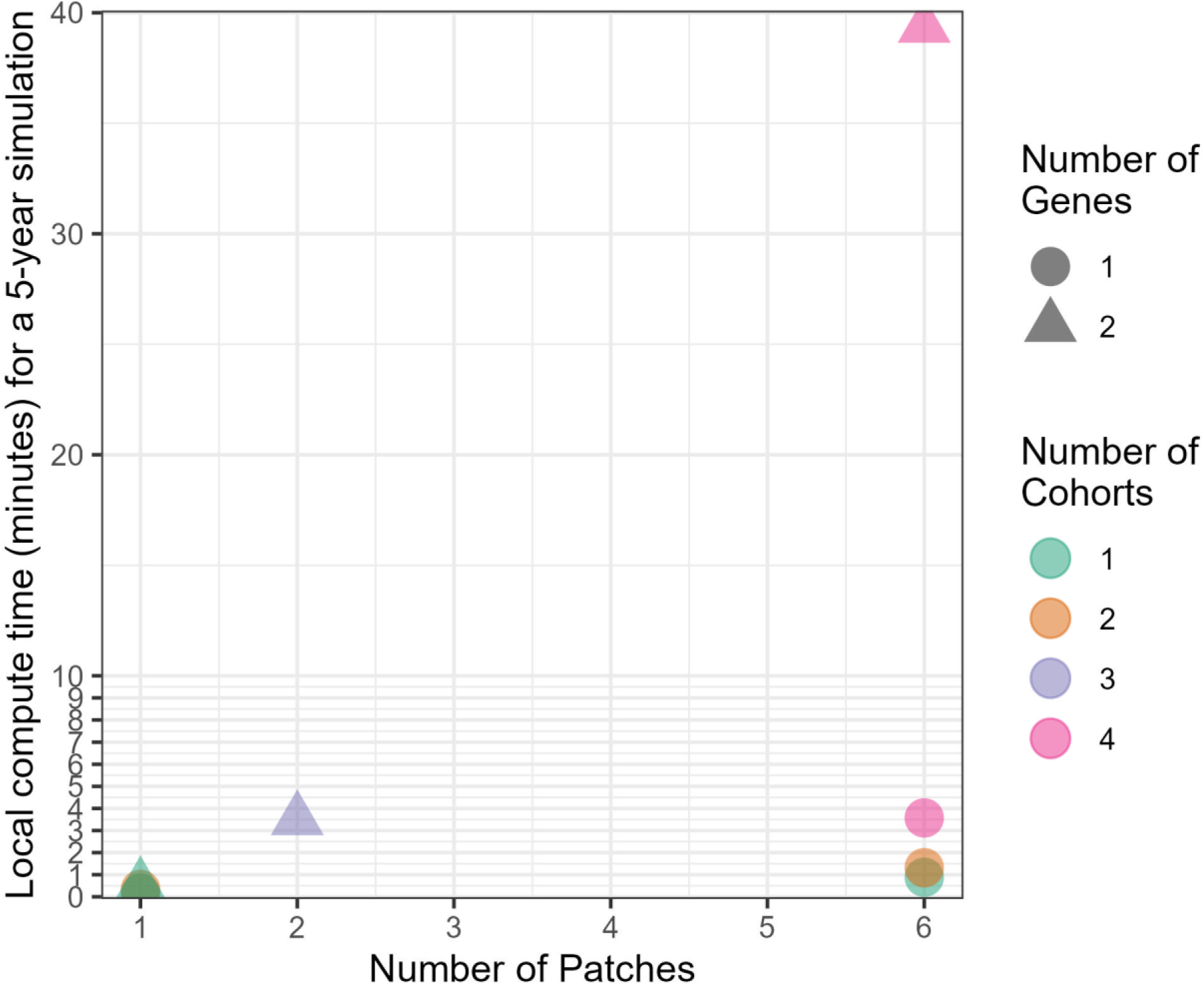
GLOWORM Simulation Framework Computation Time. The local compute time is reported for a single simulation which had a duration of 5-years. It should be noted that this plot is only used to highlight how much the relative computation time is impacted by altering the number of features (cohorts, patches, genes) included in the simulations.

### S5 Model Validation

#### S5.1 Model Validation Overview

The purpose of this model validation is to understand how the GLOWORM model framework performs against real-world empirical data.

As part of this modelling exercise, we attempt to validate our simulation platform against published empirical results from field trials. This helps us to better understand the performance of our model in relation to trial data.

We perform our model validation against three data sets.

1. Stear study (Stear et al., 2006)
2. Sargison study (Sargison et al., 2012)
3. Waller study (Waller and Thomas, 1975)

We would note that validating against trial data does have several challenges. First, we are limited to validating against the reported findings from trials. Trials are expensive and time-consuming to conduct, which unfortunately limits the amount (and variety) of data which is collected. In general, the primary outcome which we are validating against is the reported faecal egg counts from these studies.

Uncertainty in values from published literature:

1. Exact geolocation. This impacts the latitude and longitude for the extraction of the climate data (rainfall and temperature).
2. Pasture contamination.
3. Parasite level in adult ewe populations.
4. Stocking rates and livestock rearing density.
5. Treatment timings (exact); and exact treatment efficacies.

The SCOPE trial (Rose Vineer et al., 2025) hopes to rectify some of the issues with validating and testing models against field trials. Whereby mathematical modellers are directly integrated into the trial team, ensuring trial design is designed around parameters which can be included into a mathematical modelling framework.

#### S5.2 Validation against Stear Study

We attempt to validate our simulations against the results in Figure 4 in (Stear et al., 2006). For simplicity we refer to this study as the Stear study. This data is the mean from two study sites replicated over multiple years. We summarise the conditions in Table S5.1 noting in particular any key time-frames or values with which we use to set up our simulations. The data from both farm locations was used to produce, but we note that the details of Farm 1 (“west central Scotland”) are better detailed, and therefore only attempt to replicate Farm 1. We simulate two cohorts, a ewe cohort and a lamb cohort.

**Table S5.1.** Validation of GLOWORM using the Stear Study.

| **Table S5.1 Validation of GLOWORM using the Stear Study** |  |  |
| --- | --- | --- |
| Study Aspect | Stear Study | GLOWORM Simulations |
| Parasite species | *Teladorsagia circumcincta*, *Cooperia* spp., *Trichostrongylus vitrinus*, *Trichostrongylus axei*, and *Haemonchus contortus*. | We assume a single species infection with the parasite *T. circumcincta*. |
| Geolocation and weather | The location of the farm is described as being in “west central Scotland”. | We assume the study was in an area bounded by the latitude and longitude coordinates of: latitude: min = 55.69, max = 55.81; and longitude min = -4.55, max = -4.39. We sample locations in the latitude and longitude direction at 0.02 intervals, giving a total of 63 sampled geolocations. Extract the weather data for these coordinates using the openmeteo R package (Pisel, 2023) |
| Timeframe | April-October for years 1990 to 1995 | April-October for years 1990 to 1995 |
| Lambing | Lambing occurred during a 3-week period in April-May. | Lambing occurs as a single event where all lambs are born (creation of a new cohort) on 15<sup>th</sup> April. |
| Drug Treatments | A single treatment was given in the months May to October to all lambs. No drug resistance was detected or reported. No mention of drug treatments to parental ewes. | Drug treatments are given to all lambs on the first day of each month (May to October). We assume a single day drug effect, which reduces the within host parasite stages each by 95%.<br>No treatments are given to the parental Ewes. |
| Weaning | Weaning occurred at 3 or 4 months of age (mid-July to mid-August). | Weaning occurred on 1<sup>st</sup> August, whereby the parental ewes (Cohort 1) is removed from the simulation. |
| Movement | Lambs were moved onto new pasture at weaning. | Lambs were moved onto new pasture (patch) at weaning. |
| Pasture contamination | The fields are described as having transmission for the prior 10 years | We implement pre-contaminated pasture (patches), detailed in the simulation R scripts. |
| Immunity | N/A | We assume the immunity of the lambs is reduced to account for their immaturity (the exposure-immunity relationship in the GLOWORM model was developed for adult livestock), with a reduction of 25%.<br>We assume Ewe immunity is reduced to account for the stresses of birth, such that immunity is reduced by 50%. |

The output from the model validation against the Stear study is given in Figure S5.1. It shows good performance for the overall shape, by capturing the overarching peaks, and comparative magnitude of each peak. For example the increasing peak sizes from May to July, and a decline and partial plateauing of the peak sizes between August and September.

**Figure S5.1.**
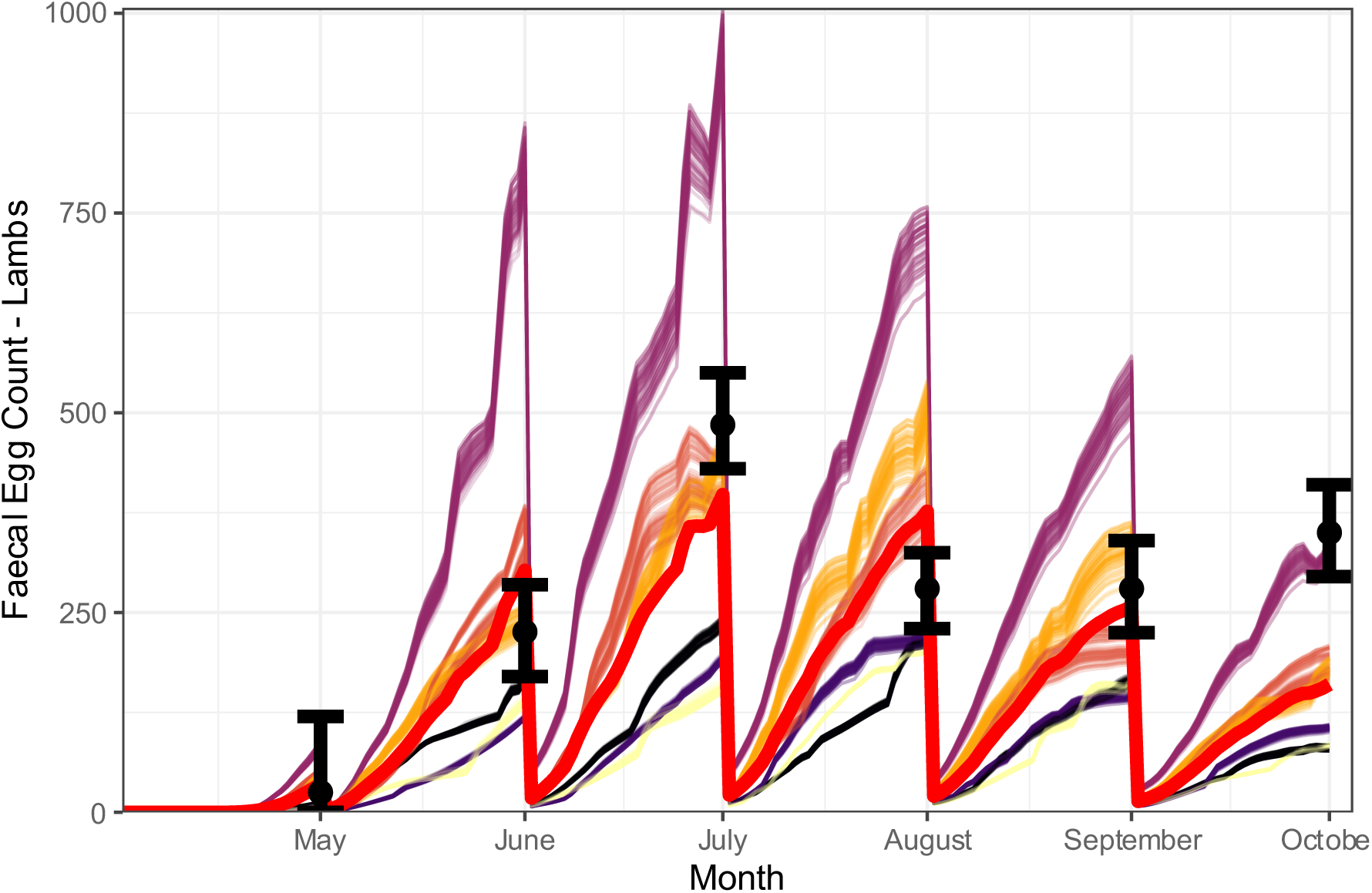
Validation of GLOWORM using the Stear Study. The black point and bars the mean and 95% confidence intervals from the Stear Study (Sargison et al., 2012). The thicker red line is the mean of the GLOWORM validation simulations (n = 378 simulations, 63 locations x 5 years). The thinner coloured lines are the outputs from each individual simulation, with the colour indicating year.

#### S5.3 Validation against Sargison Study

We next attempt to validate the GLOWORM simulation framework against the Sargison Study (Sargison et al., 2012). In brief, this study compared giving moxidectin to parental ewes as either an oral drench (~5-week residual activity) or a long-acting injectable (~16-week residual activity) to control the peri-parturient rise in FEC. With lambs then receiving three rounds of treatment with ivermectin. The oral drench (OD) and long-acting injectable (LA) groups were kept on separate fields, and we therefore conduct simulations for each group separately. The study replicated twice, so there are two OD groups and two LA groups. We simulate two cohorts, a ewe cohort and a lamb cohort. We attempt to replicate the results presented in Figure 2 in Sarginson et al., 2006. We summarise the conditions in Table S5.2 noting in particular any key time-frames or values with which we use to set up our simulations.

**Table S5.2.** Validation of GLOWORM using the Stear Study.

| Table S5.2 Validation of GLOWORM using the Stear Study |  |  |
| --- | --- | --- |
| Study Aspect | Sargison Study | GLOWORM Simulations |
| Parasite species | *T. circumcincta*<br>*Nematodirus battus* | *T. circumcincta* |
| Geolocation and weather | Southeast Scotland | We assume the study was in an area bounded by the latitude and longitude coordinates of: latitude: min = 55.58, max = 55.8; and longitude min = -3.46, max = -3.12. We sample locations in the latitude and longitude direction at 0.02 intervals, giving a total of 216 sampled geolocations. Extract the weather data for these coordinates using the openmeteo R package (Pisel, 2023) |
| Timeframe | The year of the study was 2010. Sample collection began 18th March and ended 26th August. | We extracted weather data from the year 2010.<br>Simulations started March 1<sup>st</sup> to allow for a short burn-in to allow the FECs in the parental ewes to reach the desired level (~600 FEC). Simulations ended 26<sup>th</sup> August, therefore simulations had a total duration of 215 days. |
| Lambing | Lambing occurs between the 29<sup>th</sup> March and 15<sup>th</sup> April. | We take a midpoint of 7<sup>th</sup> April as the lambing day, when all lambs are born (and use this to create a new cohort). |
| Drug Treatments | Parental Ewes: One group received a moxidectin oral drench (~ 5-week residual activity), the other group received moxidectin as a long-acting injectable (~16-week residual activity). The treatments were given between 29<sup>th</sup> March and 15<sup>th</sup> April.<br>All lambs received ivermectin on the days 13<sup>th</sup> May, 10<sup>th</sup> June and 19<sup>th</sup> July. The FECRT for the four groups was 55%, 0%, 47%, and 0%; giving a mean across all four groups of 25.5%. | Ewes (Cohort 1):<br>For the oral drench, treatment on 7<sup>th</sup> April, which has a parasite kill efficacy of 95% for 35 days.<br>For the long-acting injectable, receive treatment on 7<sup>th</sup> April, which has a parasite kill efficacy of 95% for 35 days.<br>All lambs (Cohort 2) receive treatment on days 13<sup>th</sup> May, 10<sup>th</sup> June and 19<sup>th</sup> July. The treatment has a single day parasite killing efficacy of 0.745 (calculated as 1 minus the mean FECRT). |
| Weaning | Weaning occurred 109 days after April 15<sup>th</sup>. | Weaning occurred 109 days after April 15<sup>th</sup>, whereby the parental ewes (Cohort 1) are removed from the simulation. |
| Movement | Prior to turnout the Ewes and lambs were housed. Turnout occurred between 29<sup>th</sup> March to 15<sup>th</sup> April. | We take a midpoint of 7<sup>th</sup> April as the turnout day. From here they move from a patch where transmission does not occur to standard patch. |
| Pasture contamination | Pasture is assumed to be contaminated given previous needs for anthelmintic treatment of the flocks. | We implement pre-contaminated pasture (patches), detailed in the simulation R scripts. |
| Immunity | N/A | We assume the immunity of the lambs (Cohort 2) is reduced to account for their immaturity (the exposure-immunity relationship in the GLOWORM model was developed for adult livestock), with a reduction of 25%.<br>We assume Ewe immunity (Cohort 1) is reduced to account for the stresses of birth, such that immunity is reduced by 50%. |

Figure S5.2 shows the simulated model output when compared to the Sargison Study (Sargison et al., 2012). The model performs well in capturing the overarching behaviour of the system, with the FEC rising shortly after lambing, before gradually declining. The mean values for the long-acting injectable (Figure S5.2, right panel), are marginally lower than for the Oral Drench (left panel), after the effect of the oral drench is no longer active. This difference is very slight.

**Figure S5.1.**
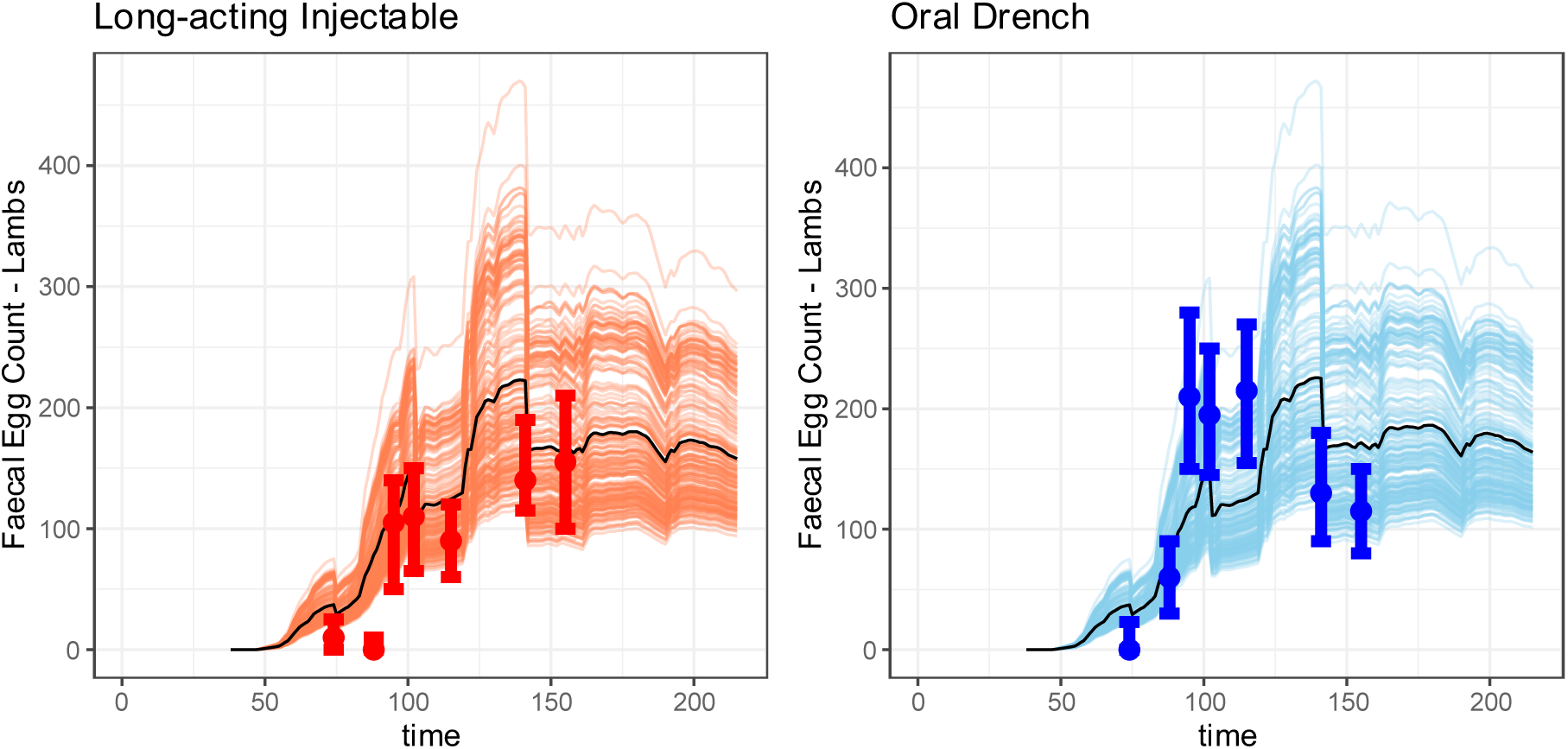
Validation of GLOWORM using the Sargison Study. Left Plot: This is the validation against the long-acting injectable, where the red point and bars are the mean and 95% confidence intervals from the Sargison Study (Sargison et al., 2012), and the light red lines are the output from each individual GLOWORM simulation, with the black line being the mean from those simulations (n=216). Right Plot: This is the validation against the oral drench, where the blue point and bars are the mean and 95% confidence intervals from the Sargison Study (Sargison et al., 2012), and the light blue lines are the output from each individual GLOWORM simulation, with the black line being the mean from those simulations (n=216).

#### S5.4 Validation against Waller Study

We next validate the GLOWORM simulation framework against the Waller Study (Waller and Thomas, 1975), for the parasite *Haemonchus contortus*.

**Table S5.4.**
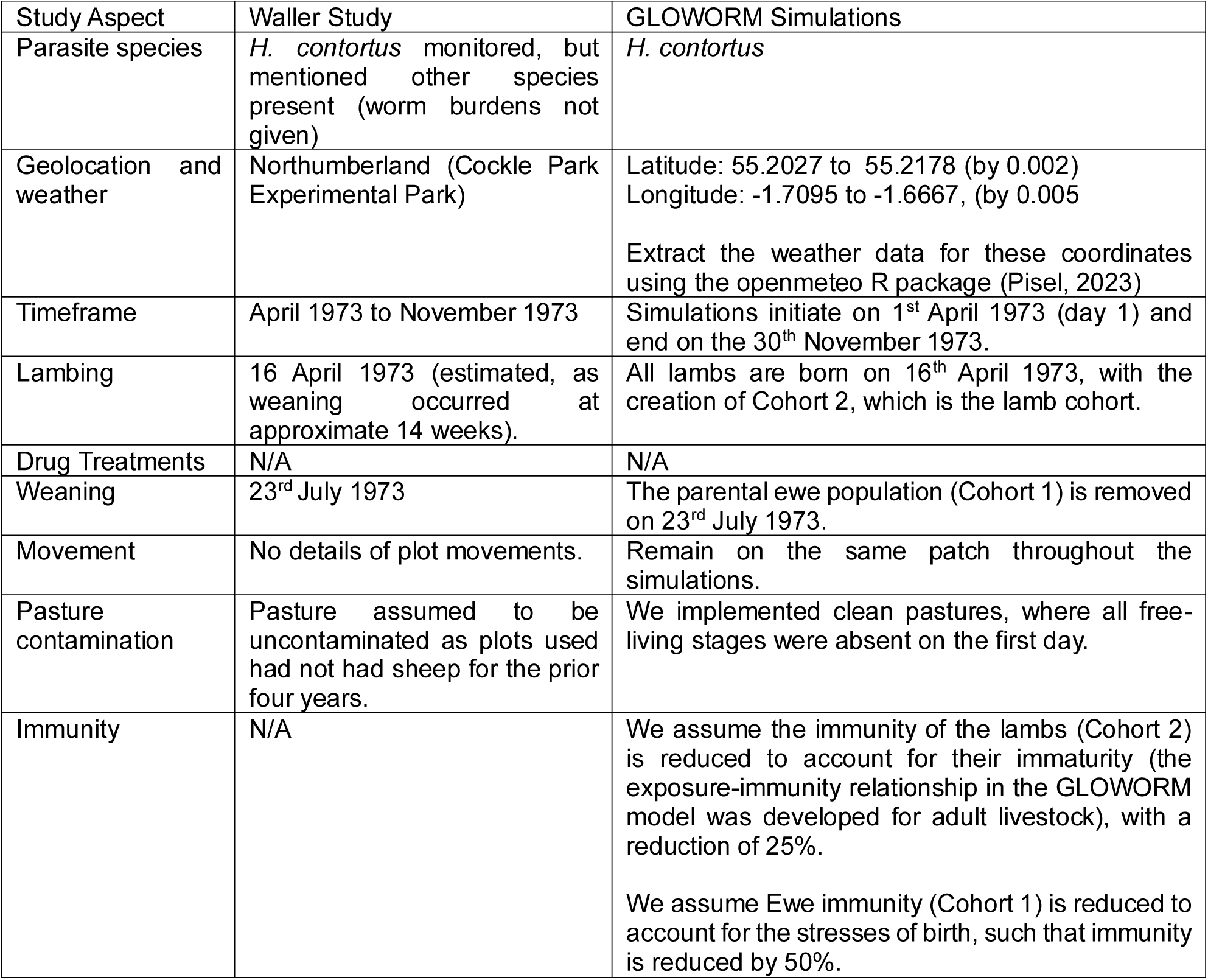
Validation of GLOWORM using the Waller Study.

| Table S5.4 Validation of GLOWORM using the Waller Study |  |  |
| --- | --- | --- |
| Study Aspect | Waller Study | GLOWORM Simulations |
| Parasite species | *H. contortus* monitored, but mentioned other species present (worm burdens not given) | *H. contortus* |
| Geolocation and weather | Northumberland (Cockle Park Experimental Park) | Latitude: 55.2027 to 55.2178 (by 0.002)<br>Longitude: -1.7095 to -1.6667, (by 0.005)<br>Extract the weather data for these coordinates using the openmeteo R package (Pisel, 2023) |
| Timeframe | April 1973 to November 1973 | Simulations initiate on 1<sup>st</sup> April 1973 (day 1) and end on the 30<sup>th</sup> November 1973. |
| Lambing | 16 April 1973 (estimated, as weaning occurred at approximate 14 weeks). | All lambs are born on 16<sup>th</sup> April 1973, with the creation of Cohort 2, which is the lamb cohort. |
| Drug Treatments | N/A | N/A |
| Weaning | 23<sup>rd</sup> July 1973 | The parental ewe population (Cohort 1) is removed on 23<sup>rd</sup> July 1973. |
| Movement | No details of plot movements. | Remain on the same patch throughout the simulations. |
| Pasture contamination | Pasture assumed to be uncontaminated as plots used had not had sheep for the prior four years. | We implemented clean pastures, where all free-living stages were absent on the first day. |
| Immunity | N/A | We assume the immunity of the lambs (Cohort 2) is reduced to account for their immaturity (the exposure-immunity relationship in the GLOWORM model was developed for adult livestock), with a reduction of 25%.<br>We assume Ewe immunity (Cohort 1) is reduced to account for the stresses of birth, such that immunity is reduced by 50%. |

**Figure S5.4.**
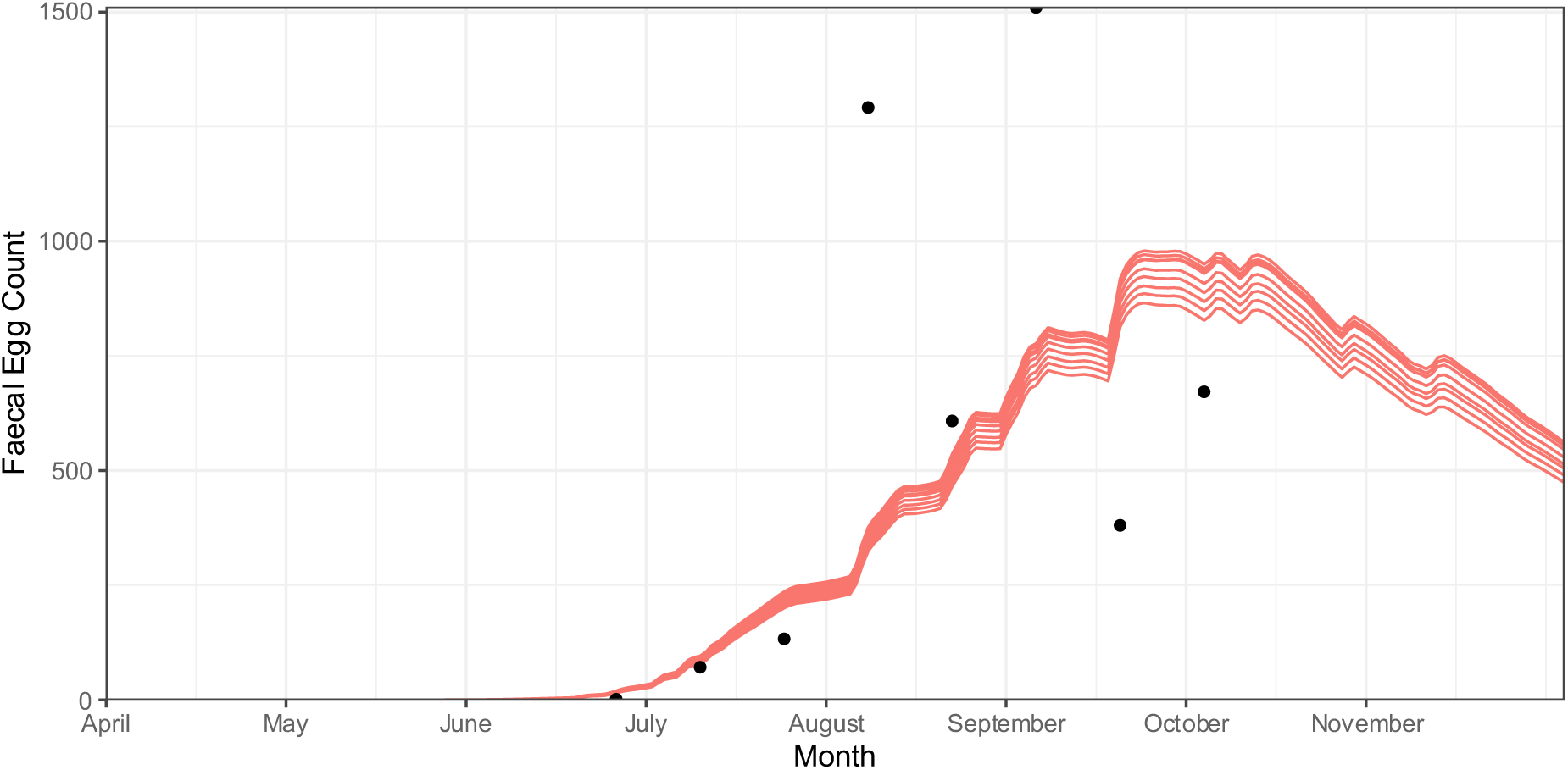
Validation of GLOWORM using the Waller Study. Lines are the simulated faecal egg counts of the lambs. Points are the faecal egg counts from the Waller Study (estimated using plotdigitiser).

##### Comments on validation

The validation provides a good fit during the early phase of the infection. Bisects the later points; but as no confidence intervals were provided in the Waller Study. Given the variability of these points over a short timeframe, it could be assumed the mean values lies within these ranges and therefore the model still matches the overarching transmission dynamics of *H. contortus*.

## Notes

### Competing Interest Statement

The authors have declared no competing interest.

